# GwEEP - A comprehensive approach for genome-wide efficiency profiling of DNA modifying enzymes

**DOI:** 10.1101/2020.08.06.236307

**Authors:** Charalampos Kyriakopoulos, Karl Nordström, Paula Linh Kramer, Judith Gottfreund, Abdulrahman Salhab, Julia Arand, Fabian Müller, Ferdinand von Meyenn, Gabriella Ficz, Wolf Reik, Verena Wolf, Jörn Walter, Pascal Giehr

## Abstract

A precise understanding of DNA methylation dynamics on a genome wide scale is of great importance for the comprehensive investigation of a variety of biological processes such as reprogramming of somatic cells to iPSCs, cell differentiation and also cancer development. To date, a complex integration of multiple and distinct genome wide data sets is required to derive the global activity of DNA modifying enzymes. We present GwEEP - *Genome-wide Epigenetic Efficiency Profiling* as a versatile approach to infer dynamic efficiency changes of DNA modifying enzymes at base pair resolution on a genome wide scale. GwEEP relies on genome wide *oxidative Hairpin Bisulfite sequencing* (HPoxBS) data sets, which are translated by a sophisticated hidden Markov model into quantitative enzyme efficiencies with reported confidence around the estimates. GwEEP in its present form predicts *de novo* and maintenance methylation efficiencies of Dnmts, as well as the hydroxylation efficiency of Tets but its purposefully flexible design allows to capture further oxidation processes such as formylation and carboxylation given available data in the future. Applied to a well characterized ES cell model, GwEEP precisely predicts the complex epigenetic changes following a Serum-to-2i shift i.e., (i) instant reduction in maintenance efficiency (ii) gradually decreasing de novo methylation efficiency and (iii) increasing Tet efficiencies. In addition, a complementary analysis of Tet triple knock-out ES cells confirms the previous hypothesized mutual interference of Dnmts and Tets. GwEEP is applicable to a wide range of biological samples including cell lines, but also tissues and primary cell types.

**MOTIVATION:** Dynamic changes of DNA methylation patterns are a common phenomenon in epigenetics. Although a stable DNA methylation profile is essential for cell identity, developmental processes require the rearrangement of 5-methylcytosine in the genome. Stable methylation patterns are the result of balanced Dnmts and Tets activities, while methylome transformation results from a coordinated change in Dnmt and Tet efficiencies. Such transformations occur on a global scale, for example during the reprogramming of maternal and paternal methylation patterns and the establishment of novel cell type specific methylomes during embryonic development *in vivo*, but also *in vitro* during (re)programming of induced pluripotent stem cells, as well as somatic cells. In addition, local (de)methylation events are key for gene regulation during cell differentiation. A detailed understanding of Dnmt and Tet cooperation is essential for understanding natural epigenetic adaptation as well as optimization of *in vitro* (re)programming protocols. For this purpose, we developed a pipeline for quantitative and precise estimation of Dnmt and Tet activity. Using only double strand methylation information, GwEEP infers accurate maintenance and *de novo* methylation efficiency of Dnmts, as well as hydroxylation efficiency of Tets at single base resolution. Thus, we believe GwEEP provides a powerful tool for the investigation of methylome rearrangements in various systems.

## INTRODUCTION

Genetic information encoded in the DNA is regulated by epigenetic mechanisms, such as DNA methylation [1, 2, 3, 4]. In mammals, methylation of DNA is restricted to cytosine and is almost exclusively found in a palindromic CpG dinucleotide context [5, 6, 7]. Generation of 5-methylcytosine (5mC) is catalyzed by the DNA methyltransferases Dnmt1, Dnmt3a and Dnmt3b. These enzymes catalyze the transfer of a methyl group from s-adenosyl methionine to the fifth carbon atom of cytosine. Dnmt1 is responsible for maintaining existing methylation patterns after replication. Via interaction with Uhrf1 and PCNA, Dnmt1 is tightly associated with the replication machinery [8, 9]. Furthermore, the cooperation with Uhrf1 modulates Dnmt1 to be receptive for hemimethylated DNA generated after replication [10, 11]. Thus, the protein complex post-replicatively copies the methylation pattern from the inherited to the newly synthesized DNA strand [12, 13]. Dnmt3a and Dnmt3b methylate DNA independently of its methylation status (hemimethylated or unmethylated) and are mainly responsible for the establishment of new methylation patterns during development [14, 15]. However, several studies indicate that the strict separation of Dnmt1 and Dnmt3a/b activity is not coherent and that under certain conditions, these enzymes exhibit overlapping functions [16, 17, 18]. Once established, 5mC can be further processed by a family of di-oxigenases, the ten-eleven translocation enzymes Tet1, Tet2 and Tet3 [19, 20, 21]. These Fe(II) and oxoglutarate-dependent enzymes consecutively oxidize 5mC to 5-hydroxymethyl cytosine (5hmC), 5-formyl cytosine (5fC) and ultimately to 5-carboxy cytosine (5caC) [22, 23]. 5hmC is the most abundant oxidative variant and can be found in numerous cell types [24, 25, 26]. Each oxidation step changes the chemical properties of the base and with it its biological function [27, 28, 29]. Several mechanisms have been proposed in which oxidative cytosine derivatives (oxC) serve as an intermediate during the course of active or passive demethylation [30, 31, 32, 33, 34]. Such removal of 5mC occurs locally during cell differentiation, but also on a genome-wide scale in the zygote, as well as during the maturation of primordial germ cells (PGCs) [35, 36, 37]. Global loss of 5mC has been observed in cultivated mouse embryonic stem cells (ES cells) during their transition from Serum to 2i medium. Under classical Serum/LIF conditions, ES cells exhibit DNA hypermethylation, whereas upon transition to GSK3 and Erk1/2 inhibitors (2i) containing medium, the cells experience a gradual genome-wide loss of 5mC [38, 39, 40]. Even though several studies examined the influence of Tets and oxCs [41, 42, 43, 44] within the genome, the precise contribution of Tets and oxCs towards maintaining or change existing cell type specific methylomes remains elusive. A contextual understanding of the local and spatial connections between Dnmts and Tets during processes of development, cell division and differentiation is of great importance, as they build the basis for a rational development of novel epigenetic cancer therapies and controlled reprogramming approaches in regenerative stem cell medicine.

To address the complex interplay between Dnmts and Tets we developed a comprehensive experimental and computational pipeline - GwEEP (**G**enome **w**ide **E**nzyme **E**fficiency **P**rofiling), which determines the efficiencies of DNA modifying enzymes across the genome. GwEEP comprises genome wide hairpin (oxidative) bisulfite sequencing - HP(ox)BS, a pipeline for extracting the double strand methylation pattern from HPBS, as well as a sophisticated hidden markov model to infer Dnmt’s *de novo* and maintenance methylation, as well as Tet’s hydroxylation efficiencies. Here we provide a detailed protocol for the generation of hairpin (oxidative) bisulfite libraries and furthermore, a detailed description data processing and HMM application. Lastly, we provide an application example where we applied GwEEP to a widely used model system, following the adaptation of ES cells during their transition from Serum/LIF to 2i based media.

## MATERIAL

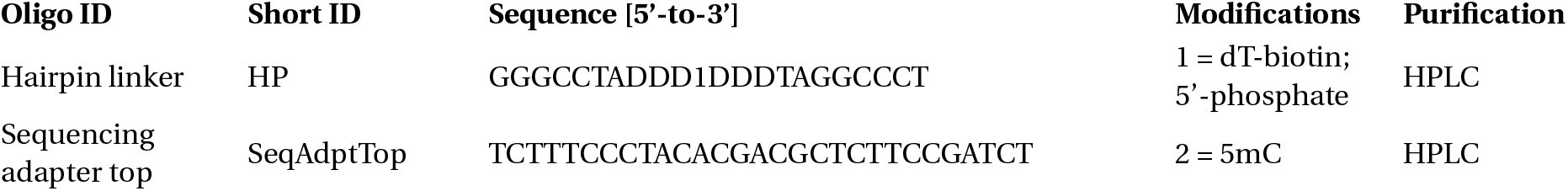
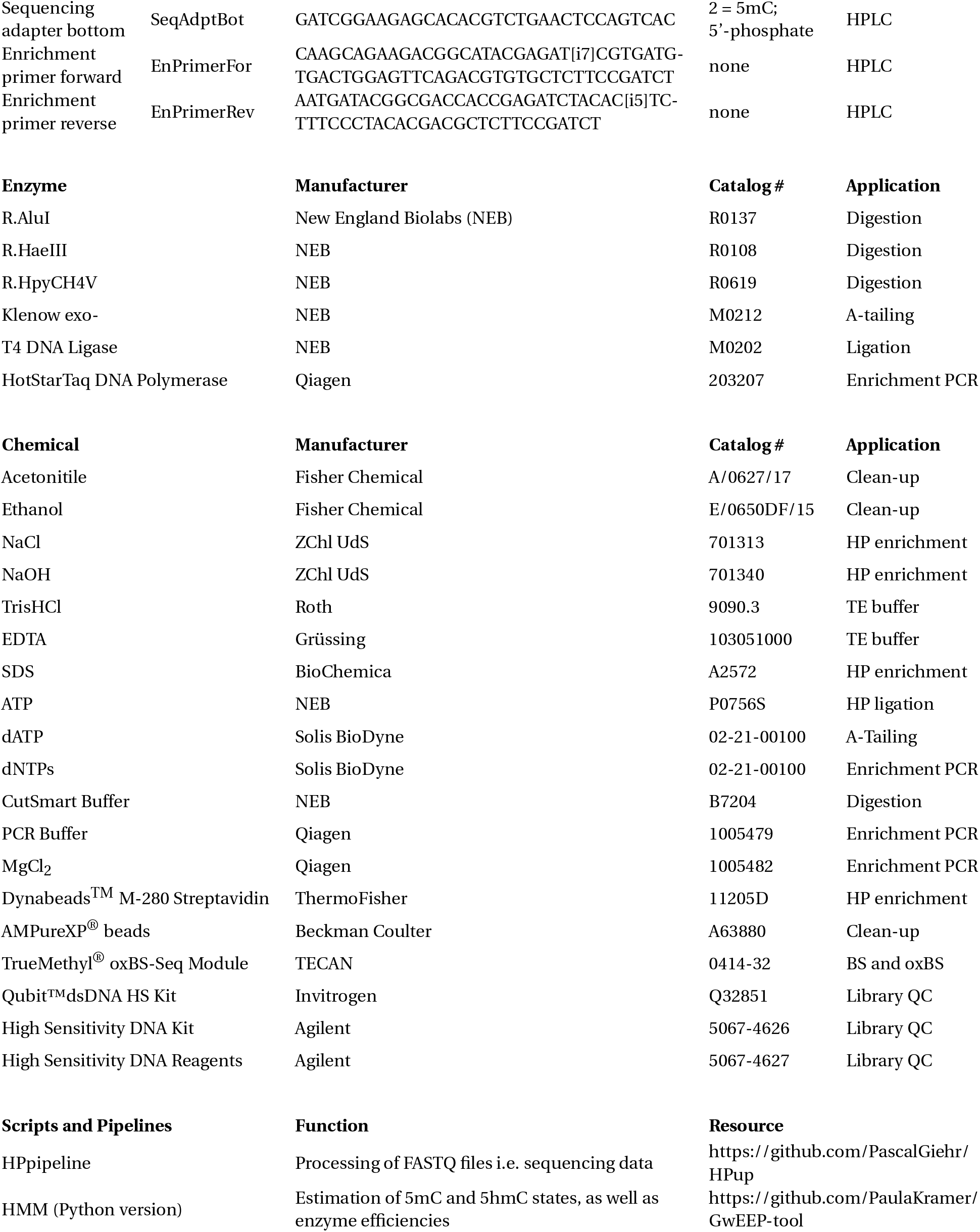

## METHODS

Our pipeline consists of three major parts: (i) the construction of an oxidative hairpin bisulfite library (RRHPoxBS), (ii) a computational pipeline (HPup) which extracts the double strand DNA methylation values from Illumina sequencing data and (iii) a hidden Markov model (GwEEP-tool) that calculates conversion error corrected 5mC and 5hmC distributions and furthermore infers the enyzmatic efficiencies of Dnmts (maintenance and *de novo* methylation efficiency), as well as Tets (hydroxylation efficiency)

### RRHPoxBS - Reduced Representation Hairpin oxidative Bisulfite Sequencing

#### Digestion of genomic DNA

For GwEEP we use endo nucleases (RE) in order to enrich for selected genomic regions. This increases the sensitivity of the assay (as little as 18 ng of DNA can be used - SI Sec 14) and, at the same time, reduces the sequencing costs considerably. The RE must not be sensitive against 5mC and 5hmC, in order to avoid a bias of the methylation analysis by blocked restriction. Ideally, the recognition/restriction site of the RE should not contain any CpG dyad. In the present protocol, we used a combination of three different REs i.e., R.AluI, R.HaeIII and R.HpyCH4V to obtain reads from regulatory (HaeIII), as well as inter and intra genomic regions (AluI and HpyCH4V). For the customization of GwEEP we recommend cuRRBS, which determines the best suited RE(s) based on a list of *regions of interest* [45].

Unless the amount of DNA is limited, we strongly advice a proper quality control using *Agarose*-*gel*-*elec-trophoresis* and *Qubit Fluorometer* based quantification. The best results are obtained by using 300 - 400 ng high quality DNA per used RE. However, as little as 2 ng per enzyme (6 ng in total) can be used. For each RE, prepare the reaction outlined in Table 2 and incubate for at least 3h at the temperature optimum of the RE (here 37°C), followed by a 20 min heat inactivation at 65°C or 80°C.

**Table 2:** Digest of genomic DNA

| Component | Concentration | Volume [μl] |
| --- | --- | --- |
| Genomic DNA | 2 - 400 ng | X |
| Endo Nuclease<br>(R.AluI, R.HaeII or R.HpyCH4V) | 10 U | Y |
| CutSmart Buffer | 10x | 2.0 |
| ddH<sub>2</sub>O | – | ad 20.0 |
| Total Volume |  | 20.0 |

**Table 3:**
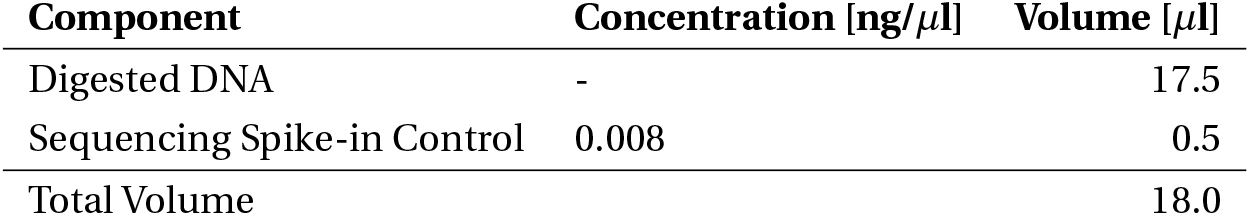
Addition of Spike-in Control

**NOTE: Incubation over night might increase the sensitivity of the assay, especially when using low DNA amounts or dealing with sample impurities. However, this is only recommended for RE that do not exhibit any star activity**.

After inactivation the reactions are combined and subjected to a purification step using magnetic beads i.e., SPRI^®^ or AmpureXP^®^ beads. Use a sample:bead ratio of 1:2 (60*µ*l:120*µ*l) for clean-up, wash twice with 200 *µ*l freshly prepared 80% ethanol (EtOH) and elute the DNA in 17.5 *µ*l of 1x CutSmart buffer.

At this stage we strongly recommend to add a suitable spike-in in order to calculate conversion rates of bisulfite and oxidative bisulfite reactions independently from the biological sample. We apply 4 pg of Sequencing Spike-in Control (CEGX) per sample (SI Sec. 6).

#### End-repair and A-tailing

In the present study, we applied REs which generate blunt-end DNA fragments and therefore require a subsequent A-tailing. For this, we rely on the polymerase I large (Klenow) fragment, which lacks both 5^*′*^*- >* 3^*′*^, as well as 3^*′*^*- >* 5^*′*^ exonuclease activity (Klenow exo-). The reaction given in Table 4 is incubated for 30 min at 37°C and inactivated for 20 min at 75°C:

**Table 4:** A-tailing reaction

| Component | Concentration | Volume [μl] |
| --- | --- | --- |
| Digested DNA | – | 18.0 |
| Klenow exo- | 5 U/μl | 1.0 |
| dATP in 2x CutSmart | 1 mM | 1.0 |
| Total Volume |  | 20.0 |

**NOTE: In case REs producing sticky-ends are used, the above reaction is not sufficient for library preparation. Instead, please rely on an end-repair and A-tailing workflow**.

The A-tailing prevents self-ligation of DNA fragments and facilitates the ligation of hairpin linker (HP) and sequencing adapter (SA). Purification after heat-inactivation of Klenow exois not required. Instead, proceed immediately to section *Ligation of hairpin-linker and sequencing adapter*.

#### Ligation of hairpin-linker and sequencing adapter

Prior to the ligation reaction, SA and HP have to be transferred into a double stranded state. First, solve each oligo nucleotide in 1xTE for a final concentration of 100*µ*M. Next, join 10 *µ*l SeqAdptTop and 10 *µ*l SeqAdptBot in a 0.2 ml reaction tube and fill 20*µ*l of HP in a second 0.2 ml reaction tube. Place both tubes into a thermocylcer and incubate for 5 min at 95°C followed by a slow cool-down of about 0.3°C/sec. This will facilitate the annealing of SeqAdptTop and SeqAdptBot, as well as the proper folding of the HP.

Figure 2 displays the sequence and structure of SA and HP before and after annealing/folding. The HP comprises four distinct features, (i) a 3’ T overhang complementary to the A-overhang of the DNA molecules, (ii) unmethylated cytosine within the sequence, which permits the calculation of C-to-T conversion rate after sequencing, (iii) a unique molecular identifier (UMI), which allows to identify individual original ligation events and the remove of clonal PCR amplificates and (iv) a biotinylated T, for enrichment of HP containing DNA molecules after ligation. The ligation of SA and HP are catalysed by the T4 DNA ligase following the reaction outlined in Table 5.

**Table 5:**
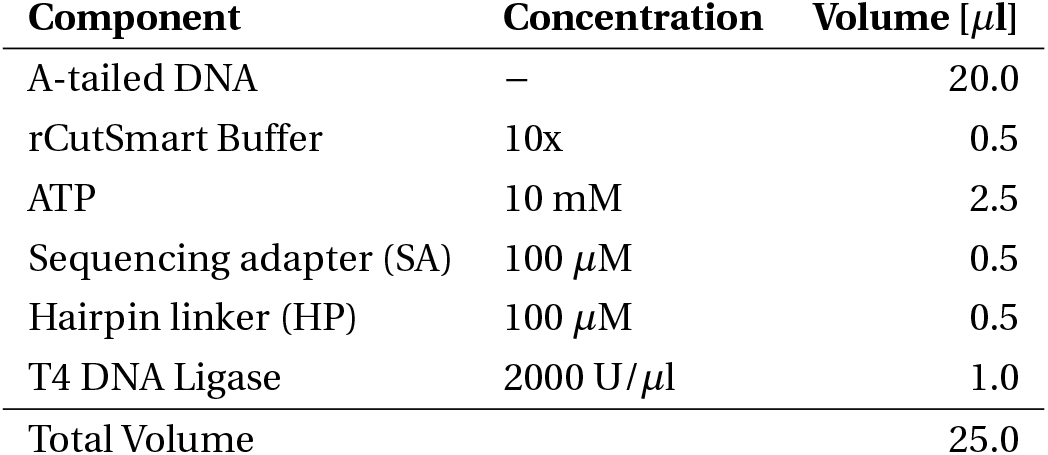
Ligation reaction

**Figure 1:**
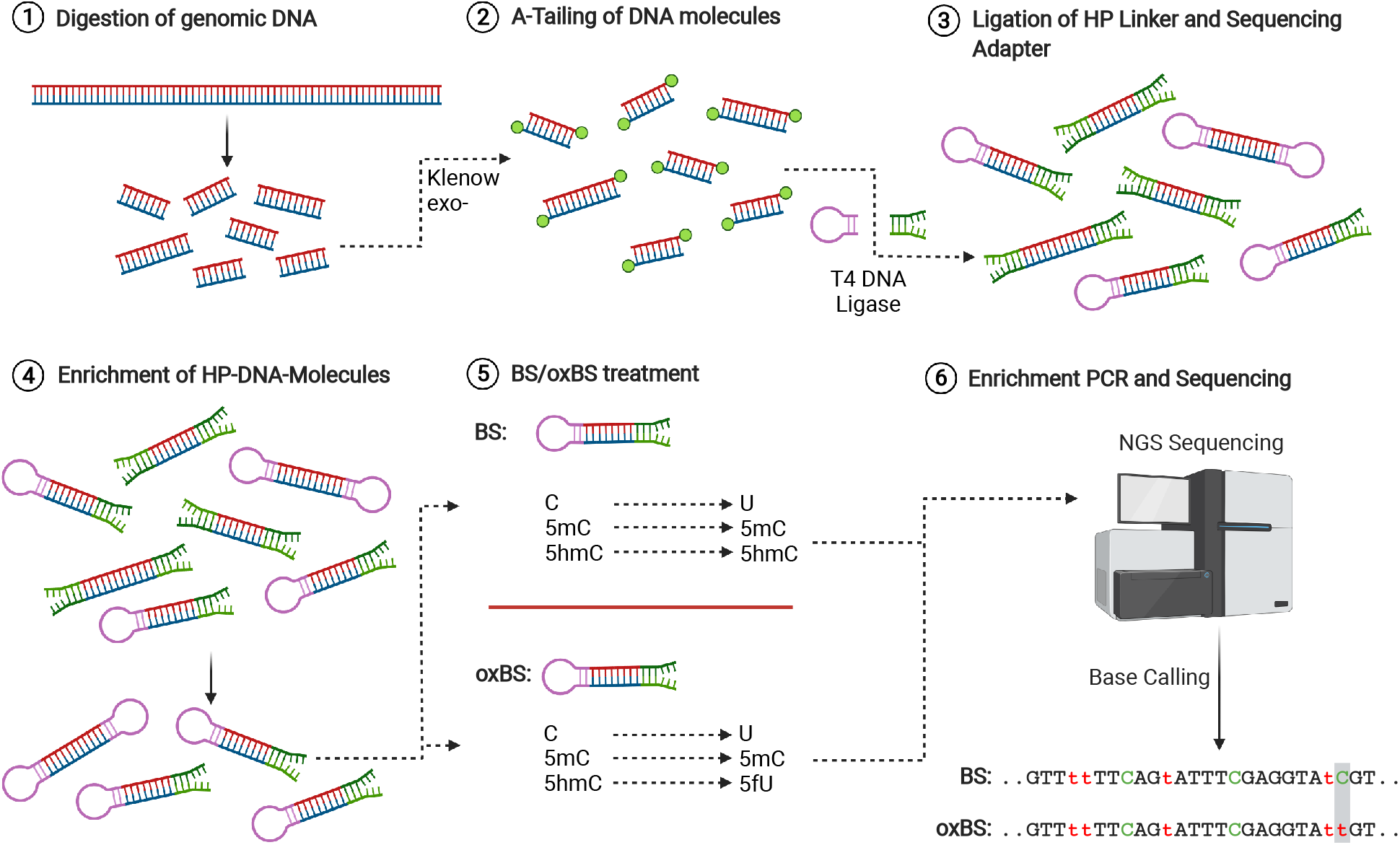
Laboratory Pipeline Overview - (1) genomic DNA is digested by endo nucleases, followed by (2) Klenow exocatalyzed A-tailing. (3) A-Tailed DNA molecules are subjected to sequencing adapter and hairpin linker ligation and (4) subsequent enrichment of hairpin-ligated molecules. (5) half of the library is used for Bs, the other half for oxBs treatment. (6) After amplification and indexing using PCR, the libraries are sequenced on an Illumina platform with minimum 100 bp in a paired-end mode. Created with BioRender.com

**Figure 2:**
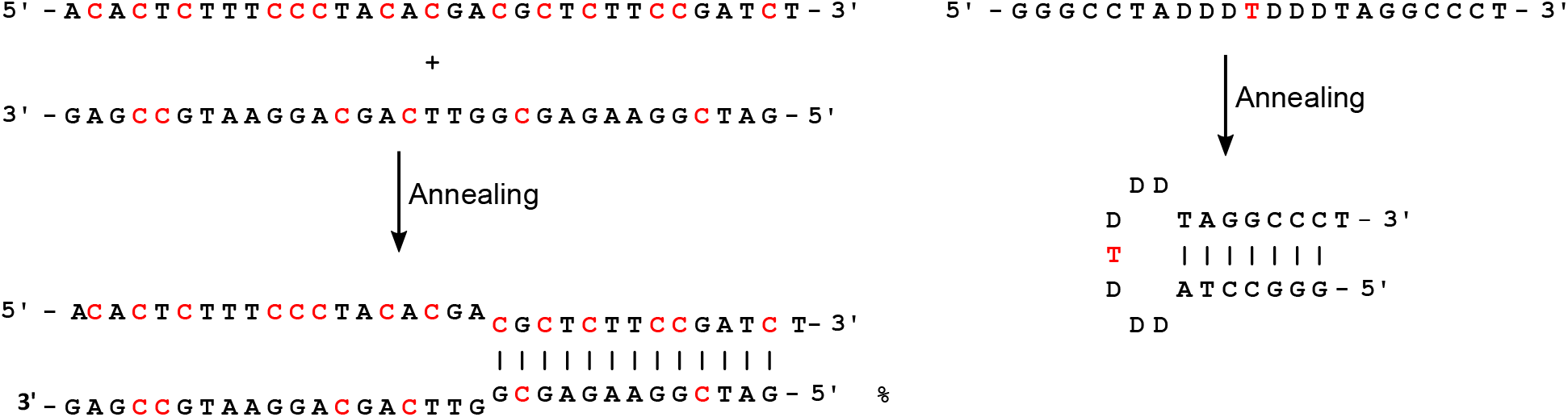
Annealing of Sequencing Adapter and Hairpin Linker - **Left:** Annealing of SeqAdptTop and SeqAdptBot, red Cs indicate methylated Cs; **Right:** Folding of the Hairpin linker, red T indicates biotinylated dT.

**Figure 3:**
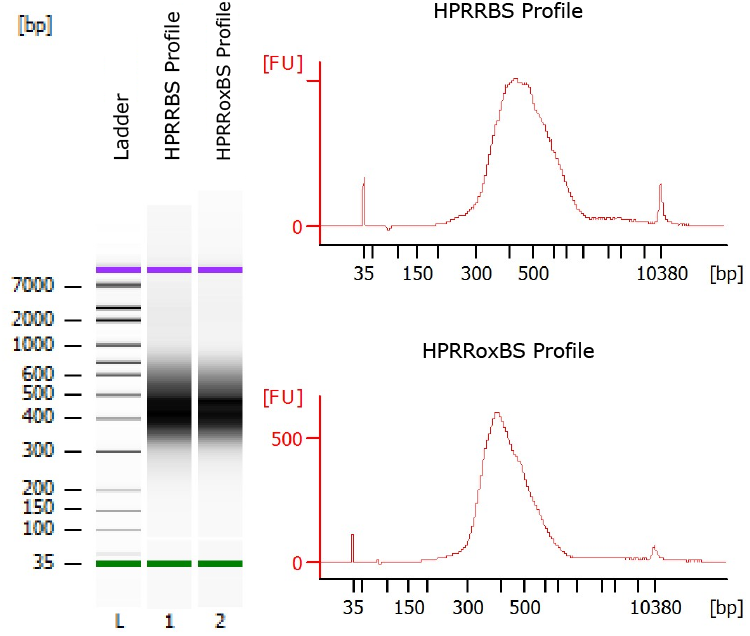
Bio-Analyzer Profiles - Typical Bio Analyzer profile of HPRRBS an HPRRoxBS libraries after enrichment PCR.

**Figure 4:**
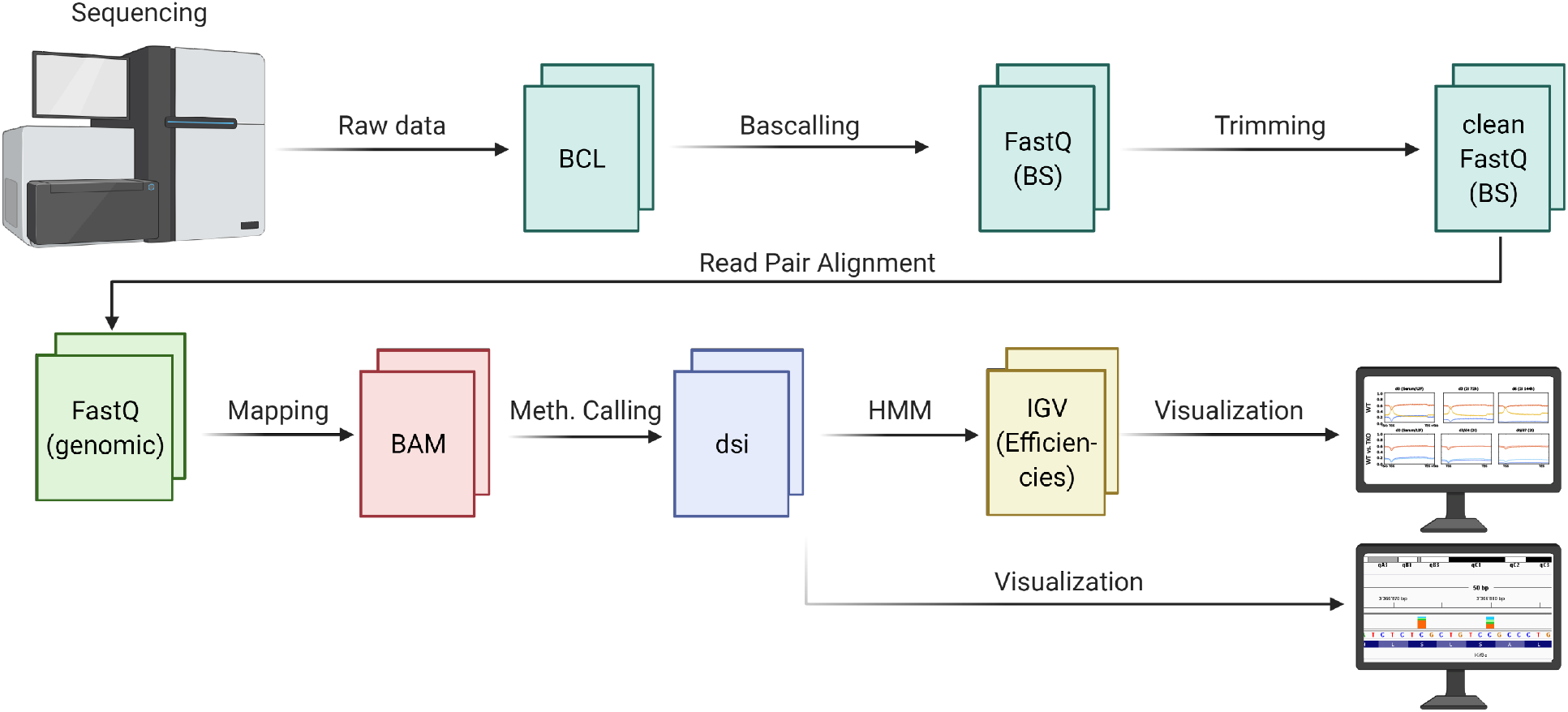
Data Processing and Analysis Pipeline - Illumina raw data is processed into base calls (FASTQ) and trimmed for adapter and hairpin linker sequences. Bisulfite reads from the same molecules are paired to restore the genomic sequence for efficient mapping. Subsequently, the doublestrand information is annotated and stored in - Created with BioRender.com

The reaction is incubated at 16°C for 16 h and inactivated at 65°C for 10 min. The ligation of SA and HP is a stochastic event, meaning that three distinct types of DNA molecules will be generated: (i) molecules with SA on both ends, (ii) molecules with HP on both ends and (iii) molecules with SA on one side and HP on the other side.

**This is a safe stopping point. The DNA can be stored for 24 h at 4°C or long term (days to weeks) at -20°C or -80°C. Please avoid repeated thawing and freezing, as this can lead to strand breaks and thus significantly reduces the number of usable DNA molecules**.

NOTE: Including other modification such as 5mC, 5hmC, 5fC and 5caC, in the HP linker sequence, will allow to determine the conversion of non-canonical cytosine forms during BS and oxBS treatment.

#### Enrichment of Hairpin-DNA-adapter molecules

In order to deplete the unwanted non-HP molecules (SA on both sides) the library is subjected to a purification step using streptavidin coated beads. Start with an AMPure beads clean-up to remove proteins and excessive HPs that would likely saturate the streptavidin beads. Use a sample:bead ratio of 1:2 (25*µ*l:50*µ*l), wash twice with freshly prepared 80% EtOH and elute in 50*µ*l ddH_2_O.

1. Per library i.e., sample, transfer 10*µ*l of Dynabeads™ M-280 Streptavidin into a 1.5 ml reaction tube. Place the tube onto a magnetic stand and carefully remove the supernatant without disturbing the beads.
2. Add 1 ml 1xBW buffer (5 mM TrisHCl, 0.5 mM EDTA, 1 M NaCl), vortex thoroughly, place the mixture back onto the magnetic stand and wait until the solution is clear. Carefully remove and discard the supernatant.
3. Repeat step (2) for a total of 2 wash steps. At the end of wash step three, solve the beads in 50*µ*l 2xBW buffer per library.
4. Transfer 50 *µ*l of beads to each library, mix briefly by flicking the tube and incubate for 20 min at room-temperature while rotating.
5. Collect all liquid at the bottom of the tube though brief centrifugation and place the tube onto the magnetic stand. Wait until the solution is clear and carefully remove and discard the supernatant.
6. Add 200 *µ*l 0.1N NaOH and mix by vortexing. Spin down all liquid, place the tube back onto the magnetic stand and wait until the solution is clear. Remove and discard the supernatant.
7. Repeat step (6) for a total of 2 wash steps.
8. Add 200 *µ*l 0.5xTE buffer to the beads, mix by vortexing and place the tube back onto the magnetic stand. Wait until the solution is clear. Remove and discard the supernatant.
9. Repeat step (8) for a total of 2 wash steps. At the end of the second wash step, remove TE buffer completely, but do not let the beads dry fully.
10. Resuspend the beads in 50 *µ*l 1xTE with 1% SDS.
11. Lock the tube and incubate the mixture for 5 min at 100°C in order to dissolve the the biotin-streptavidin-interaction. Place on ice for 2 min.
12. Briefly spin down and place the tube onto the magnetic stand. Wait until the solution is clear and transfer the supernatant containing the DNA into a new 1.5 ml DNA low-bind reaction tube.

**The use of 1.5 ml DNA-low-bind reactions tubes might increase the final yield of the library. This is especially relevant when working with low input samples**.

**This is a safe stopping point. The DNA can be stored for 24 h at 4°C or long term (days to weeks) at -20°C or -80°C. Please avoid repeated thawing and freezing, as this can lead to strand breaks and thus significantly reduces the number of usable DNA molecules**.

#### BS and oxBS treatment

For bisulfite and oxidative bisulfite treatment, we rely on the TrueMethyl®oxBS Module from TECAN. Here, we will provide a short summary of the individual steps of the protocol.

##### DNA Purification

Bring the Magnetic Bead Solution 1 (TrueMethyl Kit) and the Binding Buffer 1 (TrueMethyl Kit) to room temperature. Mix bead and buffer according to manufacturer instruction. To 50 *µ*l DNA (in 1x TE and 1% SDS), add 100 *µ*l bead-buffer solution and mix well by pipetting. Incubate at room temperature for 20 min. Transfer the solution onto the magnet and wait until the solution is clear (5 min). Carefully remove and discard the supernatant and wash 3x with 200 *µ*l freshly prepared 80% acetonitrile. Let the beads dry for 5 min until all acetonitrile is evaporated. Remove the reaction tube from the magnet and solve the beads in 18 *µ*l denaturation solution (TruMethyl Kit). Elute and denature the DNA at 37°C for 5 min. Put the tube back onto the magnet and wait until the solution is clear.

##### DNA oxidation

Transfer 9 *µ*l into two 0.2 ml reaction tubes, respectively. To the tube intended for BS, add 1 *µ*l ultrapure water (TrueMethyl Kit) in the other, meant for oxBS, add 1 *µ*l oxidant solution. Mix both reactions by vortexing and immediately incubate at 40°C for 10 min. Centrifuge at 14000 x g for 10 min. A black precipitate will form at the bottom of the tube of the oxBS reactions. The supernatant should remain orange. Any other color (yellow, brown, transparent) indicates impurity of the sample and in our experiments indicates failing of the 5hmC oxidation. Transfer the supernatant from oxBS samples into a new 0.2 ml reaction tube.

##### Bisulfite treatment

Add 700 *µ*l of the Bisulfite Dilutent (TrueMethyl Kit) to one aliquot of Bisulfite Reagent (TrueMethyl Kit). Incubate for 15 min at 60°C while shaking. Spin down briefly and add 30 *µ*l of the bisulfite solution to BS and oxBS sample, respectively. Incubate the reaction according to the temperature profile outlined in table 6:

**Table 6:** Bisulfite Conversion Temperature Profile

| Step Number | Incubation Step | Temperature | Time |
| --- | --- | --- | --- |
| 1 | Denaturation | 95°C | 05:00 min |
| 2 | Sulfonation | 60°C | 20:00 min |
| 3 | Denaturation | 95°C | 05:00 min |
| 4 | Sulfonation | 60°C | 40:00 min |
| 5 | Denaturation | 95°C | 05:00 min |
| 6 | Sulfonation | 95°C | 45:00 min |
| 7 | Hold | 20°C | ≤ 16:00:00 h |

Prepare Magnetic Bead Solution 2 by mixing 2.4 *µ*l Magnetic Beads Solution (TrueMethyl Kit) with 200 *µ*l Binding Buffer 2 (TrueMEthyl Kit). Mix well by vortexing. Add 160 *µ*l bead solution to 40 *µ*l BS and 40 *µ*l oxBS reaction, respectively. Mix well by pipetting and incubate for 5 min. Place the reaction onto an magnetic stand, wait until the solution is clear and discard the supernatant. Wash the beads with 200 *µ*l freshly prepared 70% ethanol and resuspend the beads in 200 *µ*l Desulfonation Buffer(TrueMethyl Kit). Incubate for 5 min, remove the supernatant and wash the beads twice with 200 *µ*l 70% ethanol. Let the beads dry completely for 15 min and elute the DNA in 12.5 *µ*l Elution Buffer (TrueMethyl Kit).

#### Enrichment-PCR and sequencing

The enrichment PCR amplfies the library molecules and at the same time introduces the remaining part of the sequencing adapter i.e., indices (i5 and i7), as well as the sequences required for flow cell binding (P5 and P7). Table 7 and Table 8 provide the reaction conditions and the temperature profile of the enrichment PCR, respectively.

**Table 7:** Enrichment PCR

| Component | Concentration | Volume [μl] |
| --- | --- | --- |
| BS or oxBS DNA | – | 10.0 |
| HotStarTaq Buffer | 10x | 5.0 |
| MgCl<sub>2</sub> | 25 mM | 2.0 |
| dNTPs | 10 mM | 4.0 |
| EnPrimerFor | 10 μM | 0.8 |
| EnPrimerRev | 10 μM | 0.8 |
| HoStarTaq DNA Polymerase | 5 U/μl | 0.7 |
| ddH<sub>2</sub>O | – | 26.7 |
| Total Volume |  | 50.0 |

**Table 8:** Enrichment PCR Temperature Profile

| PCR Step | Temperature | Time | Number of Cycles |
| --- | --- | --- | --- |
| Initial Denaturation | 95°C | 5:00 min | 1 |
| Denaturation | 94°C | 0:30 min |  |
| Annealing | 58°C | 1:00 min | 7-18 |
| Elongation | 72°C | 1:00 min |  |
| Final Elongation | 72°C | 7:00 min | 1 |
| Hold | 4°C | ∞ | 1 |

After amplification, libraries are purified using AMPureXP®beads with a sample:bead ratio of 1:1 (50*µ*l:50*µ*l), wash twice with freshly prepared 80% EtOH and elute in 10*µ*l ddH_2_O or 0.1xTE. QC is performed by quantification of 2 *µ*l of the library using Qubit HS Fluorometer and fragment size distribution is determined by loading 1 *µ*l using the Agilent Bioanalyzer HS assay. In our case, the average library size lies by about 400bp. The library size might vary depending on the used restriction enzymes.

**This is a safe stopping point. The DNA can be stored for 24 h at 4°C or long term (days to weeks) at -20°C or -80°C. Please avoid repeated thawing and freezing, as this can lead to strand breaks and thus significantly reduces the number of usable DNA molecules**.

### Computational Analysis

#### HPup - Processing Sequencing Data

In a first instance, the sequenced reads are subjected to a hairpin sequencing pipeline (HPup) that will restore the double strand information and output the methylation counts (uncorrected) for CpGs, GpCs and CpHs dyads. For this, the sequencing data was processed similar to what has been previously described by Porter *et al*. [46]. We used the Ruffus pipeline framework [47] for implementation of our workflow. In an initial step, sequencing adapter and low quality base calls (Q<20) were trimmed from the ends of each read (FastQ files) using TrimGalore! [48]. Next, all reads were screened and purged from the hairpin linker sequence using cutadapt [49]. The hairpin linker sequence is stored in an additional file and used to calculate the C-to-T conversion rate during (oxidative) bisulfite treatment and furthermore, can be used to identify redundant reads generated by PCR.

Trimmed read pairs (read 1 and read 2 from paired-end-sequencing) were locally aligned with the Smith-Waterman algorithm. We allow for G-to-A and C-to-T mismatches to cope with bisulfite treatment. After aligning the bisulfite sequence of the read pairs, we can reconstitute the genomic sequence, which allows for a faster, more efficient and precise mapping of the reads to the reference genome. In addition, the pipeline determines the methylation state of each cytosine and classifies symmetric dyads, i.e., CpGs and GpCs into fully-methylated (both DNA strands are methylated), hemi-methylated (only one strand is methylated) at the plus strand, hemi-methylated on the minus strand and unmethylated (both strands are unmethylated). The resulting sequences were aligned to the mouse genome (mm10) with GEM-mapper (beta build 1.376) [50](54), after which the methylation information was reintroduced with a custom pileup function based on HTSJDK [51] and ratios for the four methylation states were calculated for each cytosine.

The hairpin pipeline comes with a configuration file in which all required parameters can be defined by the user. This includes the path to the input- and output-directory, as well as the adapter and hairpin linker sequences for trimming. The library generates three output files per sample: (i) a log file, containing the information about the individual processing steps (parameter, success and duration of individual steps). (ii) a statistic file which provides the read count after each processing step (raw reads, trimmed, paired, aligned), as well as the conversion rates of C, 5mC, 5hmC and 5fC, calculated based on hairpin linker and spike-in sequence. Lastly, (iii) a dsi-file (double strand information), which stores the methylation information of each cytosine is generated.

The DSI file is a header-less, tab-separated text-file and contains in total 11 columns. Columns 1 to 7 contain general information: (1) the chromosome number, (2) the genomic location of the analyzed cytosine, (3) the strand of the analyzed cytosine, (4) total number of reads, (5) number of methylated reads, (6) number of unmethylated reads and (7) sequence context of the cytosine (CG, GC or non-CpG). Columns 8 to 11 contain double strand specific methylation information: (8) counts of fully methylated dyads (CpG or GpC; NA = non-CpG), (9) counts of plus-strand methylated dyads, (10) counts of minus strand methylated dyads and (11) count of unmethylated dyads. DSI files can be visualised in the Integrative Genomics Viewer (IGV). Each sample will be displayed as bar diagram-track, in which each cytosine is represented by a stacked bar summarizing the frequency of the methylation states (fully *=* orange, hemi-plus *=* dark-green, hemi-minus *=* light-green and unmethylated *=* blue). Hovering the mouse cursor above the bar will show the entire 11-column information of the DSI file for the given position.

#### GwEEP-tool - Hidden Markov modelling of single CpG methylation

We provide the DSI files containing the BS and oxBS counts for each sequenced CpG as input to an HMM analysis (GwEEP- tool). We also incorporate the (oxidative-) bisulfite conversion rates (Figure 5) in our model to account for possible conversion errorsConversion rates of BS and oxBS are derived from a set of short (60bp) synthetic DNA sequences containing C, 5mC, 5hmC and 5fC, respectively (SI Sec. 6). The computational core of our model is the HMM previously described in [52, 53].We use Bayesian inference to estimate the distribution of methylation stages given the BS and oxBS counts. We also determine posterior distributions of the corresponding methylation efficiencies. In the HMM, we combine cell division i.e. DNA replication, methlyation (*de novo* and maintenance) as well as hydroxylmethylation efficiency. Figure 6 illustrates the possible transitions with the efficiencies that are to be estimated. The advantage of our Bayesian inference approach, compared to alternative approaches such as Maximum Likelihood Estimation, is that it yields useful estimates also in the case of relatively small counts. We apply the Metropolis-Hastings algorithm to a prior distribution and sample from the posterior distribution of the Dnmts, Tets efficiencies and the methylation levels of the CpGs at three available time points. Intuitively, the estimated efficiencies reflect the values that best explain the transition from the initial to the final DNA methylation state of the ES cells by considering all time points in between. Finally, our model also provides an estimate for maintenance of 5hmC during replication i.e. the maintenance probability *p*, which determines the likelihood that a given 5hmC is recognized by Dnmt1 as “methylated” after replication. Our model is now available as an efficient and user-friendly python script: https://github.com/PaulaKramer/GwEEP-tool.

**Figure 5:**
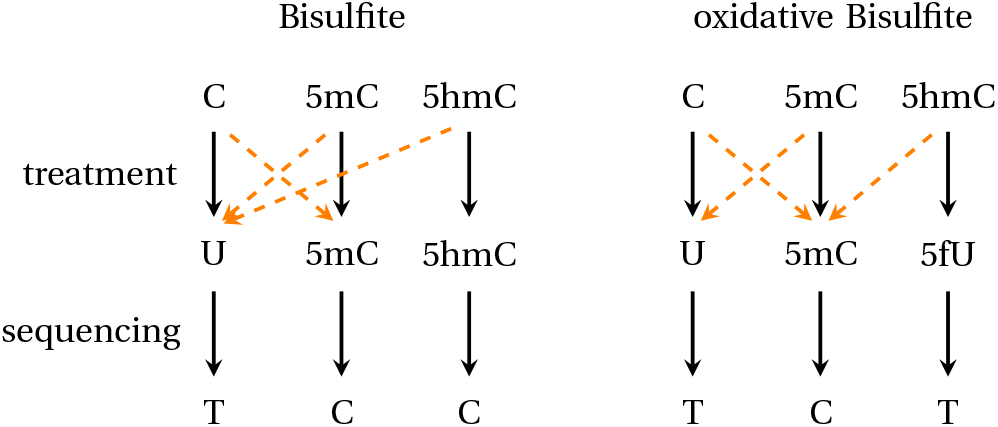
Bisulfite conversions (left) and oxidative bisulfite conversions (right). Possible conversion errors (orange arrows) can occur during the bisulfite or oxidative bisulfite sequencing approach and are considered in the modeling approach.

**Figure 6:**
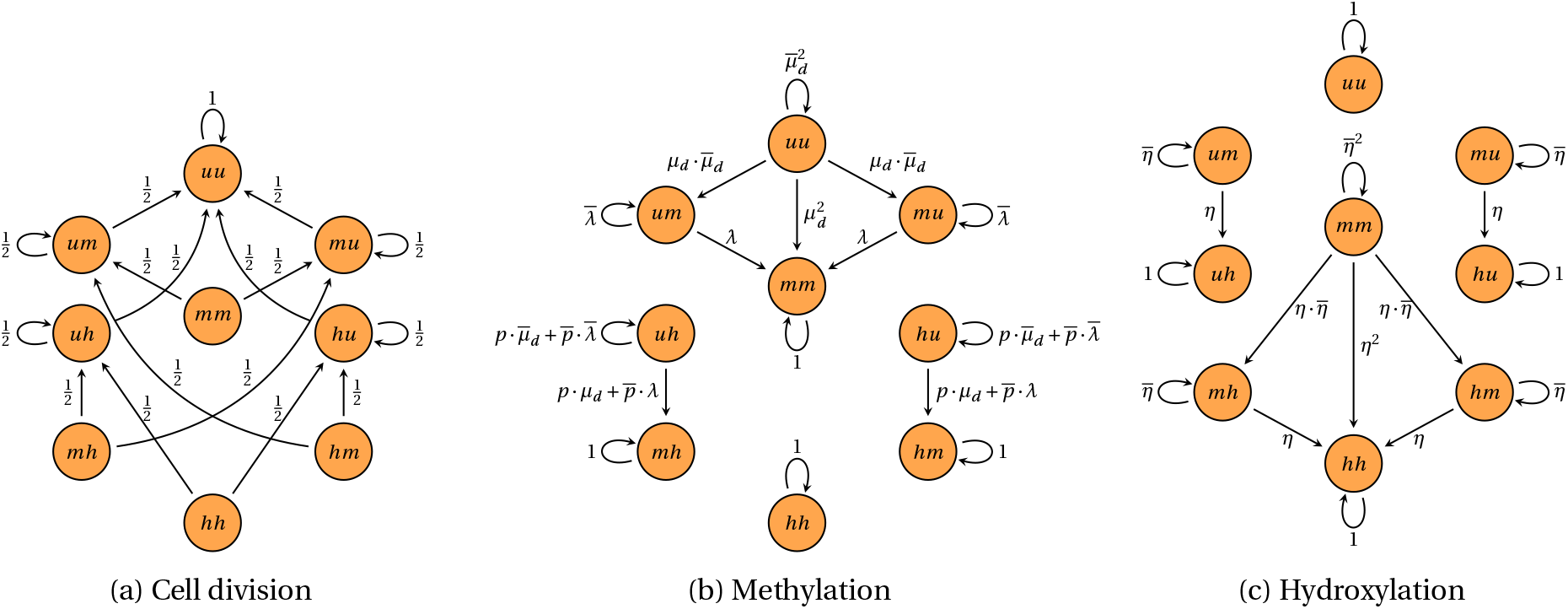
Transitions between methylation states: *u* indicates unmethlyated, *m* represents methylated and *h* are hydroxylated stages. *µ*_*d*_ describes the efficiency of *de novo* methylation, *µ*_*m*_ the efficiency of maintenance methylation, *η* hydroxylation efficiency. *λ*, the overall methylation efficiency (*de novo +* maintenance), is defined as: *λ = µ*_*m*_ *+µ*_*d*_ *-µ*_*m*_ *· µ*_*d*_. The parameter *p* describes the probability that maintenance methylation does not consider hemi-hydroxylated sites.

## RESULTS

We applied GwEEP on a well established ES cell system to precisely map 5mC and 5hmC across the genome in a time series experiment to study the enzymatic contribution of Tets and Dnmts for the progressive genome wide DNA (de)methylation. For this we generated a high resolution data set based on the above described genome wide hairpin sequencing approach: Reduced Representation Hairpin oxidative Bisulfite Sequencing (RRHPoxBS). The design of the RRHPoxBS (Combination of R.AluI, R.HaeIII and R.HpyCH4V) approach covered up to 4 million CpG dyads for which we could infer the precise distributions of 5mC and 5hmC. Following a strict read and conversion quality control, we filtered for sufficient sequencing depth and ended up with about 2 million CpGs per sample for subsequent comparative modeling. To follow the dynamics of the enzymes over time we generated six data sets for WT ES cells, i.e., BS and oxBS libraries for three different time points, starting with Serum/Lif (d0), followed by 72h 2i (d3) and 144h 2i (d6). For a comparison we also generated four datasets for Tet TKO cells starting with Serum/Lif (d0) followed by 48h in 2i (d2), 96h in 2i (d4) and 168h in 2i (d7).

### Impaired loss of 5mC in Tet TKO ES cells

Under primed conditions (Serum/LIF) WT ES cells show a level of 78% methylation. About 56% of CpGs are fully methylated while 22% (Fig.: 7.A) are found in a hemimethylated state. Among cultivation in 2i medium the DNA becomes progressively demethylated, such that after six days in 2i only 30% of CpGs retain a methylated state (fully or hemimethylated). For all time points, we find that hemimethylation is equally distributed among both DNA strands (Fig.: 7.A and SI Sec. 4.3 Figure 23). Overall, our results agree nicely with previously published whole genome methylation profiles [38]. Note, that the data from Ficz *et. al* originate from the very same ES cell sample.

**Figure 7:**
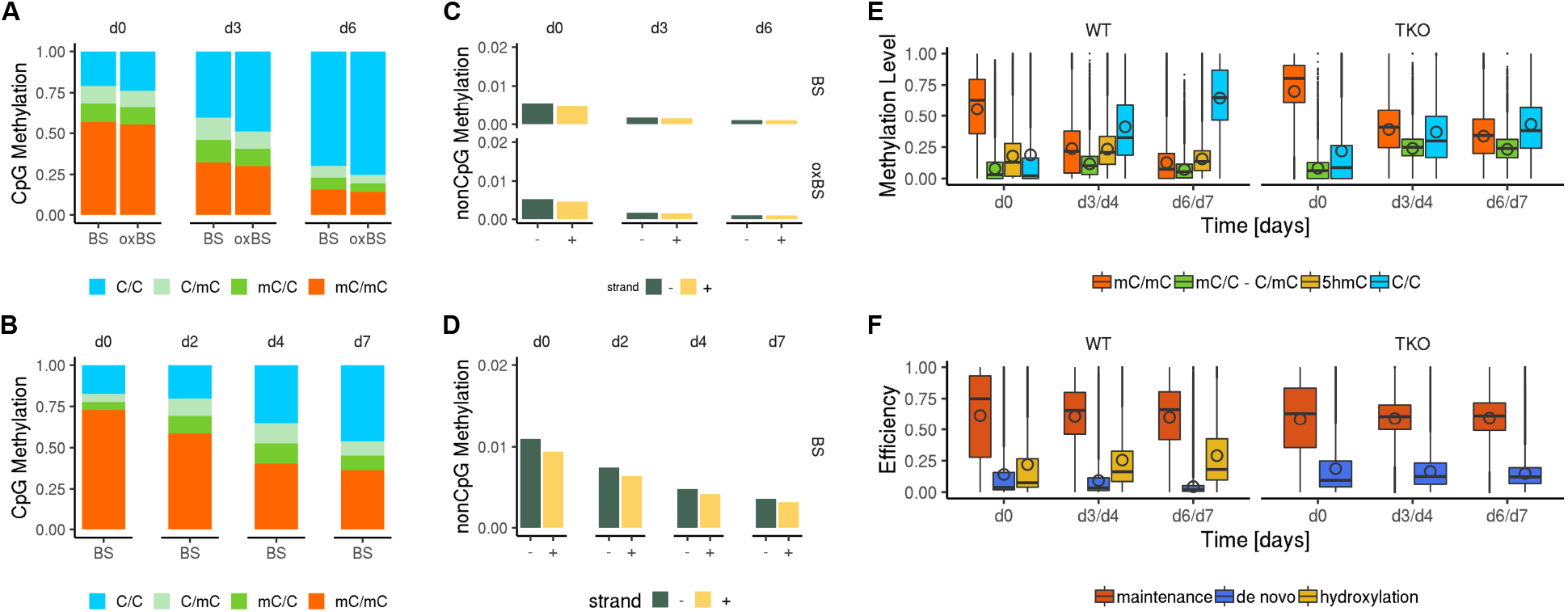
Hairpin Sequencing Data and HMM Results - **A** Average CpG methylation level based on uncorrected hairpin sequencing counts for WT ES cells; **B** Average CpG methylation level based on uncorrected hairpin sequencing counts for Tet TKO ES cells; **C** Average nonCpG methylation level based on uncorrected hairpin sequencing counts for WT ES cells; **D** Average CpG methylation level based on uncorrected hairpin sequencing counts for Tet TKO ES cells; **E** calculated and conversion error corrected 5mC and 5hmC distribution after HM modelling; **F** HMM derived maintenance methylation, *de novo* methylation and hydroxylation efficiencies.

In addition, we observe that oxBS samples always display lower methylation levels than BS samples (Fig.: 7.A). This difference corresponds to the amount of 5hmC of each sample. We detect the main difference in the hemimethylated proportion, indicating that a considerable amount of 5hmC might exist in a hemi(hydroxy)methylated (5hmC/C or C/5hmC) state (Fig.: 7.A). Initially, the amount of 5hmC is quite low but we observe a notable increase at d3, while later (d6) the amount of 5hmC decreases again (Fig.: 7.A).

ES cells lacking Tet enzymes (Tet TKO) show a marginal increase of methylated CpG dyads (82% fully or hemimethylated, Fig.: 7.B) in comparison to WT under primed serum conditions. However, in relation to WT the TKO cells show a higher frequency of fully methylated CpGs (72%) and a reduced proportion of hemimethylated CpGs (hemiCpGs; 10%) (Fig.: 7.B). We concluded that in WT ES cells, the enhanced presence of hemiCpGs is directly coupled to 5mC oxidation by Tets.

Moreover, in contrast to previous data sets, [38, 39, 54] the RRHoxBS data allow us distinguish hemi-from unmethylated CpGs and therefore to precisely estimate the demethylation kinetics, revealing that in WT ES cells the generation of unmethylated cytosine is 8% per day, while in Tet TKO cells it drops to 4.2% (SI Sec. 4.4 Figure 24). This indicates that the presence of Tets has a considerable influence on DNA demethylation kinetics.

RRHPoxBS sequencing also allowed us to accurately determine the amount, location and distribution of non-CpG methylation in (WT and Tet TKO) ES cells. In both, WT and TKO, we find CpA to be the most frequent methylated non-CpG motif SI Sec. 4.7 Figure 32 and 33). Over time, nonCpG methylation becomes gradually reduced upon cultivation in 2i. In WT ES cells the number of methylated nonCpGs was identical in BS and oxBS libraries, indicating that nonCpGs are not a substrate for Tet oxidation (Fig.: 7.C). In Tet TKO cells, the number of methylated nonCpGs is approximately doubled as compared to WT ES cells. Since nonCpG methylation is strictly dependent on the presence of *de novo* methylation activities by Dnmt3a/b [18], the higher nonCpG methylation in TKO cells, both under primed (=2%) and naive (=0.6% after 168h 2i) conditions (Fig.: 7.D), points clearly towards an increased *de novo* methylation activity by Dnmt3a/b in the absence of Tet enzymes.

### 5mC oxidation affects maintenance and *de novo* methylation

The estimated methylation levels for WT and Tet TKO ES cells fit nicely to the hairpin methylation data (SI Sec. 3, Fig.: 3), indicating a high accuracy of our model. Consequently, we observe a constant decline of fully methylated CpGs in WT ES cells over time (Fig.: 7.E).

Moreover, in WT ES cells, the HMM estimates a notable amount of 5hmC at all time points. Note that the displayed amount of 5hmC (yellow) refers to the sum of all possible 5hmC states, i.e., 5hmC/5hmC, 5hmC/5mC, 5mC/5hmC, 5hmC/C, C/5hmC. The highest amount of 5hmC is observed at d3, meaning that WT ES cells display a transient increase of 5hmC after cultivation in 2i. We observe a similar behaviour for hemiCpGs in WT ES cells. The parameter estimation by our model illustrates a mean maintenance methylation efficiency of about 61.4% at d0, which remains almost constant over time (60.1% at d6) (7.F). In contrast, *de novo* methylation efficiency shows a strong decrease (from 14.1% to 4.5% at d6) and the hydroxylation efficiency an increase (from 22.2% at d0 to 29.1% at d6) over time. This observation is in agreement with previous observations which demonstrated a reduction in RNA and protein levels of Dnmt3a/b in 2i, but an increased expression of Tet1/2 on a genome wide level [38, 54].

In Tet TKO cells maintenance efficiency lies by 58.8% at d0, which represents a marginal reduction compared to WT ES cells. Similarly to WT, maintenance efficiency remains stable over time (58.6% at d7) also in Tet TKO cells.

The most pronounced difference between WT and Tet TKO cells we see in *de novo* methylation efficiency. More specifically in Tet TKO *de novo* begins from (20.3%) at day 0 and exhibits only a slight decrease over time (16.2% at d7).

Finally, the model output also confirms the reduced demethylation rate in Tet TKO cells, previously observed in the hairpin sequencing data and suggests a substantial contribution of 5hmC and Tet enzyme on DNA demethylation. In fact, the model favors a scenario in which 5hmC is less well recognized (probability of non recognition equals p=0.66, SI Sec. 1.1) by the maintenance machinery after replication, promoting a faster demethylation process.

### Tets prevent the spreading of DNA methylation

We next related the model estimates to genomic, enzymatic and epigenetic features first focusing on Dnmt and Tet enzyme efficiencies across large genome segments with distinct methylation states. We used MethylSeekR [55], to partition the genome into four states: highly methylated regions (HMRs), partially methylated domains (PMDs), low methylated regions (LMRs) and unmethylated regions (UMRs). The segmentation was performed on whole genome bisulfite sequencing (WGBS) data from WT ES cells cultivated under Serum/Lif conditions (Ficz et al.) [38] on the identical cell batch used for our study. The estimated methylation levels (sum of 5mC and 5hmC) for WT ES cells by our model fully agreed with those derived from WGBS (SI Sec. 4.8, Fig.: 35.C and 35.E). This not only confirmed the accuracy of our model output but also denoted that we can use the precise WGBS segmentation for further analysis. We found that the majority of the WT ES cell genome (86%) consists of large HMRs (SI Sec. 4.8, Fig.: 35.A and Fig.: 35.B) followed by shorter (13%) PMDs. The residual 2% of the genome are found to be LMRs (0.4%) or UMRs (1.5%)(SI Sec. 4.8, Fig.: 35.A and Fig.: 35.B).

We assigned the 5mC and 5hmC modifications levels, their distribution at CpG dyads and the corresponding Dnmt/Tet enzyme efficiencies determined by our model to CpGs in the individual segments (Fig.: 8.A and 8.B). In WT ES cells, all segments show a continuous reduction in DNA methylation over time. This is particularly evident in segments with initial high 5mC levels, i.e., HMRs and PMDs. HMRs and PMDs also exhibit the highest amount of 5hmC and hemiCpGs, which transiently increases at d3 and d6 (Fig.: 8.A). LMRs and UMRs show different kinetics as both the amount of 5hmC and hemiCpGs constantly decline over time. The increase of hemiCpGs in HMRs and PMDs is a clear sign of impaired maintenance methylation in naive ES cells linked to the reported temporal increase in Tet expression and loss of Dnmt1 activity [38, 54].

**Figure 8:**
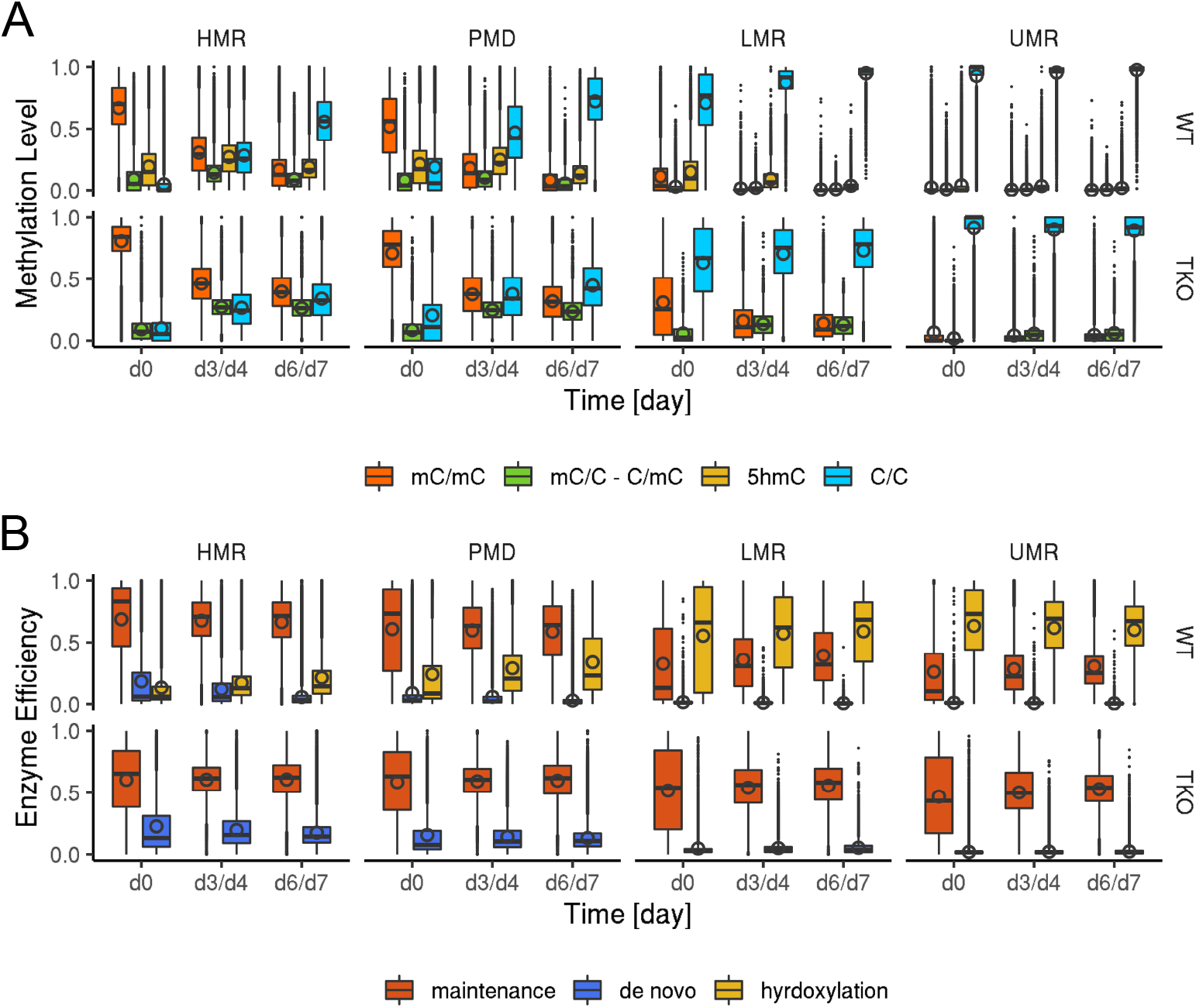
Methylation and Efficiency Profiles of Methylome Segments - (A) Estimated methylation distibution of HMRs, PMDs, LMRs and UMRs; orange = fully methylated CpGs, green = hemimethylated CpGs, yellow = CpG containing 5hmC and blue = umethylated CpGs. (B) Estimated enzyme efficiencies in HMRs, PMDs, LMRs and UMRs; red = maintenance methylation efficiency by Dnmts, blue = *de novo* methylation efficiency by Dnmts, yellow = hydroxylation efficiency by Tets

Based on the methylation data, our model predicts high maintenance methylation efficiency in HMRs (69%) and PMDs (61%), but low maintenance efficiency in LMRs (32%) and UMRs (26%)(Fig.: 8.B). Additionally, we observe a relatively high *de novo* methylation efficiency at HMRs (18%) in primed ES cells (Fig.: 8.B). Overall, *de novo* methylation efficiency strongly decreases upon cultivation in 2i, which corresponds well with the previous described loss of Dnmt3a/b under these conditions. In contrast, hydroxylation efficiency is high in UMRs (63%) and LMRs (55%), but low in HMRs (13%) and PMDs (24%)(Fig.: 8.B). Together, our results indicate regional differences and an antagonistic behaviour of Dnmts and Tets. This antagonism has been validated by estimating a robust spatial correlation measure between the efficiencies of Dnmts and Tets across the whole genome (SI Sec. 3.3, Fig.: 12).

Both WT and TKO ES cells show overall a decline of DNA methylation across all segments over time (Fig.: 8.A). However, in TKO cells, all segments retain a substantial higher frequency of fully methylated CpG dyads across all time points. This observation indicates a reduced demethylation rate in all segments under the absence of Tets. Surprisingly, under primed conditions (d0) Tet TKO cells show a higher number of unmethylated CpGs in HMRs as compared to WT ES cells.

Most importantly, comparing the Tet TKO with WT ES cells, we observe a strong change in Dnmt efficiencies. Maintenance methylation efficiency shows a reduction in HMRs and PMDs of TKO cells, while it clearly increases in LMRs and UMRs (Fig.: 8.B), resulting in almost equal maintenance activity across all segments. In the case of *de novo* methylation efficiency, we observe a more stable and slightly increased activity in all segments.

### Tets regulate Dnmts in TSS and TFBS

The genome wide antagonistic effects of Dnmts’ and Tets’ activity across segments and clusters prompted us to plot the enzymatic efficiencies of CpGs across genes, histone marks or ChIP profiled TFBS using DeepTools [56] (Fig.: 9) in order to investigate regularities and general local dependencies. In WT cells the enzymes’ efficiencies across genes and TFBS show once more an opposing behavior: At transcription start sites (TSS) and TFBS, high hydroxylation efficiency is coupled to reduced methylation (both maintenance and *de novo*) efficiency. This inverse behavior at TSS remains upon 2i cultivation. *De novo* methylation almost disappears across the entire gene including the gene body. Under primed conditions *de novo* methylation is absent in TSS but has a strong presence in the gene body and it almost disappears from the entire gene over time after the transition to 2i. The observed efficiency profiles for maintenance methylation, de novo methylation and hydroxylation, correspond nicely to Uhrf1, Dnmt3a/b, as well as Tet1 ChIP profiles, respectively (SI Sec. 4.6 Fig. 30 and 31).

**Figure 9:**
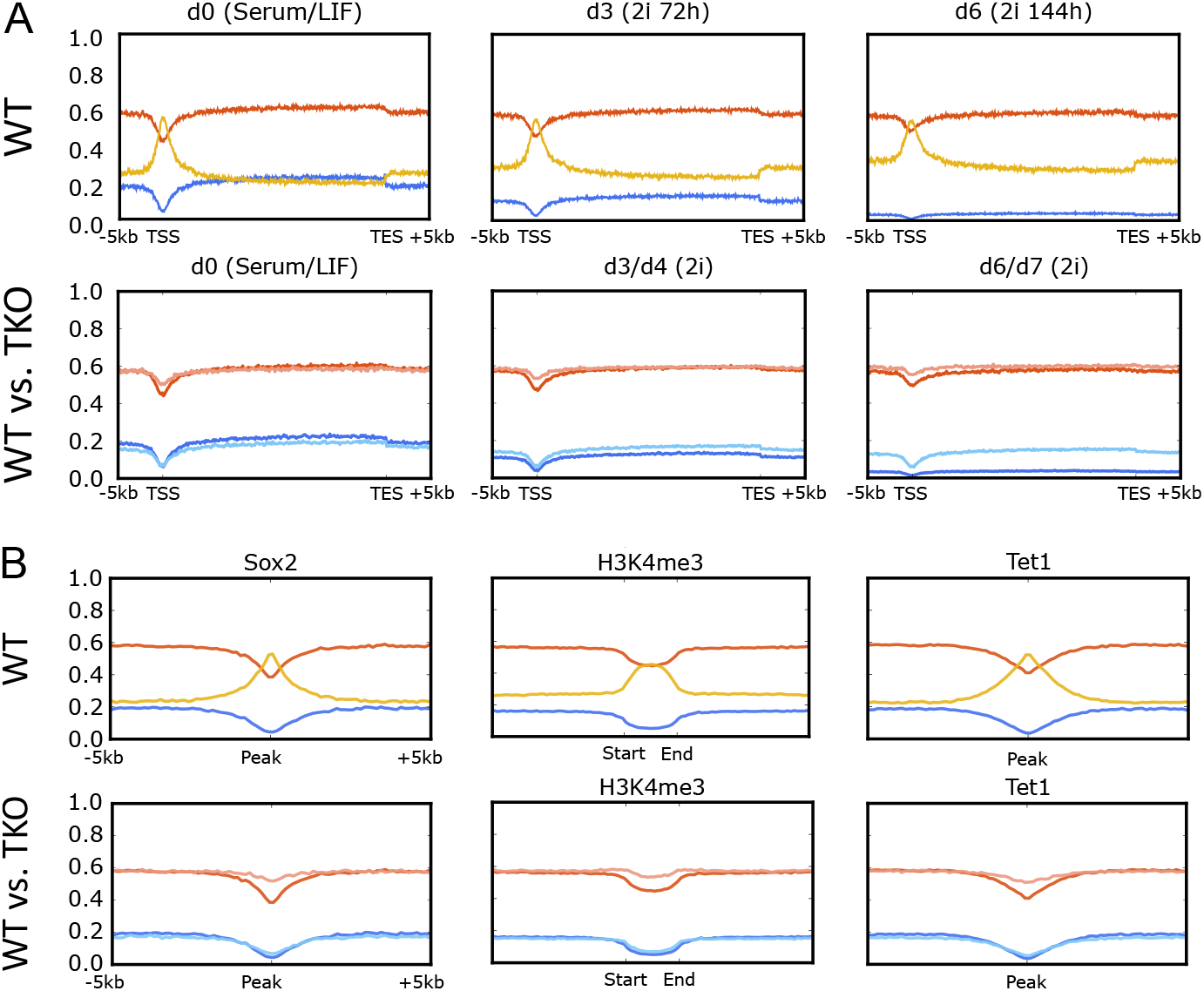
Average efficiency profiles across genes and protein binding sites. (A) Average maintenance, *de novo* and hydroxylation efficiency of WT and Tet TKO cells across genes; (B) Average maintenance, de novo, as well as hydroxylation efficiency across selected chromatin marks and protein binding sites. red = maintenance methylation efficiency, blue = *de novo* methylation efficiency, yellow = hydroxylation efficiency. Dark colors indicate the efficiencies in WT ES cells, while light colors refer to the efficiencies of Tet TKO ES cells.

In Tet TKO ES cells the TSS associated drop in maintenance methylation is much less pronounced and almost absent at d6/d7. In addition, *de novo* methylation is only mildly reduced upon cultivation in 2i and clearly maintained across the gene body (Fig.: 8.A). Regulatory regions marked by Sox2, H3K4me3 and Tet1 enrichment show a strong hydroxylation activity in WT cells which is inversely linked to an impaired maintenance and de novo methylation activity. Interestingly, the lack of Tet activity in TKO cells does not change de novo methylation but maintenance activity across regulatory regions (Fig.: 8.B).

## DISCUSSION

In this study, we provide a comprehensive approach for measuring genome wide DNA-methylation and efficiency profiling. GwEEP allowed us to infer how the activity of Dnmts and Tets contribute to modify CpGs and nonCpGs across the genome in a functional context. It is important to note that technically our RRHPoxBS data resemble and corroborate the overall methylation dynamics observed by classical RRBS and WGBS ([38, 54]). However, RRHPoxBS data provide three important novel features: (i) a genome-wide representation of up to 4 million CpGs uniformly distributed across the genome, (ii) a precise determination of 5mC and 5hmC levels at a single CpG dyad and (iii) a precise mapping of hemimethylated states and positions of nonCpG methylation. The combination with our HMM allows us to calculate accurate 5mC and 5hmC level by considering the conversion errors during BS and oxBS. Furthermore, compared to previous models, we simultaneously infer the genome wide efficiencies of maintenance methylation, *de novo* methylation and hydroxylation efficiencies.

The overall evaluation of our RRHPoxBS data showed that in mouse ES cells (as previously described for somatic cells [18]), hemiCpGs are almost equally distributed on both DNA strands, following the behavior of symmetric CpG methylation. This suggests that hemimethylation is most likely the result of (strand-) undirected *de novo* methylation or active and passive demethylation events, respectively. Furthermore, we detect more hemimethylation in WT compared to Tet TKO cells, which indicates that Tet enzymes enhance the passive loss of 5mC. Indeed, our model predicts that 5hmC is probably less well recognized by Dnmt1 after replication, such that hydroxylation enhances passive demethylation, which is in agreement with recent in vitro studies [30, 31]. In contrast to equally distributed hemimethylation we observe a slight increase in the minus strand presence of nonCpG methylation. We cannot find a simple biological (sequence context) or technical (calling/mapping) explanation for this bias. Non-CpG methylation is always occurring in close vicinity to CpG methylation (18) but in contrast to CpGs we find that nonCpGs are not a substrate for Tet enzymes, i.e., we do not find any indication of 5hmC in the non-CpG context. The amount of nonCpG methylation however is strongly enhanced in the absence of Tet enzymes, suggesting an increase of Dnmt3a and 3b efficiency in the absence of Tets. Our model provides strong evidence that Dnmts and Tets do not act independently at a given CpG, but clearly in an opposed manner. Generally, we observe a high maintenance and de novo efficiency at the majority of the genome, i.e., HMRs and PMDs (or inter-/intragenic regions), while the activity of Tet enzymes is highest at UMRs and LMRs, such as promoters, TFBS (Sox2, Pou5f1) and TSS. Recent studies based on chromatin immunoprecipitation support our findings, revealing binding of Dnmt3a/b at the gene body and HMRs, whereas Tet1 binding was observed across methylation valleys (LMRs and UMRs) [41, 57].

The impairment of maintenance methylation has been identified so far as the main driver of 2i induced DNA demethylation [54] and a role for Tet or oxidative cytosine forms, on the other hand, has only been recognized for selected loci [38, 54]. The comparison of WT and Tet TKO ES cells in the present study, however, discloses a notable reduction within the demethylation rate of Tet TKO, compared to WT ES cells. On average, we detect a reduction in the demethylation rate of almost 50% from around 8% to 4% loss per day. This indicates that Tets and their oxidized cytosine products are essential for an effective demethylation during the Serum-to-2i shift and probably other biological demethylation processes with similar enzymatic compositions.

The loss of Tet enzymes is naturally expected to result in an impaired removal of 5mC and it does at least for CpGs located in LMRs and UMRs, where we observe a notable increase in their methylation level. Nevertheless, under primed conditions and within HMRs, we paradoxically observe more unmethylated CpGs (hypomethylation) in Tet TKO ES cells compared to WT ES cells. Recently, L’opez-Moyado *et al*. conducted a systematic investigation of genome wide methylation profiles from various cell types carrying distinct Tet KO genotypes [42]. Similar to our observations they detected a pronounced loss of DNA methylation in heterochromatic compartments (i.e., HMRs and PMDs) of Tet TKO mouse ES cells.

L’opez-Moyado *et al*. propose a mutual exclusive localization of Dnmts and Tets in WT ES cells, while in Tet KO cells Dnmts invade domains which were previously occupied by Tets. Indeed, in the absence of Tets, our model predicts a clear misregulation in both maintenance and de novo methylation efficiency. In Tet TKO ES cells, we see an increase in maintenance methylation efficiency, but at the same time a reduction in HMRs and PMDs. Moreover, we observe an increase in de novo methylation efficiency at PMDs. Together, this indicates a displacement of Dnmt1, as well as Dnmt3a/b, which fits to the hypothesized model by L’opez-Moyado *et al*.. In addition, Tet TKO cells exhibit a more stable, almost persistent de novo methylation under naive conditions. The increased non-CpG methylation of Tet TKO cells detected by RRHPoxBS further supports this finding. This shows that in the absence of Tets, ES cells also fail to effectively down-regulate de novo methylation efficiency in 2i.

Taken together, we summarize that Tet enzymes work against methylation in three ways. (i) They guarantee an efficient conversion of 5mC at accessible regions and act against its establishment during a cell replication either via passive or active demethylation, (ii) They inhibit the effectiveness of the maintenance machinery over regions that should remain unmethylated. (iii) Finally, they ensure an efficient down-regulation of the de novo enzymes, which can not be observed in their absence.

## CONCLUSION

We describe a novel combination of experimental and computational approach - GwEEP to investigate the contributions of Tets and Dnmts to the establishment of distinctive DNA methylation patterns across the genome. In GwEEP, we generate strand specific (hydroxy)methylation data and using a sophisticated HMM we infer the distriution of 5mC and 5hmC at individual CpGs across the genome and furthermore, derive accurate efficiency profiles of Dnmts (*de novo* and maintenance) and Tets (hydroxylation). GwEEP also works with low amounts of DNA and is there fore suitable for demanding samples and rare cell types. Moreover, by combining our hairpin protocol with 5fC or 5caC detecting chemistry, GwEEP is easily expandable for the estimation of 5fC/5caC distribution and the inference of formylation and carboxylation efficiencies of Tets. Our analysis of WT and Tet TKO mES cells shows that Dnmts and Tets exhibit clear antagonistic efficiencies at individual CpGs. The comparison of WT and Tet TKO ES cells demonstrates that Tet enzymes contribute notably to the loss of DNA methylation in the present model system. Moreover, Tet enzymes seem to protect unmethylated regions against both *de novo* and maintenance methylation efficiency and to restrict the activity of Dnmts in highly methylated regions, guaranteeing the formation and maintenance of cell type specific methylation patterns.

## DATA AVAILABILITY

All raw and processed hairpin sequencing data sets, as well as the HMM output are deposited at NCBI’s Gene Expression Omnibus (GEO) repository under accession number GSE169070.

## ACKNOWLEDGEMENTS

We thank Jasmin Kirch and Gilles Gasparoni for performing sequencing using the Illumina HiSeq2500 system and Andreas Firczynski for the optimization of the processing pipeline for the hairpin sequencing data. Particular thanks to Jim Robinson and the entire IGV team for implementing the dsi-file format into the IGV genome browser.

## AUTHOR’S CONTRIBUTIONS

Conceived and designed the experiments: CK PG WR VW JW. Wrote the manuscript: CK PG JW VW. Performed the experiments: PG JA FvM GF. Processed the raw data: KN. Designed/implemented the computational methods: CK. Analyzed the data: CK. Performed meta analysis: CK PG KN AS FM.

## SOFTWARE AVAILABILITY

All Matlab scripts written for implementing the single CpG stochastic model, run it in a parallel fashion on a multi-core environment and perform the subsequent computational analysis presented in the manuscript and the appendix are shared via GitHub (https://github.com/kyriakopou/hydroxyGit/tree/master/genomeWide)

