## Appendix for "GwEEP - A comprehensive approach for genome-wide efficiency profiling of DNA modifying enzymes"

### 1 Hidden Markov Model (HMM)

We use the hidden Markov model (HMM) presented in [1] to describe the temporal evolution of a single CpG dyad over time for each experiment (bisulfite or oxidative bisulfite). The hidden states of the model correspond to the different modifications, e.g. the cytosines (C) on both strands are unmethylated or the C on the upper strand is methylated while the C on the lower strand is unmethylated, etc. The observable states are those that we measure after bisulfite (BS) or oxidative bisulfite (oxBS) hairpin sequencing. Hence, we include the conversion errors of the measurement process and we link two HMMs that describe oxidative and non-oxidative hairpin bisulfite sequencing to accurately determine hydroxymethylation levels and the efficiencies of the involved enzymes over time. Formally, we define the sets of hidden states $\mathcal{S} = \{u, m, h\}^2$ and the set of observable states $\mathcal{S}_{obs} = \{\mathrm{T}, \mathrm{C}\}^2$. A state $s \in \mathcal{S}$ describes whether the upper and the lower strand of the site is *unmethylated* $(u)$, *methylated* $(m)$ or *hydroxylated* $(h)$. E.g. in state $(h, u)$ the upper strand is hydroxylated and the lower strand is unmethylated. Similarly, a state $j \in \mathcal{S}_{obs}$ encodes whether the upper strand and the lower strand of the site has been transformed after the BS or the oxBS treatment to a thymine (T) or a cytosine (C). We use the abbreviation $hu$ for state $(h, u)$ and similar for all other states.

#### 1.1 Distribution of Hidden and Observable States

Let the vector $\pi(t)$ be the hidden states distribution at time $t$ and let $\pi(i,t) = P(\mathcal{X}(t) = i)$ represent the entry of $\pi(t)$ that corresponds to state $i \in S$.

The transition matrix of the hidden states is defined as $\mathbf{P}(t) = \mathbf{D}(t) \cdot \mathbf{M}(t) \cdot \mathbf{H}(t)$, where $\mathbf{D}(t)$ describes the modifications due to cell division, $\mathbf{M}(t)$ the modifications due to methylation, and $\mathbf{H}(t)$ the modifications due to hydroxymethylation. As in [1], we define

$$\mathbf{D}(t) = \begin{array}{c} \\ uu \\ um \\ mu \\ uh \\ hu \\ hm \\ mh \\ mm \\ hh \end{array} \begin{array}{c} \begin{array}{ccccccccc} uu & um & mu & uh & hu & hm & mh & mm & hh \end{array} \\ \begin{pmatrix} 1 & 0 & 0 & 0 & 0 & 0 & 0 & 0 & 0 \\ 1/2 & 1/2 & 0 & 0 & 0 & 0 & 0 & 0 & 0 \\ 1/2 & 0 & 1/2 & 0 & 0 & 0 & 0 & 0 & 0 \\ 1/2 & 0 & 0 & 1/2 & 0 & 0 & 0 & 0 & 0 \\ 1/2 & 0 & 0 & 0 & 1/2 & 0 & 0 & 0 & 0 \\ 0 & 1/2 & 0 & 0 & 1/2 & 0 & 0 & 0 & 0 \\ 0 & 0 & 1/2 & 1/2 & 0 & 0 & 0 & 0 & 0 \\ 0 & 1/2 & 1/2 & 0 & 0 & 0 & 0 & 0 & 0 \\ 0 & 0 & 0 & 1/2 & 1/2 & 0 & 0 & 0 & 0 \end{pmatrix} \end{array},$$

$$
\mathbf{M}(t) = \begin{array}{c} \\ uu \\ um \\ mu \\ uh \\ hu \\ hm \\ mh \\ mm \\ hh \end{array}
\begin{array}{c}
\begin{array}{ccccccccc} uu & um & mu & uh & hu & hm & mh & mm & hh \end{array} \\
\begin{pmatrix}
\bar{\mu}_d^2 & \mu_d \cdot \bar{\mu}_d & \mu_d \cdot \bar{\mu}_d & 0 & 0 & 0 & 0 & \mu_d^2 & 0 \\
0 & \bar{\lambda} & 0 & 0 & 0 & 0 & 0 & \lambda & 0 \\
0 & 0 & \bar{\lambda} & 0 & 0 & 0 & 0 & \lambda & 0 \\
0 & 0 & 0 & p \cdot \bar{\mu}_d + \bar{p} \cdot \bar{\lambda} & 0 & 0 & p \cdot \mu_d + \bar{p} \cdot \lambda & 0 & 0 \\
0 & 0 & 0 & 0 & p \cdot \bar{\mu}_d + \bar{p} \cdot \bar{\lambda} & p \cdot \mu_d + \bar{p} \cdot \lambda & 0 & 0 & 0 \\
0 & 0 & 0 & 0 & 0 & 1 & 0 & 0 & 0 \\
0 & 0 & 0 & 0 & 0 & 0 & 1 & 0 & 0 \\
0 & 0 & 0 & 0 & 0 & 0 & 0 & 1 & 0 \\
0 & 0 & 0 & 0 & 0 & 0 & 0 & 0 & 1
\end{pmatrix}
\end{array},
$$

and

$$
\mathbf{H}(t) = \begin{array}{c} \\ uu \\ um \\ mu \\ uh \\ hu \\ hm \\ mh \\ mm \\ hh \end{array}
\begin{array}{c}
\begin{array}{ccccccccc} uu & um & mu & uh & hu & hm & mh & mm & hh \end{array} \\
\begin{pmatrix}
1 & 0 & 0 & 0 & 0 & 0 & 0 & 0 & 0 \\
0 & \bar{\eta} & 0 & \eta & 0 & 0 & 0 & 0 & 0 \\
0 & 0 & \bar{\eta} & 0 & \eta & 0 & 0 & 0 & 0 \\
0 & 0 & 0 & 1 & 0 & 0 & 0 & 0 & 0 \\
0 & 0 & 0 & 0 & 1 & 0 & 0 & 0 & 0 \\
0 & 0 & 0 & 0 & 0 & \bar{\eta} & 0 & 0 & \eta \\
0 & 0 & 0 & 0 & 0 & 0 & \bar{\eta} & 0 & \eta \\
0 & 0 & 0 & 0 & 0 & \eta \cdot \bar{\eta} & \eta \cdot \bar{\eta} & \bar{\eta}^2 & \eta^2 \\
0 & 0 & 0 & 0 & 0 & 0 & 0 & 0 & 1
\end{pmatrix}
\end{array}.
$$

Here, $\mu_m$ stands for the maintenance efficiency, $\mu_d$ for *de novo* and $\eta$ for the hydroxylation efficiency, while $p$ is the probability that 5hmC is not considered during maintenance (see [1] for details).

Note, that for $\mathbf{D}(t)$ we can omit the time parameter $t$ since it is time-independent, while the other two matrices depend on $t$ as explained later. Note also, that the HMMs of BS and oxBS experiments have both the same distribution $\pi(t)$ for the hidden states (as for both experiments the same cell population is used) but different emission probabilities and that $\pi(t)$ is given by

$$\pi(t) = \pi(0) \cdot \prod_{k=1}^{t} \mathbf{P}(k).$$

Let the vectors $\pi_{bs}(t), \pi_{ox}(t)$ be the observable states distribution at time $t$, with entries $\pi_{bs}(j,t)$ and $\pi_{ox}(j,t)$, $j \in \mathcal{S}_{obs}$, for the BS and oxBS experiments, respectively. We then get:

$$\pi_{bs}(t) = \pi(t) \cdot \mathbf{E}_{bs}(t) \quad \text{and} \quad \pi_{ox}(t) = \pi(t) \cdot \mathbf{E}_{ox}(t),$$

where the entries of the emission matrices $\mathbf{E}_{bs}(t)$ and $\mathbf{E}_{ox}(t)$ are given in Table A.

### 2 Maximum Likelihood Estimation (MLE)

#### 2.1 Initial distribution of the hidden states

Let $n_{bs}(j,t)$ and $n_{ox}(j,t)$ be the number of times that state $j \in \mathcal{S}_{obs}$ has been observed during independent hairpin bisulfite (BS) and oxidative hairpin bisulfite (oxBS) measurements out of a certain number of reads (mean coverage of all samples $\approx$ 20x) at time $t$.

Since we assume that $t = 0$ is the time of the first measurement, we have observations at $t = 0$ and can estimate the unknown initial distribution over the hidden states using maximum likelihood estimation (MLE). For this, we have to solve the optimization problem: $\pi(0)^* = \arg\max_{\pi(0)} \mathcal{L}_1(\pi(0))$, subject to the constraint $\sum_{i \in \mathcal{S}} \pi(i,0) = 1$, where

$$\mathcal{L}_1(\pi(0)) = \prod_{j \in \mathcal{S}_{obs}} \pi_{bs}(j,0)^{n_{bs}(j,0)} \cdot \pi_{ox}(j,0)^{n_{ox}(j,0)}.$$

| | BS | | | | oxBS | | | |
|---|---|---|---|---|---|---|---|---|
| | TT | TC | CT | CC | TT | TC | CT | CC |
| $uu$ | $c^2$ | $c \cdot \bar{c}$ | $c \cdot \bar{c}$ | $\bar{c}^2$ | $c^2$ | $c \cdot \bar{c}$ | $c \cdot \bar{c}$ | $\bar{c}^2$ |
| $um$ | $c \cdot \bar{d}$ | $c \cdot d$ | $\bar{c} \cdot \bar{d}$ | $\bar{c} \cdot d$ | $c \cdot \bar{d}$ | $c \cdot d$ | $\bar{c} \cdot \bar{d}$ | $\bar{c} \cdot d$ |
| $mu$ | $c \cdot \bar{d}$ | $\bar{c} \cdot \bar{d}$ | $c \cdot d$ | $\bar{c} \cdot d$ | $c \cdot \bar{d}$ | $\bar{c} \cdot \bar{d}$ | $c \cdot d$ | $\bar{c} \cdot d$ |
| $uh$ | $c \cdot \bar{e}$ | $c \cdot e$ | $\bar{c} \cdot \bar{e}$ | $\bar{c} \cdot e$ | $c \cdot f$ | $c \cdot \bar{f}$ | $\bar{c} \cdot f$ | $\bar{c} \cdot \bar{f}$ |
| $hu$ | $c \cdot \bar{e}$ | $\bar{c} \cdot \bar{e}$ | $c \cdot e$ | $\bar{c} \cdot e$ | $c \cdot f$ | $\bar{c} \cdot f$ | $c \cdot \bar{f}$ | $\bar{c} \cdot \bar{f}$ |
| $hm$ | $\bar{d} \cdot \bar{e}$ | $d \cdot \bar{e}$ | $\bar{d} \cdot e$ | $d \cdot e$ | $\bar{d} \cdot f$ | $d \cdot f$ | $\bar{d} \cdot \bar{f}$ | $\bar{d} \cdot f$ |
| $mh$ | $\bar{d} \cdot \bar{e}$ | $\bar{d} \cdot e$ | $d \cdot \bar{e}$ | $d \cdot e$ | $\bar{d} \cdot f$ | $\bar{d} \cdot \bar{f}$ | $d \cdot f$ | $\bar{d} \cdot f$ |
| $mm$ | $\bar{d}^2$ | $\bar{d} \cdot d$ | $d \cdot \bar{d}$ | $d^2$ | $\bar{d}^2$ | $\bar{d} \cdot d$ | $d \cdot \bar{d}$ | $d^2$ |
| $hh$ | $\bar{e}^2$ | $\bar{e} \cdot e$ | $e \cdot \bar{e}$ | $e^2$ | $f^2$ | $f \cdot \bar{f}$ | $f \cdot \bar{f}$ | $\bar{f}^2$ |

Table A: Transition probabilities from hidden to the observable states in bisulfite sequencing (BS) and in oxidative bisulfite sequencing (oxBS).

During the optimization procedure, we use the log-likelihood

$$\ln \mathcal{L}_1(\pi(0)) = \sum_{j \in \mathcal{S}_{obs}} n_{bs}(j,0) \cdot \ln \pi_{bs}(j,0) + n_{ox}(j,0) \cdot \ln \pi_{ox}(j,0)).$$

Moreover, to allow gradient descent optimization we also compute the derivative w.r.t. $\pi(0)$ given by

$$\frac{d}{d\pi(0)} \ln \mathcal{L}_1(\pi(0)) = \sum_{j \in \mathcal{S}_{obs}} n_{bs}(j,0) \cdot \frac{\frac{d}{d\pi(0)}\pi_{bs}(j,0)}{\pi_{bs}(j,0)} + n_{ox}(j,0) \cdot \frac{\frac{d}{d\pi(0)}\pi_{ox}(j,0)}{\pi_{ox}(j,0)}. \quad (1)$$

Writing the vectors of partial derivatives $\frac{d}{d\pi(0)}\pi_{bs}(j,0)$ and $\frac{d}{d\pi(0)}\pi_{ox}(j,0)$ in a vector-matrix notation including all $j \in \mathcal{S}_{obs}$ we get

$$\frac{d}{d\pi(0)}\pi_{bs}(0) = \frac{d}{d\pi(0)}\pi(0) \cdot \mathbf{E}_{bs}(0) = \mathbf{E}_{bs}(0), \quad \frac{d}{d\pi(0)}\pi_{ox}(0) = \frac{d}{d\pi(0)}\pi(0) \cdot \mathbf{E}_{ox}(0) = \mathbf{E}_{ox}(0),$$

which after insertion into Eq. 1 gives us the gradient of the log-likelihood function w.r.t. the initial distribution of the hidden states.

#### 2.2 Estimation of the efficiencies

Let $\mathbf{v} = (\beta_0^{\mu_m}, \beta_1^{\mu_m}, \beta_0^{\mu_d}, \beta_1^{\mu_d}, \beta_0^{\eta}, \beta_1^{\eta}, p) \in \mathbb{R}^v$, be the vector of seven, i.e., $v = 7$, unknown parameters. We assume here that the efficiencies are linear functions of time (except for $p$) and so $\mathbf{v}$ contains the coefficients of these functions, e.g., $\mu_m(t) = \beta_0^{\mu_m} + t \cdot \beta_1^{\mu_m}$.

Now, after determining $\pi(0)$, (see section 2.1) we want to compute the MLE $\mathbf{v}^* = \mathrm{argmax}_{\mathbf{v}} \log \mathcal{L}_2(\mathbf{v})$, where

$$\mathcal{L}_2(\mathbf{v}) = \prod_{t \in T_{obs} \setminus \{0\}} \prod_{j \in \mathcal{S}_{obs}} \pi_{bs}(j,t)^{n_{bs}(j,t)} \cdot \pi_{ox}(j,t)^{n_{ox}(j,t)}. \quad (2)$$

Note here we assume that the cells divide every 24 hours, hence $t$ ranges over all days at which measurements were made after day0. In addition to derive the likelihood of Eq. 2 we assume that all observations made at time points $t \in T_{obs} \setminus \{0\}$ are independent. The independence assumption is well justified since during the measurement only a very small fraction of cells is taken out of a large pool and hence it is unlikely that we pick two cells with a common descendant.

Since the efficiencies are probabilities we have the constraint that for all time points in $T_{obs}$ and all efficiencies we have $0 \leq \beta_0 + \beta_1 \cdot t \leq 1$. In addition, $0 \leq p \leq 1$.

It holds

$$\ln \mathcal{L}_2(\mathbf{v}) = \sum_{t \in T_{obs} \setminus \{0\}} \sum_{j \in \mathcal{S}_{obs}} n_{bs}(j,t) \cdot \ln \pi_{bs}(j,t) + n_{ox}(j,t) \cdot \ln \pi_{ox}(j,t)$$

and we get the score vector of the log-likelihood function as

$$\frac{d}{d\mathbf{v}} \ln \mathcal{L}_2(\mathbf{v}) = \sum_{t \in T_{obs} \setminus \{0\}} \sum_{j \in \mathcal{S}_{obs}} n_{bs}(j,t) \cdot \frac{\frac{d}{d\mathbf{v}} \pi_{bs}(j,t)}{\pi_{bs}(j,t)} + n_{ox}(j,t) \cdot \frac{\frac{d}{d\mathbf{v}} \pi_{ox}(j,t)}{\pi_{ox}(j,t)}.$$

Then the matrix-vector form of the derivatives $\frac{d}{d\mathbf{v}}\pi_{bs}(j,t)$ and $\frac{d}{d\mathbf{v}}\pi_{ox}(j,t)$ can be written as

$$\frac{d}{d\mathbf{v}}\pi_{bs}(t) = \frac{d}{d\mathbf{v}}\pi(t) \cdot \mathbf{E}_{bs}(t) \text{ and } \frac{\mathrm{d}}{\mathrm{d}\mathbf{v}}\pi_{\mathrm{ox}}(\mathrm{t}) = \frac{\mathrm{d}}{\mathrm{d}\mathbf{v}}\pi(\mathrm{t}) \cdot \mathbf{E}_{\mathrm{ox}}(\mathrm{t}), \ \forall \mathrm{t} \in \mathrm{T}_{\mathrm{obs}}.$$

Considering now, the forward Kolmogorov equation for the HMM and its derivative w.r.t. the parameters it suffices to simultaneously solve the following two equation systems.

$$\begin{aligned} \pi(t) &= \pi(t-1) \cdot \mathbf{P}(t) \\ \frac{d}{d\mathbf{v}}\pi(t) &= \frac{d}{d\mathbf{v}}\pi(t-1) \cdot \mathbf{P}(t) + \pi(t-1)\frac{d}{d\mathbf{v}}\mathbf{P}(t), \ \forall t \geq 1 \end{aligned} \tag{3}$$

with $\frac{d}{d\mathbf{v}}\pi(0) = 0$ and $\pi(0) = \pi(0)^*$. The derivative of the transition matrix is

$$\frac{d}{d\mathbf{v}}\mathbf{P}(t) = \frac{d}{d\mathbf{v}}(\mathbf{D} \cdot \mathbf{M}(t) \cdot \mathbf{H}(t)) = \mathbf{D} \cdot \Big(\frac{d}{d\mathbf{v}}\mathbf{M}(t) \cdot \mathbf{H}(t) + \mathbf{M}(t) \cdot \frac{d}{d\mathbf{v}}\mathbf{H}(t)\Big).$$

Now, applying the chain rule and taking into account that $\mu_m = \beta_0^{\mu_m} + \beta_1^{\mu_m} t$ we get for the entry that corresponds to $\beta_0^{\mu_m}$

$$\frac{d}{d\beta_0^{\mu_m}}\mathbf{M}(\mu_m) = \frac{d}{d\mu_m}\mathbf{M}(\mu_m) \cdot \frac{d}{d\beta_0^{\mu_m}}\mu_m = \frac{d}{d\mu_m}\mathbf{M}(\mu_m)$$

and

$$\frac{d}{d\beta_1^{\mu_m}}\mathbf{M}(\mu_m) = \frac{d}{d\mu_m}\mathbf{M}(\mu_m) \cdot \frac{d}{d\beta_1^{\mu_m}}\mu_m = \frac{d}{d\mu_m}\mathbf{M}(\mu_m) \cdot t.$$

In a similar fashion we get the first derivatives w.r.t. all the other components of parameter vector $\mathbf{v}$. Applying once more the product rule in Eq. (3), and using similar arguments as above we can additionally compute the second partial derivatives $\frac{d}{d\mathbf{v}_i d\mathbf{v}_j} \ln \mathcal{L}_2(\mathbf{v})$, which will give us the $(i,j)$-th entry of the Hessian matrix $\mathcal{H} = \nabla\nabla^{\mathrm{T}} \ln \mathcal{L}_2(\mathbf{v})$.

##### Standard deviations and confidence intervals

The observed Fisher information is defined as $\mathcal{J}(\mathbf{v}^*) = -\mathcal{H}(\mathbf{v}^*)$, where $\mathbf{v}^*$ is the maximum likelihood (ML) estimator. We use the inverse of the expected Fisher information $\mathcal{I}(\mathbf{v}) = \mathbb{E}[\mathcal{J}(\mathbf{v})]$ to estimate the covariance matrix of the MLE. Hence, we approximate the covariance of our ML estimator as $\Sigma_{\mathbf{v}^*} = -\mathcal{H}^{-1}(\mathbf{v}^*))$. Then, in order to approximate the standard deviations of the efficiencies' functions over time, i.e., $\sigma(\mu_m(t)), \sigma(\mu_d(t))$ and $\sigma(\eta(t))$, we exploit the identity that if $f(t) = \beta_0 + \beta_1 \cdot t$ then

$$\sigma(f(t)) = \sqrt{\mathrm{Var}(\beta_0 + \beta_1 \cdot \mathrm{t})} = \sqrt{\mathrm{Var}(\beta_0) + \mathrm{t}^2\mathrm{Var}(\beta_1) + 2\mathrm{t}\mathrm{Cov}(\beta_0, \beta_1)}.$$

We then determine the confidence intervals for a fixed confidence level $\beta = 95\%$. For instance the confidence interval for the maintenance methylation function will be

$$\mu_m(t) \pm z \cdot \sigma(\mu_m(t))$$

where $z = F^{-1}\big(\frac{\beta+1}{2}\big)$ and $F$ is the cumulative distribution function (cdf) of the standard normal distribution. Similarly, we get the confidence intervals for all remaining parameters.

### 3 Bayesian Inference for Whole Genome Data

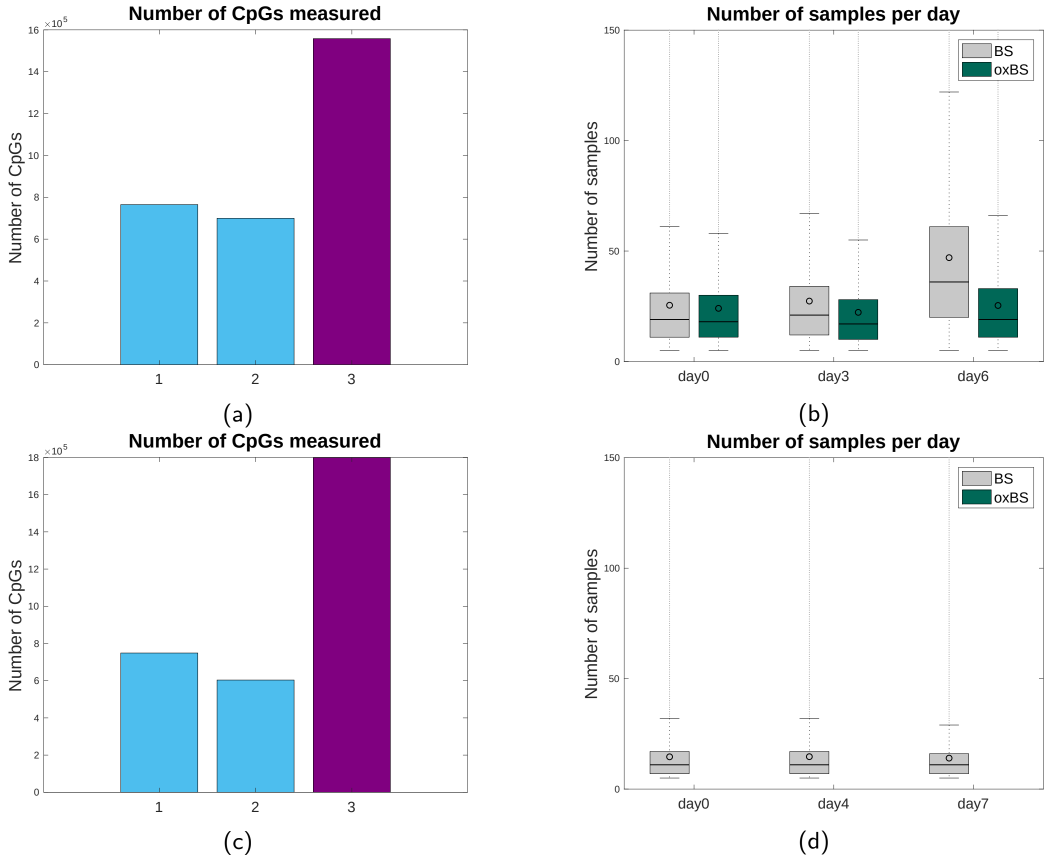


Figure 1: Number of CpGs with observations at one, two, or three days in WT (a) and Tet TKO (c). Average number of independent single CpG samples (sequencing depth) per day for BS and oxBS of WT (b) and for BS of Tet TKO (d) data.

We have double stranded single base pair resolution data from bisulfite (BS) and oxidative bisulfite sequencing (oxBS) for $3{,}022{,}903$ CpGs in wild type (WT) cells and for $3{,}151{,}985$ from BS data in Tet triple KO (Tet TKO) cells. In case of each of $1{,}464{,}801$ CpGs in WT and of $1{,}352{,}297$ in Tet TKO with only one or two observation time points available we predict for every measurement time point only the levels of the hidden states by performing a MLE for the (hydroxy-)methylation levels as described in Section 2.1 for estimating the initial distribution. In case of a CpG with three observation time points ($1{,}558{,}102$ in WT and $1{,}799{,}688$ in Tet TKO, see purple column in Figure 1a, 1c) we assume a linear behavior of the efficiencies over time and we analyse the HMM as described in Section 2.2 for estimating both the values of (hydroxy-)methylation efficiencies and levels over time. Using a computer cluster consisting of 32 machines with 16 physical kernels each, we are able to efficiently parallelize the computations for large bunches of all available CpGs.

Due to the low depth sequencing per time point and experiment ($40x$ for BS, $29x$ for oxBS in WT, and $14x$ coverage for BS in Tet TKO on average, see Figure 1b) we assume that the asymptotic properties of the MLE around the true parameter value do not hold [2, 3], especially in cases where the true parameter values are close to boundary constraints [4].

For that reason, we additionally use a Bayesian Inference (BI) approach to get the posterior distribution of the model parameters, i.e, the efficiencies over time. For all CpGs we choose as prior distribution the multivariate normal distribution $\mathcal{N}(\boldsymbol{\mu}, \Sigma)$, where the mean $\boldsymbol{\mu}$ is the average of the estimated efficiencies in [1]. Similarly, $\Sigma$ is the average of the corresponding covariance matrices. Note that in [1] MLE was sufficient due to the better coverage. Finally, we make a comparison between the MLE and the BI methods and we confirm that a BI method that incorporates an informative prior distribution should be preferable for epigenome-wide analysis especially for the regions where the coverage is low [5, 6].

##### Metropolis-Hastings

We apply BI by sampling from the multi-dimensional posterior $P(\mathbf{v}|\text{data}) = \frac{\mathcal{L}_2(\text{data}|\mathbf{v})P(\mathbf{v})}{\int_{\mathbf{v}} P(\text{data},\mathbf{v})}$ and avoid to approximate the normalizing factor $\int_{\mathbf{v}} P(\text{data}, \mathbf{v})$. Hence, we apply a Metropolis-Hastings MCMC approach using an asymmetric and truncated proposal distribution. The bounds of the truncation are determined s.t. the constraints for the efficiencies constantly hold for the time span of the observations, i.e., efficiencies are in $[0, 1]$ for all $t \in [0, t_{\max}]$. Hence, in every state $\mathbf{x} \in \mathbb{R}^v$ (with $v = 7$) of the MCMC we generate the next sample from a product of truncated univariate normals $\mathcal{N}(\mathbf{y}) = \prod_i f(\mathbf{y}_i|\mathbf{x}_i, \sigma_i^2/c, a_i, b_i)$, around the current MCMC point $\mathbf{x}$, where $\mathbf{x}_i$ refers to the $i-$th entry of the parameter vector for $i = 1, \ldots, 7$, $\sigma_i^2/c$ is the univariate normal variance and $a_i, b_i$ are the truncation bounds for parameter $\mathbf{x}_i$. Consider position $i$ where $\mathbf{y}_i$ refers to the gradient of an efficiency and $\mathbf{y}_{i-1}$ to the corresponding intercept. We sample the next value for each efficiency by sampling first the intercept $\mathbf{y}_{i-1}$ value from the truncated normal distribution within the interval $[a_{i-1}, b_{i-1}] = [0, 1]$ and based on this realization we sample the gradient $\mathbf{y}_i$ value from the truncated normal in $[a_i, b_i]$, where $a_i = -\mathbf{y}_{i-1}/t_{\max}, b_i = (1 - \mathbf{y}_{i-1})/t_{\max}$ as it is being illustrated in Figure 2. The bounds of probability $p$ are set as those of an intercept, i.e., $[a_i, b_i] = [0, 1]$.

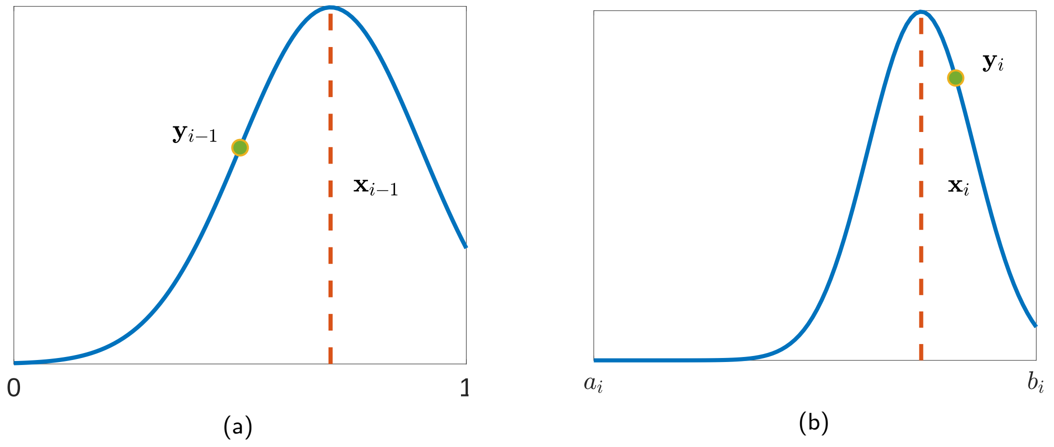


Figure 2: Metropolis Hasting's update step: We sample a new efficiency vector using two truncated normal distributions in two steps: (a) Step 1: We sample the intercept $\mathbf{y}_{i-1}$ from the truncated normal with mean $\mathbf{x}_{i-1}$ and bounds [0, 1]. (b) Step 2: We sample the gradient $\mathbf{y}_i$ from the truncated normal distribution with mean $\mathbf{x}_i$ and bounds $[a_i, b_i]$, which depend on the sampled intercept $\mathbf{y}_{i-1}$ of Step 1.

Note that the variance of parameter $\mathbf{x}_i$ we used for the proposal distribution is the same as the variance of the prior distribution $\sigma_i^2 = \Sigma_{i,i}$ normalized by a scale factor $c$. Since it is well known that the efficiency of Metropolis-Hastings algorithm crucially depends on the scaling of the proposal density, we empirically choose a $c = 50$ to normalize the standard deviation of the proposal distribution[1] s.t. the average MCMC acceptance ratio is around $25\%$ of the total number of generated samples [7] As final

[1]A low acceptance ratio indicates a wide proposal, while a high acceptance ratio indicates a narrow proposal and in both extreme cases the convergence is slow.

estimators of the BI method we get the sample mean of the posterior distribution and we build credible intervals using the corresponding sample covariance.

##### Fit of whole-genome data vs the model

Using box plots, we compare the levels of CC, TT and CT-TC CpG dyads for the whole genome present in the data of BS and oxBS in WT (Figure 3a, 3b) and of BS in Tet TKO (Figure 3c, 3d) and the probabilities of the observable states predicted by the two HMMs using MLE or BI for estimating the model's parameters. The circles inside the plots correspond the the mean value of each box plot and the horizontal lines to the medians. The bottom and the top of the boxes are the first and the third quartiles. The values for the whiskers correspond to the $\pm 2.7 \cdot s_{\mathrm{data}}$ interval from the sample mean, where $s_{\mathrm{data}}$ is the sample standard deviation of the data. To quantify the goodness of the fit for each estimation method we report in Table B the average Kullback-Leibler divergence $D_{KL}(P||Q) = \sum_i P(i) \ln \frac{P(i)}{Q(i)}$ between the data distribution $P$ and the distribution $Q$ predicted by the model. Note that the model fit to the data reported by the average Kullback-Leibler divergence metric is better for the MLE than for BI for both WT and Tet TKO data. This is to be expected since MLE always tries to maximize the likelihood of the data no matter how well the data samples represent the true underlying distribution.

In Figure 4 we plot the average efficiencies computed by the two estimation methods (MLE vs BI) at days 0,3,6 for WT and days 0,4,7 for Tet TKO. We average over all CpGs along the DNA for which we sampled at all three measurement time points. We observe that there are some major differences between the MLE and the BI estimates. First in WT the ML estimates show an evident decrease of maintenance over time while BI estimates show maintenance to be almost constant. In addition, the hydroxylation activity seems to slightly drop using MLE while BI estimates that it increases. In the Tet TKO experiment, the ML estimates give a completely unexpected increase of maintenance activity, while *de novo* seems to be not affected compared with its WT behavior. On the contrary, BI estimators for maintenance in Tet TKO remain almost unchanged comparing with their WT - BI behavior, while interestingly *de novo* seems to drop in a much slighter rate in the absence of Tet enzymes. Looking carefully at the prediction of the enzymatic activity we have several reasons to trust more the results of the BI method than those of MLE. In the WT data we observe that the BI estimates are in line with the genome wide behavior being described in the literature for the vast majority of the examined regions [8, 1]. Furthermore, the prediction of the remaining *de novo* activity being present mainly in the BI and not in the ML estimates for the Tet TKO data is in line with the detection of remaining nonCpG methylation in our RRHPoxBS data set which is not part of the model and therefore presents an independent readout of Dnmt3a and 3b activity. In addition, looking at the box plots we note that the dispersion of the efficiencies values is evidently smaller for the BI estimates. This shows the higher precision of the BI estimates for the efficiencies comparing with the ML estimates.

To quantify the improvement of BI compared to MLE regarding the decrease in the uncertainty of the parameter estimators we computed the average hypervolume corresponding to the covariance matrices of the estimators in each case. The volume of the hyper-ellipse of a multivariate-normal distribution is proportional to the square root of the generalized variance, i.e., the square root of the determinant of the covariance matrix, and it is given by the function

$$V = \frac{2\pi^{v/2}}{v\Gamma(v/2)} (\chi^2_{crit})^{v/2} |\Sigma_{\mathbf{v}^*}|^{1/2},$$

where $v$ is the number of parameters, $|\Sigma_{\mathbf{v}^*}|$ is the determinant of the estimators' covariance matrix, $\chi^2_{crit}$ is the critical value for $\chi^2(v)$ and $\Gamma(x)$ is the gamma function (see Figure 5 for details). In WT the average volume of the hyper-ellipse in case of MLE is $0.0024$ while the average hyper-ellipse volume in BI is $3.5162 \cdot 10^{-5}$. In Tet TKO the average volume of the hyper-ellipse for ML estimates is $0.0480$ while in case of BI only $9.6 \cdot 10^{-4}$. In Figure 6 we plot the levels of the hidden states of the HMM for each combination of statistical estimation method (MLE vs BI) and cell type (WT vs Tet TKO). Overall we see small differences on the prediction of the hidden states even though there is some evident difference

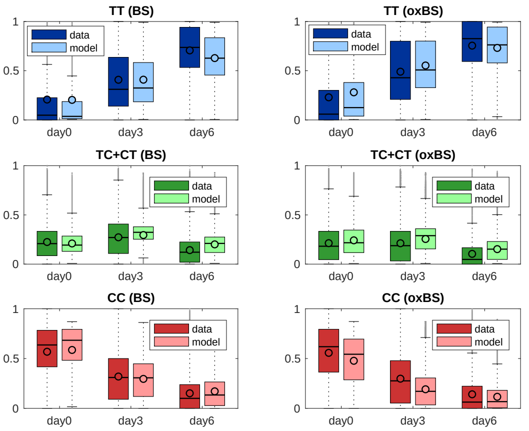


(a) WT - MLE

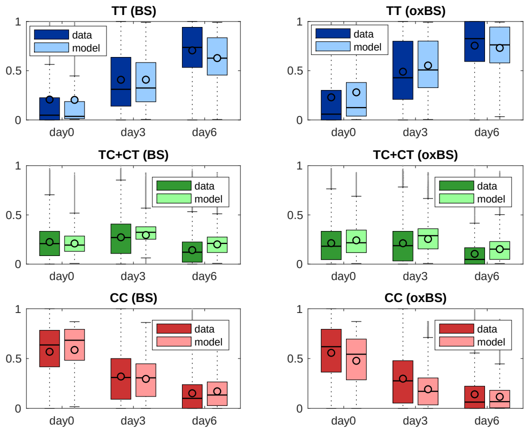


(b) WT - BI

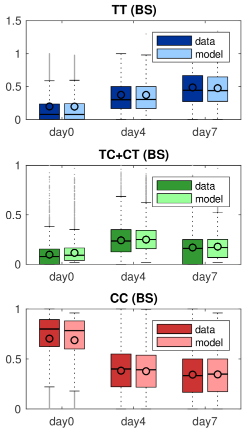


(c) Tet TKO - MLE

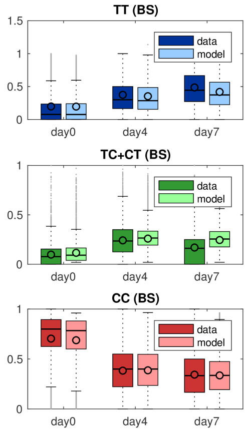


(d) Tet TKO - BI

Figure 3: Comparison between data and prediction of observable states after fitting the HMMs based on MLE (a), (c) and BI (b), (d). Dark box plots show the experimentally measured frequencies states and light box plots correspond to the values predicted by the two HMMs.

in the enzyme's efficiency estimators in particular for the Tet TKO case. This indicates again how critical an ML estimation bias can be for an accurate estimation of the efficiencies. For all the aforementioned reasons we use the BI estimates as the output of our model for all the analysis we present in the main manuscript as well as for the clustering that we describe in the sequel.

Table B: Computed Kullback-Leibler divergence between the data and the model distribution for MLE and BI, where $P_{bs}$ and $P_{ox}$ is the data distribution for BS and oxBS experiment respectively.

| experiment - method | $\hat{D}_{KL}(P_{bs}\|\|\pi_{bs})$ | $\hat{D}_{KL}(P_{ox}\|\|\pi_{ox})$ |
|---|---|---|
| WT - MLE | 0.1802 | 0.2369 |
| WT - BI | 0.2904 | 0.3941 |
| Tet TKO - MLE | 0.154 | - |
| Tet TKO - BI | 0.277 | - |

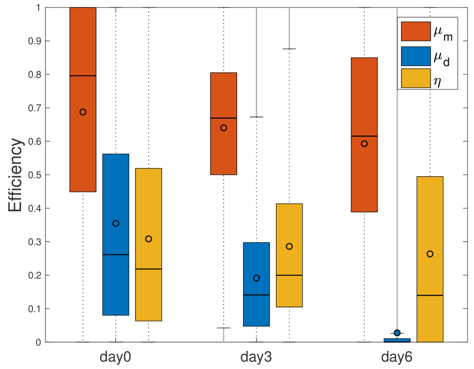


(a) WT - MLE

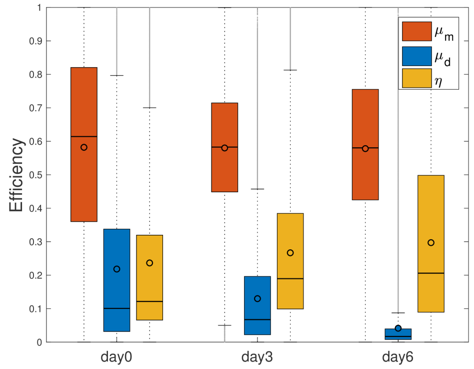


(b) WT - BI

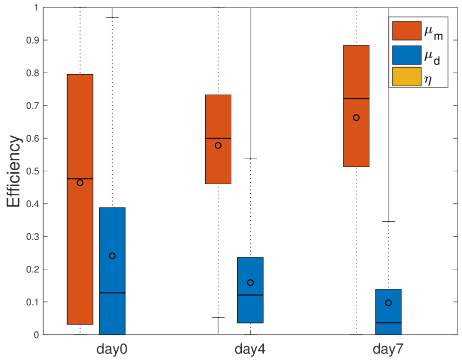


(c) Tet TKO - MLE

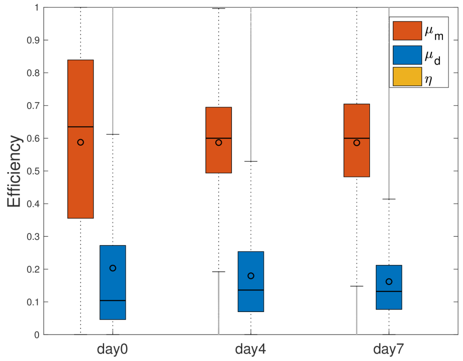


(d) Tet TKO - BI

Figure 4: Bar plots for maintenance, *de novo* and hydroxylation efficiencies over time taken by MLE (a), (c) and BI (b), (d) methods. Red = maintenance methylation efficiency ($\mu_m$), blue = *de novo* methylation efficiency ($\mu_d$), yellow = hydroxylation efficiency ($\eta$).

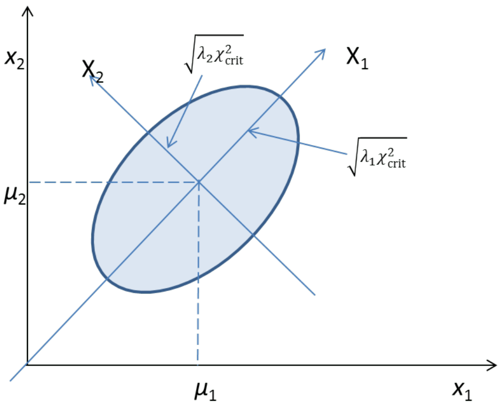


Figure 5: The ellipse has axes pointing in the directions of the eigenvectors $X_1, X_2, ..., X_p$ of the covariance matrix $\Sigma$. Here, for the bivariate normal, the longest axis of the ellipse points in the direction of the first eigenvector $X_1$ and the shorter axis is perpendicular to the first, pointing in the direction of the second eigenvector $X_2$. The half length of the axis corresponding to eigenvector $X_i$ is given by the formula $l_i = \sqrt{\lambda_i \chi^2_{crit}}$.

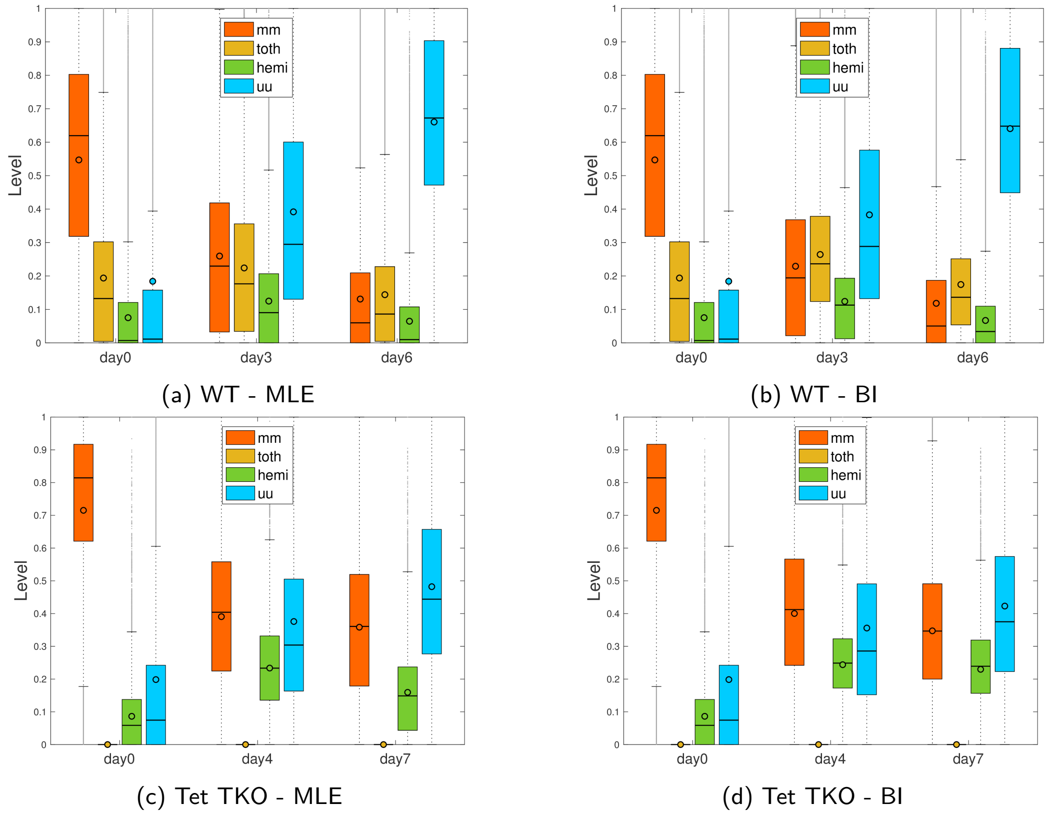


(a) WT - MLE

(b) WT - BI

(c) Tet TKO - MLE

(d) Tet TKO - BI

Figure 6: Bar plots for the hidden states levels for all CpGs in the genome estimating the parameters with MLE (a), (c) and BI (b), (d). Red = symmetric methylated CpG (mm - 5mC/5mC), yellow = 5hmC in all possible combinations (toth - 5hmC/C, C/5hmC, 5hmC/5mC, 5mC/5hmC, 5hmC/5hmC), green = hemi methylated CpGs (hemi - 5mC/C or C/5mC), blue = unmethylated CpGs (C/C).

### 4 Clustering

#### 4.1 $k$-means clustering

After computing the estimates for the efficiency functions over time for $1,558,102$ CpGs in WT we applied a $k$-means clustering of those functions to identify CpGs with similar efficiencies. For each cluster total number of clusters $k = \{1, \ldots, 10\}$ we produce $100$ initializations in order to avoid that the algorithm converges in a local optimum. In order to determine the optimal number of clusters we use three different criteria as explained in Section 4.3. According to all the three metrics we get $k_{opt} = 4$ as the optimal number of clusters. In Figure 7 we see the optimal $k$-means clustering. In clusters 2, 3, and 4 the average maintenance methylation efficiency slightly increases over time which contradicts the decreasing concentrations of the H3K9me2 and Uhrf1 [2] in 2i, especially assuming it happens on a genome wide level [8]. Hence, we have little confidence in this clustering result and we therefore developed a more sophisticated clustering approach that takes into account not only the estimated parameter's vectors but also their standard deviations [9]. Incorporating uncertainty around data points in the clustering process can generally produce different and closer to the "ground" truth clusters when there is a measurement or an estimation error. This is being illustrated in Figure 8. The "optimal" number of clusters that are being formed by the $k$-error algorithm is two according to all criteria of section 4.3 used.

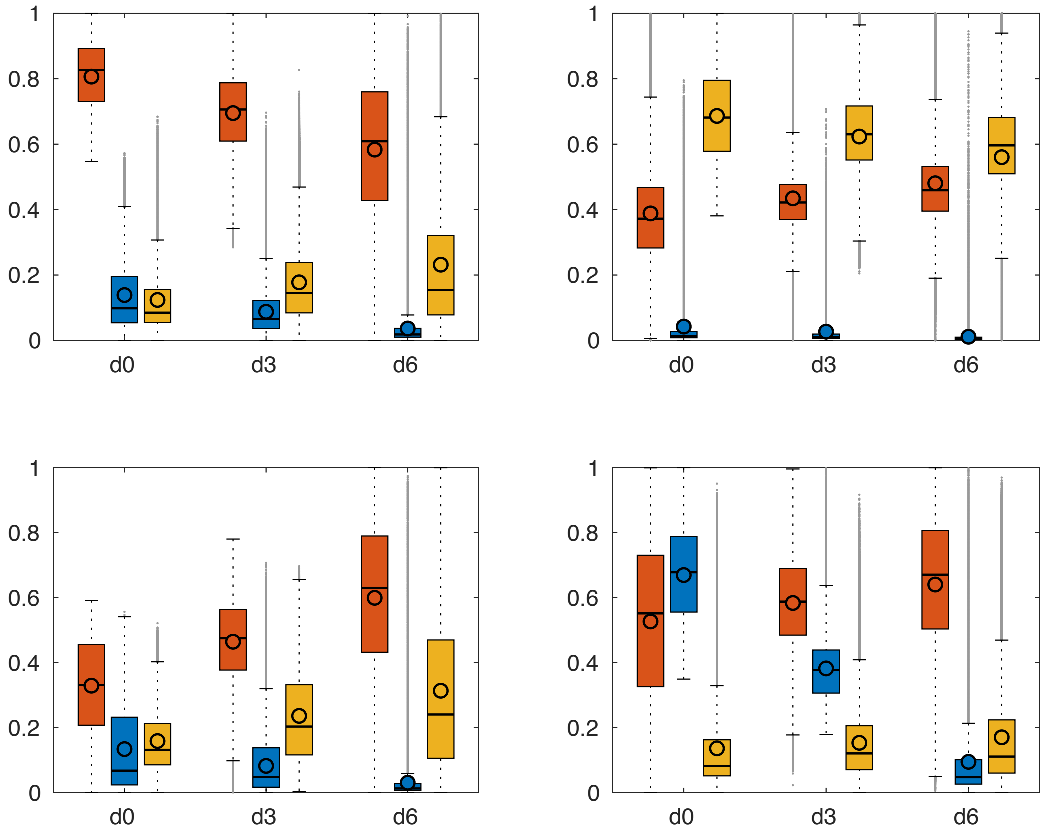


Figure 7: The optimal clustering of enzymatic efficiencies over time based on the k-means algorithm and the squared euclidean distance. Red = maintenance methylation efficiency ($\mu_m$), blue = *de novo* methylation efficiency ($\mu_d$), yellow = hydroxylation efficiency ($\eta$), y-axis = efficiency, x-axis = samples.

[2]the main cofactor of Dnmt1

Table C: Number of CpGs per cluster

| Cluster Name | Number of CpGs |
|---|---|
| Cluster 1 | 628108 |
| Cluster 2 | 257065 |
| Cluster 4 | 289119 |
| Cluster 5 | 261205 |

Table C lists the number of CpGs for each cluster. The majority of CpGs are assigned to cluster 1 (628108 CpGs), while cluster 2 (257065 CpGs), cluster 3 (289119 CpGs) and cluster 4 (261205 CpGs) are of similar size.

#### 4.2 $k$-error clustering

We use a modification of the $k$-means algorithm called $k$-error clustering [9] that takes into account the uncertainties of each data point, i.e, the covariance matrix $\Sigma_{\mathbf{v}}$ of the parameter vector of the efficiencies $\mathbf{v}$.

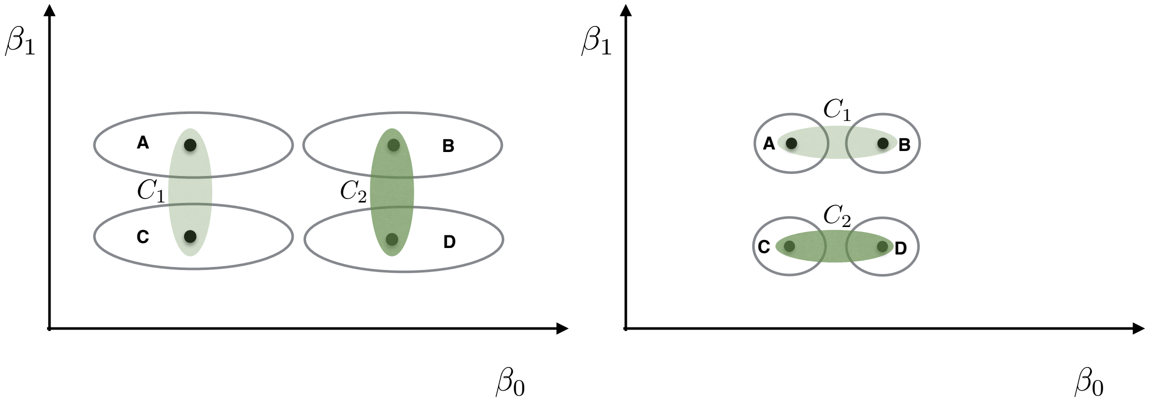


Figure 8: Illustration of the clustering of an estimated enzymatic efficiency (with intercept $\beta_0$ and gradient $\beta_1$) for CpGs A, B, C, D using $k$-means clustering (Left) vs $k$-error clustering (Right).

If $\mathbf{v}_1, \ldots, \mathbf{v}_N \in \mathbb{R}^v$ are the estimated parameter vectors and $\Sigma_1, \ldots, \Sigma_N \in \mathbb{R}^{v \times v}$ the associated covariance matrices for all input CpGs. If we assume that the estimated parameter vectors are independent and each arises from a $v-$variate normal distribution with one of $k$ possible means $\theta_1, \ldots, \theta_k$, that is $\mathbf{v}_i \sim N_p(\mu_i, \Sigma_i)$, where $\mu_i \in \{\theta_1, \ldots, \theta_k\}$ for $i = 1, \ldots, N$. We seek to find the clusters $C_1, \ldots, C_k$ such that the parameter vectors that have the same mean $\mu_i = \theta_j$ all belong to the same cluster $C_j$, for $j = 1 \ldots, k$.

Let $S_j = \{i \mid \mathbf{v}_i \in C_j\}$, hence $\mu_i = \theta_j$ for $j = 1 \ldots k$ and $\forall i \in S_j$. Given $N$ parameter vectors $\mathbf{v} = (\mathbf{v}_1, \ldots, \mathbf{v}_N)$ and their error matrices $\Sigma_1, \ldots, \Sigma_N$ we search for a partition $S = (S_1, \ldots, S_k)$ and $\theta = (\theta_1, \ldots, \theta_N)$ that maximizes the following likelihood:

$$\mathcal{L}_c(\mathbf{v}) = \prod_{j=1}^{k}\prod_{i\in S_j}\frac{1}{2\pi}^{p/2}|\Sigma_i|^{-1/2}e^{-1/2(\mathbf{v}_i-\theta_j)\Sigma_i^{-1}(\mathbf{v}_i-\theta_j)^\intercal}, \tag{4}$$

where $|\Sigma_i|$ is the determinant of matrix $\Sigma_i$ for $i = 1, \dots, N$. Maximizing the likelihood of Eq. 4 is equivalent as minimizing the total squared Mahalanobis distance of the points that belong to a cluster from the cluster centroid [9], i.e.,

$$\min_S \sum_{j=1}^{k}\sum_{i\in S_j}(\mathbf{v}_i - \hat{\theta}_j)\Sigma_i^{-1}(\mathbf{v}_i - \hat{\theta}_j),$$

where $\hat{\theta}_j$ is the ML estimate of $\theta_j$ given by

$$\hat{\theta}_j = \Big(\sum_{i\in S_j}\Sigma_i^{-1}\Big)^{-1}\Big(\sum_{i\in S_j}\mathbf{v}_i\Sigma_i^{-1}\Big) \tag{5}$$

for $j = 1, \dots, k$. Note that the estimated centroid $\hat{\theta}_j$ is a weighted mean of the point in cluster $C_j$. We refer to it as the Mahalanobis mean of $C_j$.

In addition, by using simple matrix algebra we can compute that the covariance matrix $\Psi_j$ associated with the centroid $\hat{\theta}_j$ equals

$$\Psi_j = \mathrm{Cov}(\hat{\theta}_\mathrm{j}) = \mathrm{Cov}\left(\left(\sum_{\mathrm{i}\in \mathrm{S_j}}\Sigma_\mathrm{i}^{-1}\right)^{-1}\left(\sum_{\mathrm{i}\in \mathrm{S_j}}\mathbf{v}_\mathrm{i}\Sigma_\mathrm{i}^{-1}\right)\right) = \left(\sum_{\mathrm{i}\in \mathrm{S_j}}\Sigma_\mathrm{i}^{-1}\right)^{-1}.$$

Hence, after randomly choosing an initial set of $k$ centroids (Forgy method) the $k$-error method follows as an iteration over the next two steps until no change happens to the assignment of the points.

a. Assign each data point $x_i$ to the cluster whose centroid is the closest using the squared Mahalanobis distance, i.e,

$$\arg\min_j \mathrm{d}_{i,j} = \arg\min_j (x_i - \hat{\theta}_j)\Sigma_i^{-1}(x_i - \theta_j)^\intercal. \tag{6}$$

b. For clusters $C_1, \dots, C_k$ compute the new cluster centroids $\hat{\theta}_1, \dots, \hat{\theta}_k$ as the Mahalanobis means of the clusters (Eq. 5).

The choice of the distance function used in Eq. 6 guarantees the decrease in the objective function in each iteration of $k$-error as it is shown in [9].

Table D: Number of CpGs per cluster

| Cluster Name | Number of CpGs |
|---|---|
| Cluster 1 | 792703 |
| Cluster 2 | 642794 |

Using the $k$-error clustering approach on the estimates derived from WT ES cells, CpGs were grouped into 2 distinct clusters. Cluster 1, containing 792703 CpGs, is defined by high and stable maintenance

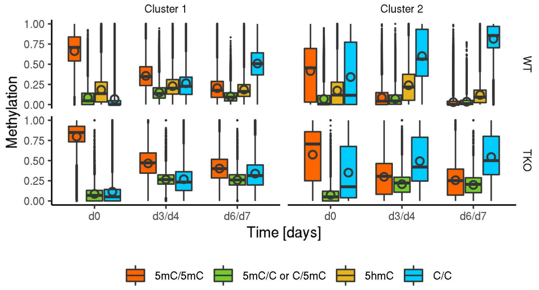


(a) DNA Methylation

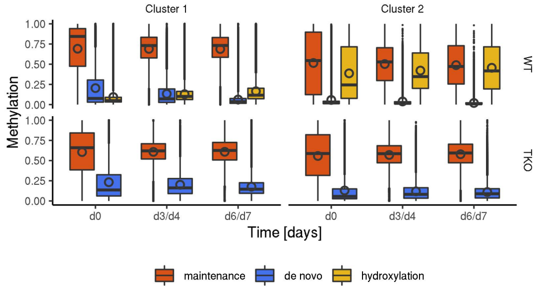


(b) Enzyme Efficiencies

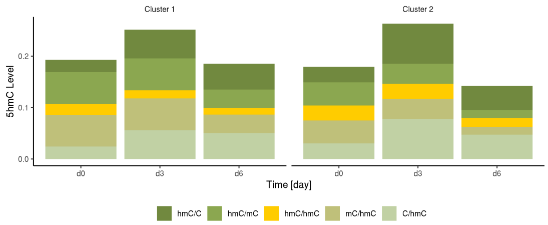


(c) Average 5hmC Distribution

Figure 9: Number of CpGs with observations at one, two, or three days in WT (a) and Tet TKO (c). Average number of independent single CpG samples (sequencing depth) per day for BS and oxBS of WT (b) and for BS of Tet TKO (d) data.

methylation efficiency, high but strongly decreasing *de novo* methylation efficiency, as well as low hydroxylation efficiency. Consequently, CpGs within cluster 1 exhibit high methylation level. Cluster 2 contains 642794 CpGs and is characterized by a strong hydroxylation efficiency, as well as intermediate but stable maintenance methylation efficiency and low *de novo* methylation efficiency.

#### 4.3 Metrics for deciding the number of clusters

In order to identify the "optimal" number of clusters we use Davies-Bouldin and Calinski-Harabasz criteria. These metrics evaluate the overall within-to-between cluster variability, each in a slightly different fashion. In addition, we use the simple but commonly used elbow method that considers the sum of squared errors (SSE) of a certain clustering. The goal of this method is to identify the number of clusters after which adding more clusters results only to a minor decrease of the SSE.

##### Davies-Bouldin Criterion

Let $R_{i,j}$ be the within-to-between cluster distance ratio for clusters $i$ and $j$ defined as

$$R_{i,j} = \frac{S_i + S_j}{M_{i,j}},$$

where $S_i$ is a measure of within cluster $i$ variance, i.e.,

$$S_i = \frac{1}{|C_i|} \sum_{x \in C_i} \mathrm{d}(x, m_i)$$

and $M_{i,j} = \mathrm{d}(m_i, m_j)$ is a measure of separation between clusters $i$ and $j$ defined as the distance between the clusters' centroids $m_i, m_j$. We define $D_i = \max_{j \neq i} R_{i,j}$, i.e., the $R_{i,j}$ of the most similar cluster to cluster $i$, and we get Davies-Bouldin index as the average over all $D_i$ indices,

$$DB = \frac{1}{N}\sum_{i=1}^{N} D_i.$$

Since the value of $DB$ represents the (worst-case) average within-to-between cluster distance ratio we decide the optimal number of clusters to be the one that provides the smallest $DB$.

##### Calinski-Harabasz Criterion

The Calinski-Harabasz criterion, alternatively called Variance Ratio Criterion (VRC), is defined as

$$CH_k = \frac{SS_B}{SS_W}\frac{(N-k)}{k-1},$$

where $SS_B$ is the overall between-cluster variance, $SS_W$ is the overall within-cluster variance, $k$ is the number of clusters and $N$ is the total number of observations. The overall between-cluster variance is defined as

$$SS_B = \sum_{i=1}^{k} |C_i| \, \mathrm{d}(m_i, m)$$

where $m_i$ is the centroid of cluster $i$ and $m$ is the overall sample mean. The overall within-cluster variance is defined as

$$SS_W = \sum_{i=1}^{k} \sum_{x \in C_i} \mathrm{d}(x, m_i),$$

where the second sum goes over all points $x$ that belong to cluster $C_i$. Intuitively, clusterings with well defined clusters have a large $SS_B$ and a small $SS_W$. Hence, the larger the $CH_k$ for varying $k$, the better the clustering. Consequently, to determine the optimal number of clusters we target to maximize $CH_k$ w.r.t. $k$.

##### Elbow method

We compute the sum of squared errors (SSE) for a range of number of clusters $k$. We choose the optimal $k$ to be the point where the graph starts to flatten significantly. In Figure 10 the optimal number of clusters is clearly two.

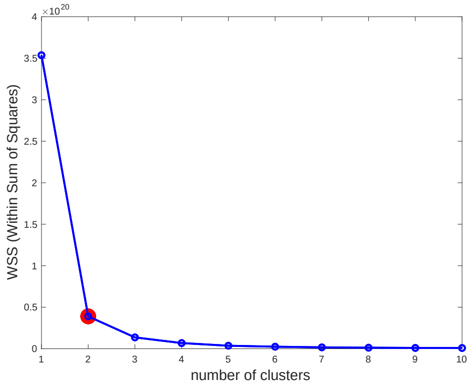


Figure 10: Elbow method: The "optimal" number of clusters is the point where the graph starts to smooth out, i.e., the "elbow" of the graph.

###### Choice of distance function of the metrics

For the evaluation of the clusterings we plug in as the distance function of the above criteria the same distance function that we used for performing the clustering. Hence, in case of $k$-means we use the square euclidean distance $\mathrm{d}(x,y) = \|x-y\|^2$ while for $k$-error we use the squared Mahalanobis distance $\mathrm{d}(x,y) = (x-y)^\intercal \Sigma_x^{-1}(x-y)$, where $\Sigma_x$ is the covariance matrix of point $x$.

### 5 Correlation of Enzyme Efficiencies and Methylation Levels

#### 5.1 Spatial Correlations of Enzyme Efficiencies

Let $X_s$ be the discrete space random process describing the dispersion of an enzymatic activity over the whole genome at a certain time point. For a space interval $\tau$ its spatial autocorrelation is defined as

$$R(\tau) = \frac{\mathbb{E}[(X_s - \mu_{X_s})(X_{s+\tau} - \mu_{X_{s+\tau}})]}{\sigma_{X_s}\sigma_{X_{s+\tau}}}.$$

Similarly the spatial cross-correlation between two random processes $X, Y$ that describe the dispersion of two different enzymatic activities over the genome is defined as

$$\rho_{X,Y} = \frac{\mathbb{E}[(X_s - \mu_{X_s})(Y_{s+\tau} - \mu_{Y_{s+\tau}})]}{\sigma_{X_s}\sigma_{Y_{s+\tau}}}.$$

We compute the sample spatial autocorrelation $\hat{R}$ and the cross-correlations $\hat{\rho}$ for all enzymatic processes in both WT and Tet TKO experiments as follows. Let genome position $s \in S(\tau)$ when both CpGs of positions $s$ and $s+\tau$ are included in our data. Then

$$\hat{R}(\tau) = \frac{1}{|S(\tau)-1|\hat{\sigma}_{X_s}\hat{\sigma}_{X_{s+\tau}}} \sum_{s\in S(\tau)} (X_s - \bar{X}_s)(X_{s+\tau} - \bar{X}_{s+\tau})].$$

In the above sample estimator $\bar{X}_s$ and $\hat{\sigma_{X_s}}$ are the sample mean and the sample standard deviation respectively of all measurements $X_s$ for which $s \in S(\tau)$. The same way we compute

$$\hat{\rho}(\tau) = \frac{1}{|S(\tau)-1|\hat{\sigma}_{X_s}\hat{\sigma}_{Y_{s+\tau}}} \sum_{s\in S(\tau)} (X_s - \bar{X}_s)(Y_{s+\tau} - \bar{Y}_{s+\tau})].$$

Fixing $\tau = 5$ we plot in Figure 11 the sample autocorrelations and sample cross-correlations between all efficiencies at all time points in WT (Figure 11a, 11c, 11e) and Tet TKO (Figure 11b, 11d, 11f) experiments. Together with the sample correlations we report $95\%$ confidence intervals following the approach of [10] and p-values for the null hypothesis that the auto or the cross-correlation is zero.

We observe a negative correlation of methylation and hydroxylation efficiencies across the entire genome and a positive correlation between *de novo* and maintenance efficiencies. Furthermore, we observe for all efficiencies a positive autocorrelation which constantly declines as the distance between CpGs increases. In WT maintenance autocorrelation flattens out at about 1500bp distance between CpGs and at the same time it starts losing its significance (p>0.01). Both *de novo* and hydroxylation activity show a high autocorrelation in the beginning (more than 0.5) and seem to influence windows of larger sizes, around 2000 bp.

In Tet TKO cells maintenance methylation activity seems to flatten out earlier than in WT, around 300 bp, which is in agreement with the observation that maintenance activity appears misregulated in Tet TKOs, in particular showing an increase at the TSS. On the contrary, the activity area of *de novo* enzymes seems to be stable compared to WT cells (around 2000 bp). Overall the spatial autocorrelations do not indicate any change of the activity window size of the enzymes over time.

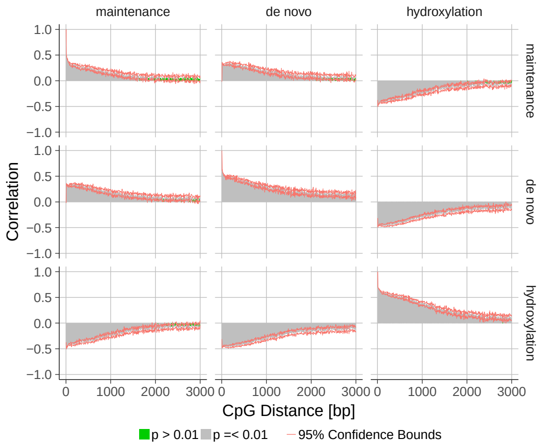


(a) WT - day 0

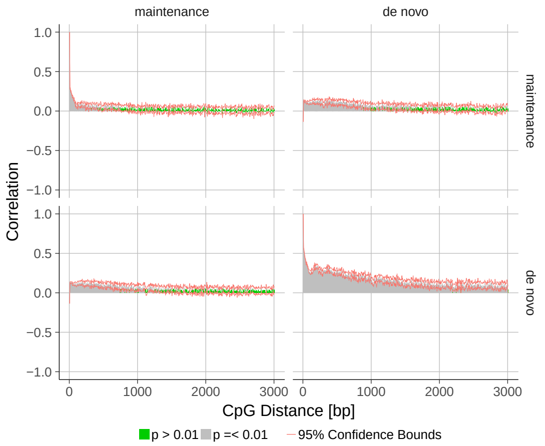


(b) Tet TKO - day 0

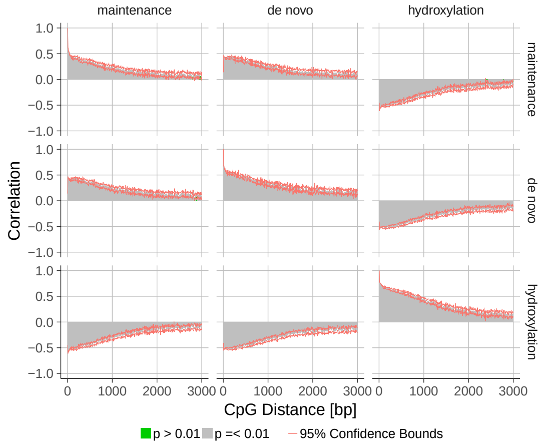


(c) WT - day 3

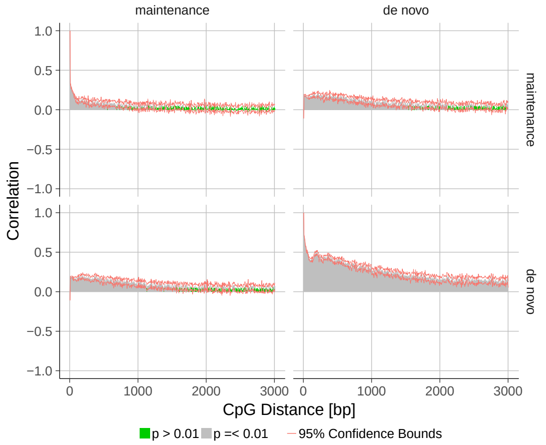


(d) Tet TKO - day 4

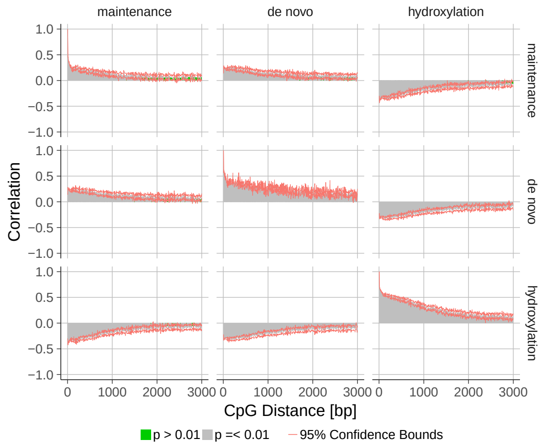


(e) WT - day 6

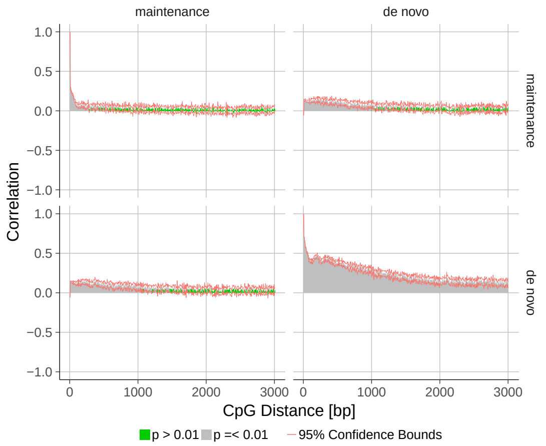


(f) Tet TKO - day 7

Figure 11: Spatial auto- and cross correlation of maintenance, *de novo*- and hydroxylation efficiency across the genome. Grey bars indicate correlations with a p value $< 0.01$, green bars correlations with p values $> 0.01$, red line shows the confidence bounds. Y-axis displays correlation, x-axis gives the distance of CpG in base pairs.

#### 5.2 Pearson Correlation of Enzyme Efficiencies and methylation Level

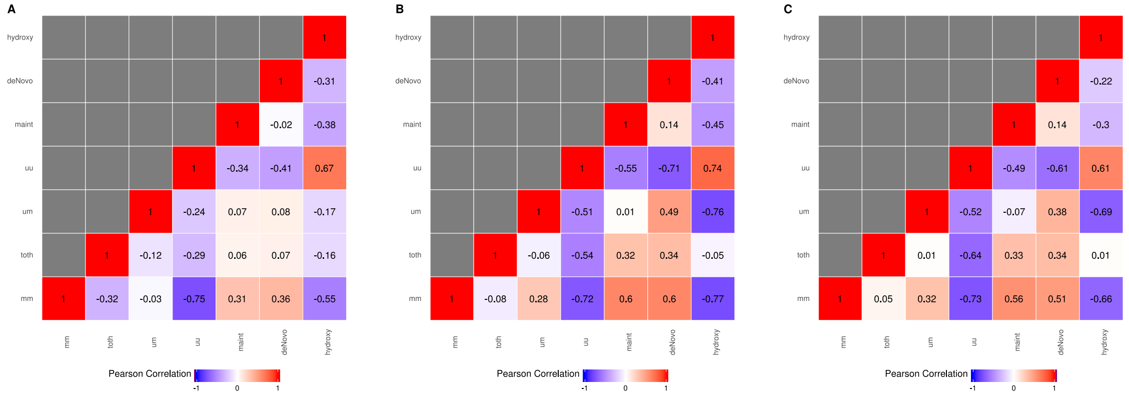


Figure 12: Pearson correlation of enzyme efficiencies and methylation level in WT ES cells for d0 (A), d3 (B) and d6 (C). mm = fully methylated (5mC/5mC), toth = hydroxylated CpG of all possible states, um = hemimethylated (5mC/C or C/5mC), uu = unmethylated (C/C), maint = maintenance methylation efficiency, deNovo = *de novo* methylation efficiency, hydroxy = hydroxylation efficiency

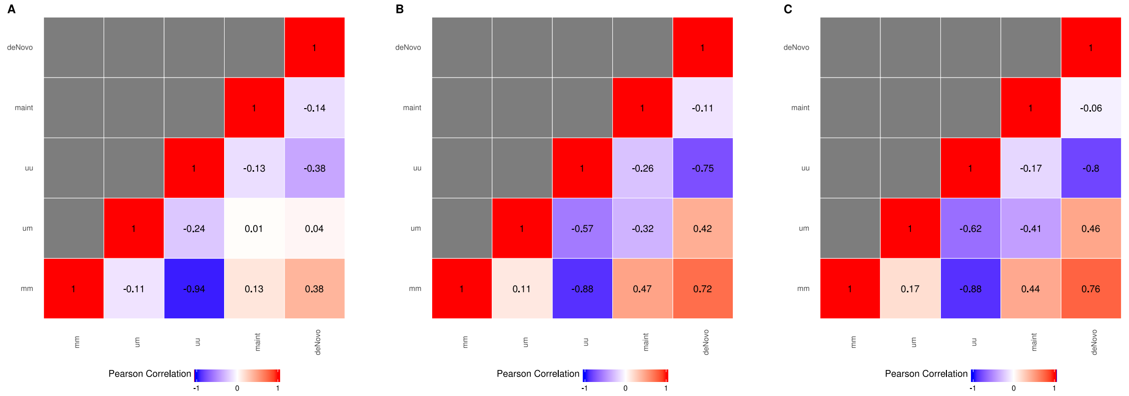


Figure 13: Pearson correlation of enzyme efficiencies and methylation level in Tet TKO ES cell for d0 (A), d4 (B) and d7 (C). mm = fully methylated (5mC/5mC), um = hemimethylated (5mC/C or C/5mC), uu = unmethylated (C/C), maint = maintenance methylation efficiency, deNovo = *de novo* methylation efficiency.

In addition to the spatial correlation, we calculated a simple pearson correlation for average efficiency and estimated modification levels.

In WT ES cells, we observe a positive correlation of fully methylated CpGs with *de novo* and maintenance efficiency at day 0. For later time points, this correlation increases. In addition, there is a positive correlation between the two methylation efficiencies and hemimethylated CpGs for day 3 and day 6, which is not present at day 0. This is likely due to an increase in hemimethylated CpGs which temporally overlap with the remaining Dnmt activity in 2i.

Interestingly, we observe no correlation between hydroxylated CpGs and hydroxylation efficiency. Instead, hydroxylation activity correlates with unmethylated CpGs. Thus, we conclude that high hydroxylation activity is not sufficient to generate stable 5hmC but will rather result in methylation free CpGs.

In Tet TKO, the correlation between maintenance methylation efficiency and fully methylated CpG dyads are reduced and in case of day 0 only comes to 0.13. However, we observe a stronger correlation of fully methylated CpGs with *de novo* methylation efficiency which points towards a misregulated methylation activity in the absence of Tet enzymes.

### 6 Spike-In analysis

To determine the conversion rate of BS and oxBS we included short oligonucleotides into our RRHPoxBS libraries. The oligo mix is part of the TrueMethyl kit from Cambridge Epigenetix and includes C, 5mC, 5hmC and 5fC at known positions. After sequencing, we calculated the conversion rates for each cytosine variant, which were than included into our model to compensate for conversion errors.

Table E: Spike-In oligo-nucleotides. M = 5mC, H = 5hmC, F = 5fC

| SpikeIn Oligo Name | DNA Sequence 5' >3' |
|---|---|
| Q1hmC | TAMGATCAMGGCGAATMCGATMGAATCAMAGTGGCGMTTTAHGAAGTGCGAMAGCMTTAG |
| Q3hmC | TAMGATCAMGGCGAATMHGATMGAATCMTTGTAGCGMTTTAHGAAGTGCGAMAGCHTTAG |
| Q6hmC | TAMGATCAHGGCGAATGHHGATMGAATCAGTMAAGCGMTTTAHGAAGTGHGAMAGCHTTAG |
| QfC | TACGATCAFGGCGAATCCGATCGAATCGTTTMGGCGCTTTACGAAGTGCGACAGCCTTAG |
| QmC | TAMGATMAMGGMGAATMMGATMGAATMTAGMTTGMGMTTTAMGAAGTGMGAMAGMMTTAG |
| SQC | TACGATCACGGCGAATCCGATCGAATCMAGATMGCGCTTTACGAAGTGCGACAGCCTTAG |

Table F: Conversion rate of cytosine variants included in the TruMethyl Spike in after BS treatment

| | C | 5mC | 5hmC | 5fC |
|---|---|---|---|---|
| Serum | 0.996332 | 0.0699681 | 0.0673588 | 0.75626 |
| 72h-2i | 0.996165 | 0.0725858 | 0.0715434 | 0.762992 |
| 144h-2i | 0.995809 | 0.0696952 | 0.0682802 | 0.739254 |

Table G: Conversion rate of cytosine variants included in the TruMethyl Spike in after oxBS treatment

| | C | 5mC | 5hmC | 5fC |
|---|---|---|---|---|
| Serum | 0.99687 | 0.0662679 | 0.964215 | 0.968836 |
| 72h-2i | 0.99656 | 0.0670022 | 0.967298 | 0.9663 |
| 144h-2i | 0.996901 | 0.0534113 | 0.949588 | 0.932044 |

### 7 Hemimethylated CpGs

Hemimethylated CpGs are the result of *de novo* methylation events and/or active and passive demethylation. Theoretically, selective methylation of a DNA strand could provide a strand specific gene regulation mechanism. Thus, we analysed the strand specific methylation of genes transcribed from plus- and minus strand under primed conditions (d0). Results are displayed in Figure 14.

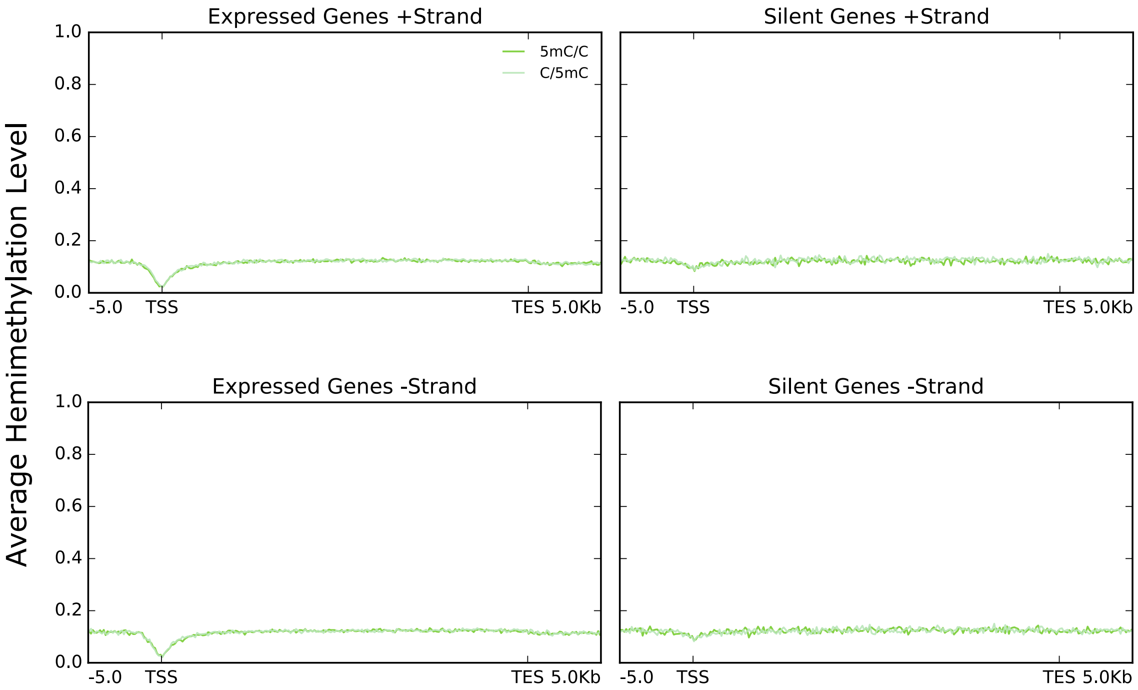


Figure 14: Average hemimethylated CpGs detected by RRHPoxBS across expressed and not/low expressed genes. Dark green = 5mC/C, light green = C/5mC

We cannot observe any methylation differences between genes expressed from upper or lower DNA strands. In both cases we observe the same amount of hemimethylation at both strands. The same holds true for low/not expressed genes.

### 8 Demethylation Kinetics

Previous studies indicated that Tet TKO cells exhibit the same demethylation kinetics as WT ES cells during their transition from Serum to 2i [8]. However, our RRHPoxBS data shows a noticeable difference in the methylation levels of WT and Tet TKO cells. Thus, we calculated the demethylation rate $r_{\mathrm{dem}}$ for each cell type to further investigate the distinct demethylation kinetics. For this, we calculated the increase of unmethylated cytosines for time points and for WT $t = \{3, 6\}$ and Tet TKO cells for time points $t = \{4, 7\}$ using the equation:

$$r_{\mathrm{dem}}(t) = \frac{\mathrm{TT}(t) - \mathrm{TT}(0)}{t}$$

Results are displayed in Figure 15.

In contrast to the previous observations, we observe distinct demethylation rates for WT and Tet TKO cells. WT cells exhibit a demethylation rate between 6 and 8%, whereas Tet TKO ES cells show a reduced demethylation rate of around 4% (Fig. 15 (a)). Consequently, the demethylation rate in Tet TKO cells w.r.t. to WT cells is reduced by 30 to 50%, demonstrating the considerable contribution of Tet enzymes to DNA demethylation (Fig. 15 (b)).

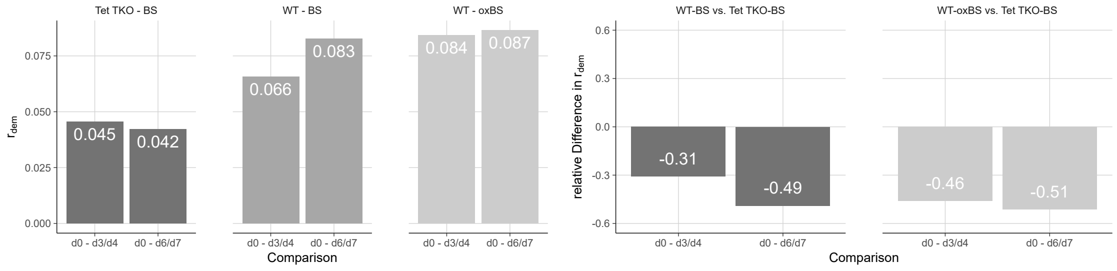


(a) (b)

Figure 15: (a) Demethylation rate in WT and Tet TKO cells (b) Relative difference in demethylation rate between WT and Tet TKO cells.

### 9 nonCpG Methylation

Frequently, DNA methylation occurs outside of a CpG context [11, 12]. Hence, we determined the sequence occurrence of nonCpG methylation in our WT samples. For our analysis, we considered only nonCpG positions which are (i) methylated above the conversion error, (ii) show at least three methylated reads and (iii) a coverage of $\geq 10$. In accordance with literature, we find that CpA is the most common methylated sequence after CpG on both DNA strands (Figure 16). Interestingly, our sequence analysis suggests, that the most frequent sequence context for nonCpG methylation at the plus strand is ApCpA (Figure 16). Furthermore, we see that the majority of all nonCpG in our data set correspond to FMRs and PMDs, whereas only a small fraction can be found in LMRs and UMRs (Figure 18). This observation nicely matches our model's prediction according to which FMRs and PMDs exhibit higher *de novo* methylation activity (main manuscript figure 9) mainly caused by Dnmt3a and 3b.

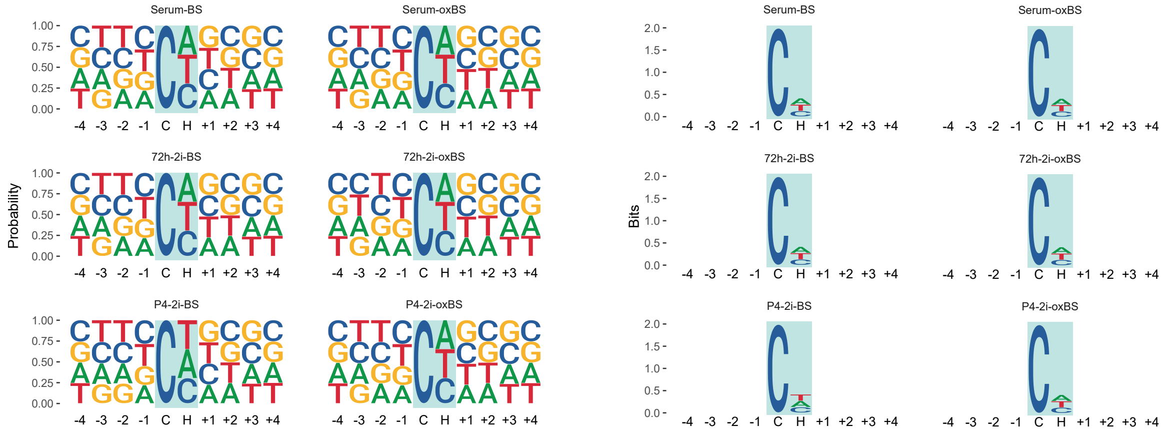


Figure 16: Occurrences of nonCpG methylation in Serum and 2i cultivated WT ES cells for the plus strand. Size of bases indicate the probability at a given position. nonCpG with 4 bases up- and downstream are shown.

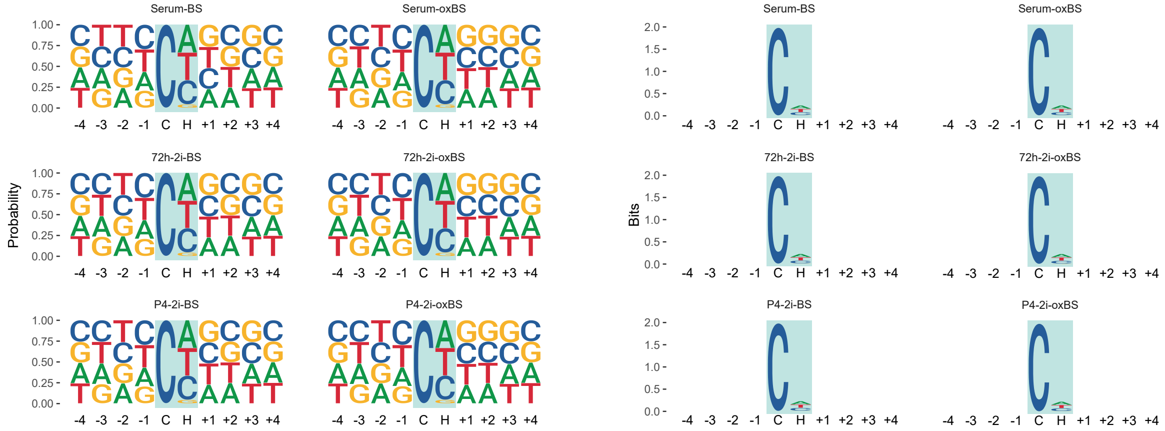


Figure 17: Occurrences of nonCpG methylation in Serum and 2i cultivated WT ES cells for the plus strand. Size of bases indicate the probability at a given position. CpG with 4bp up- and downstream are shown.

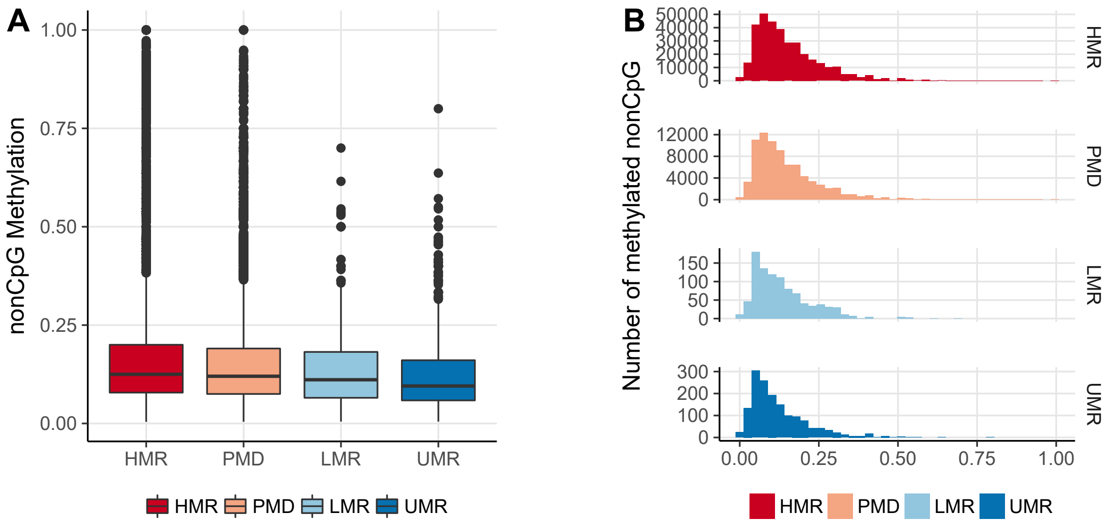


Figure 18: Methylation level and distribution of methylated nonCpG in HMRs, PMDs LMRs and UMRs. Methylation level (A). nonCpG methylation distribution in HMRs, PMDs, LMRs and UMRs (B)

.

### 10 Methylation Level and Efficiencies across Chromosomes

In this section we provide the input-data information plots as well as the output of our model for each of the 21 main chromosomes of the ES cells. In Figure 19, 20 we plot the number of CpGs for each chromosome with one, two or three observation days in WT and Tet TKO cells, respectively. We plot the average number of samples (sequencing depth) for each chromosome in WT (Figure 21) and Tet TKO (Figure 22). In Figure 23, 24 we show the efficiencies over time computed by BI and in Figure 25, 26 we report the prediction of the model for the hidden states probabilities in each chromosome in WT and Tet TKO cells.

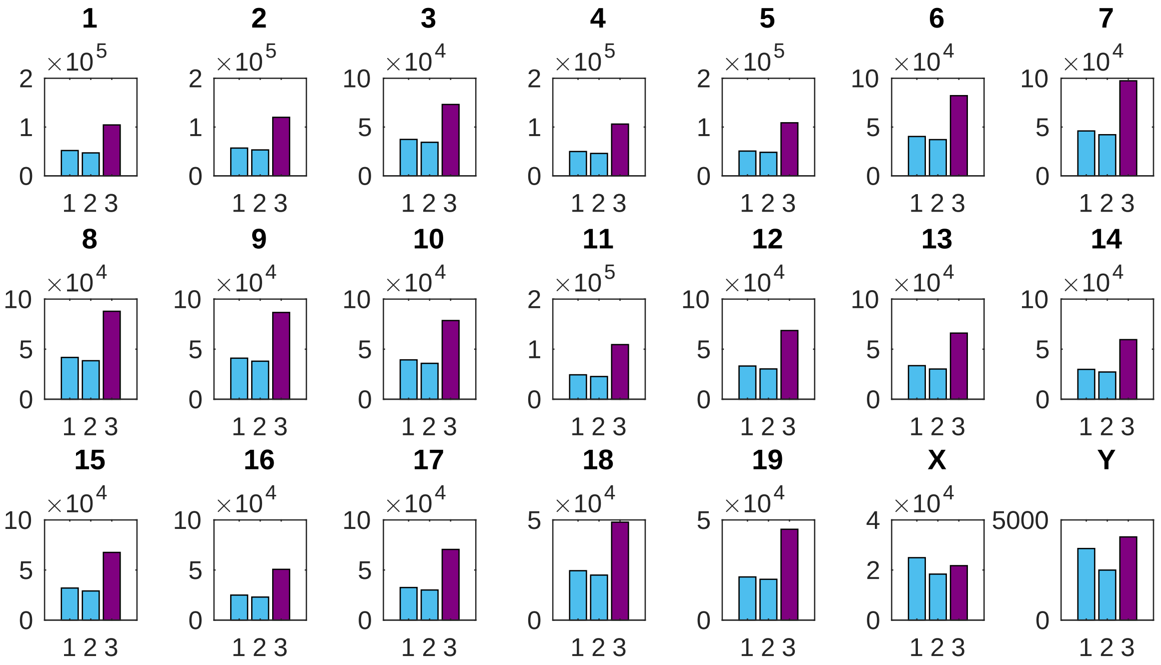


Figure 19: Number of CpGs (y-axis) with one, two or three observation days (x-axis) for each chromosome in WT data.

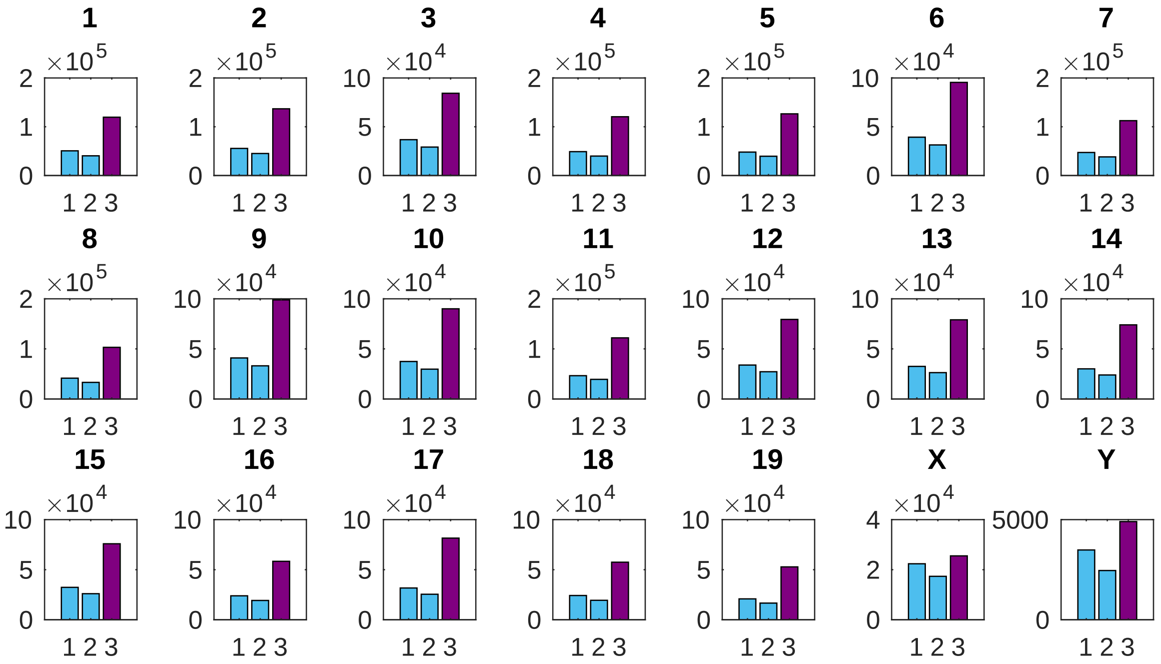


Figure 20: Number of CpGs (y-axis) with one, two or three observation days (x-axis) for each chromosome in Tet TKO data.

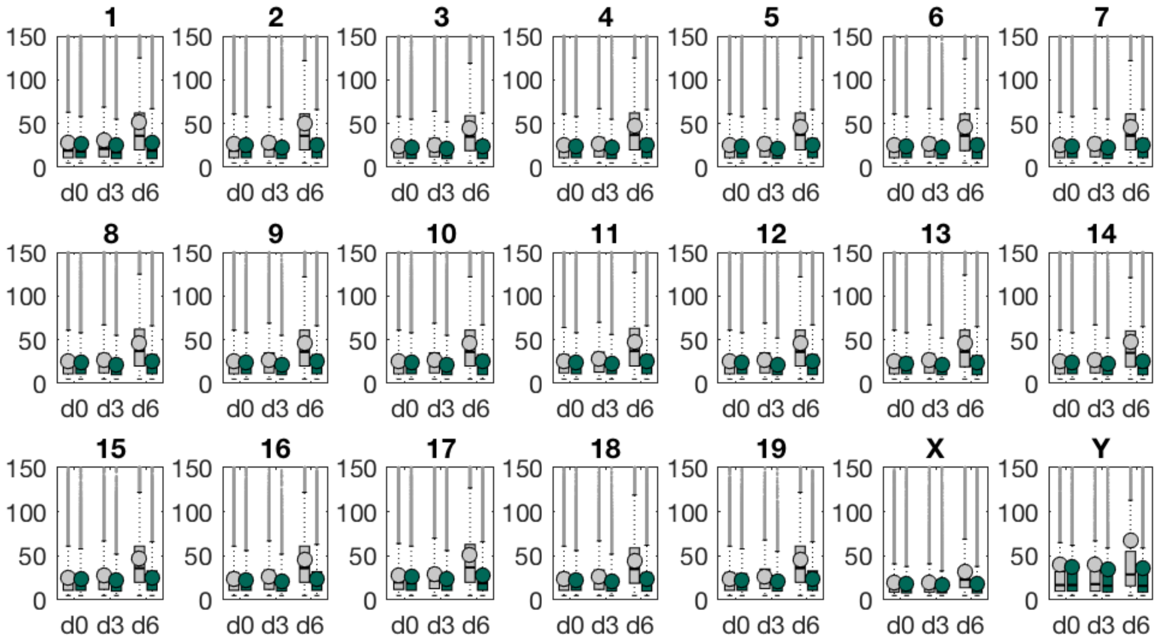


Figure 21: Average number of single CpG independent samples, i.e, depth sequencing, (y-axis) per day (x-axis) for each chromosome in WT data.

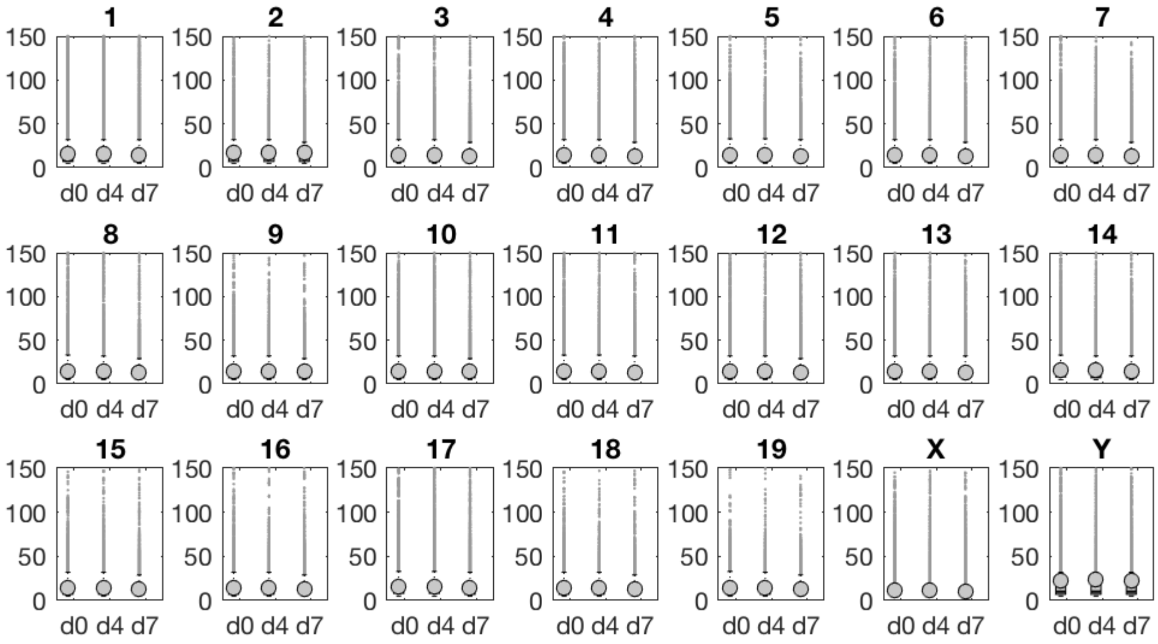


Figure 22: Average number of single CpG independent samples, i.e, depth sequencing, (y-axis) per day (x-axis) for each chromosome in Tet TKO data.

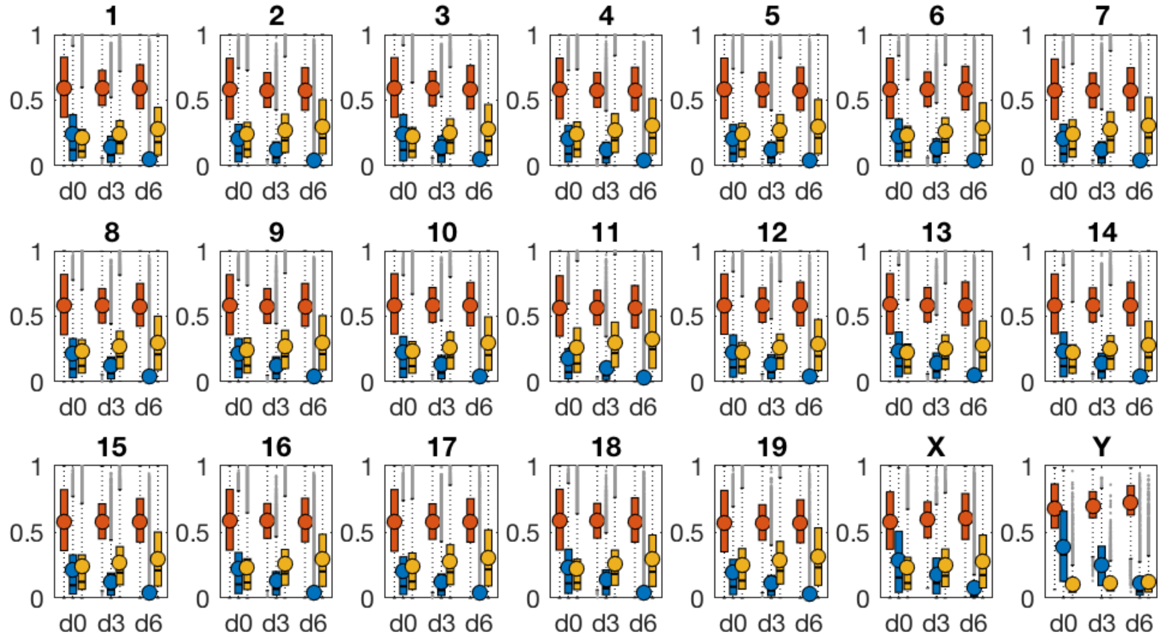

Figure 23: Bar plots for the maintenance (red), *de novo* (blue) and hydroxylation (yellow) efficiencies over time taken by BI method for each individual chromosome in WT cells.

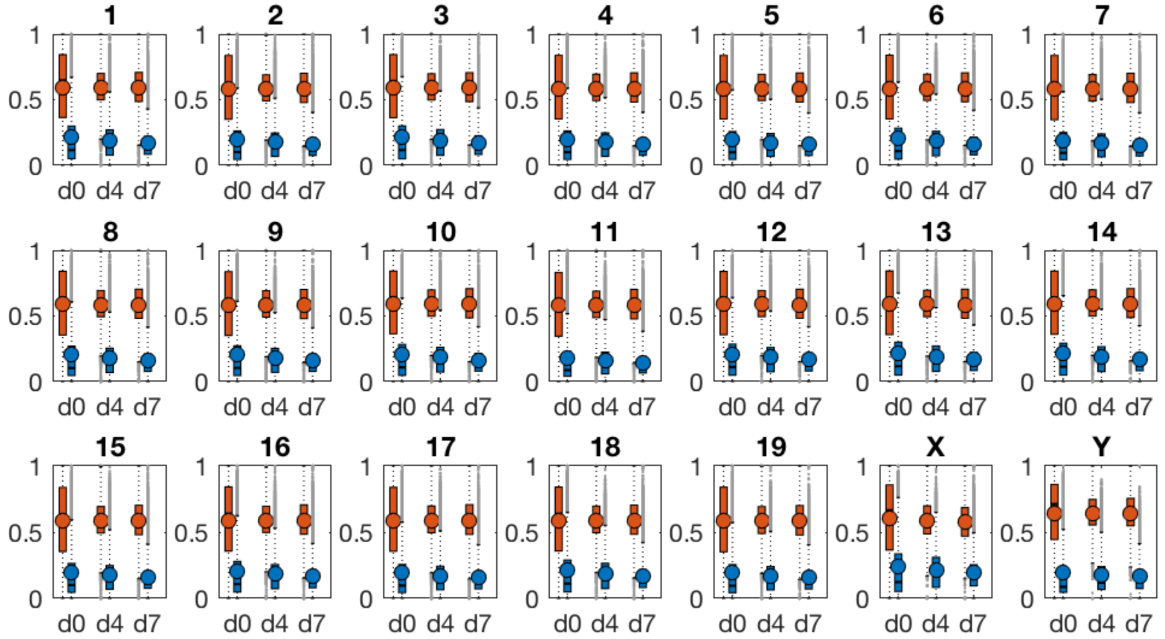

Figure 24: Bar plots for the maintenance (red) and *de novo* (blue) efficiencies over time taken by BI method for each individual chromosome in Tet TKO cells.

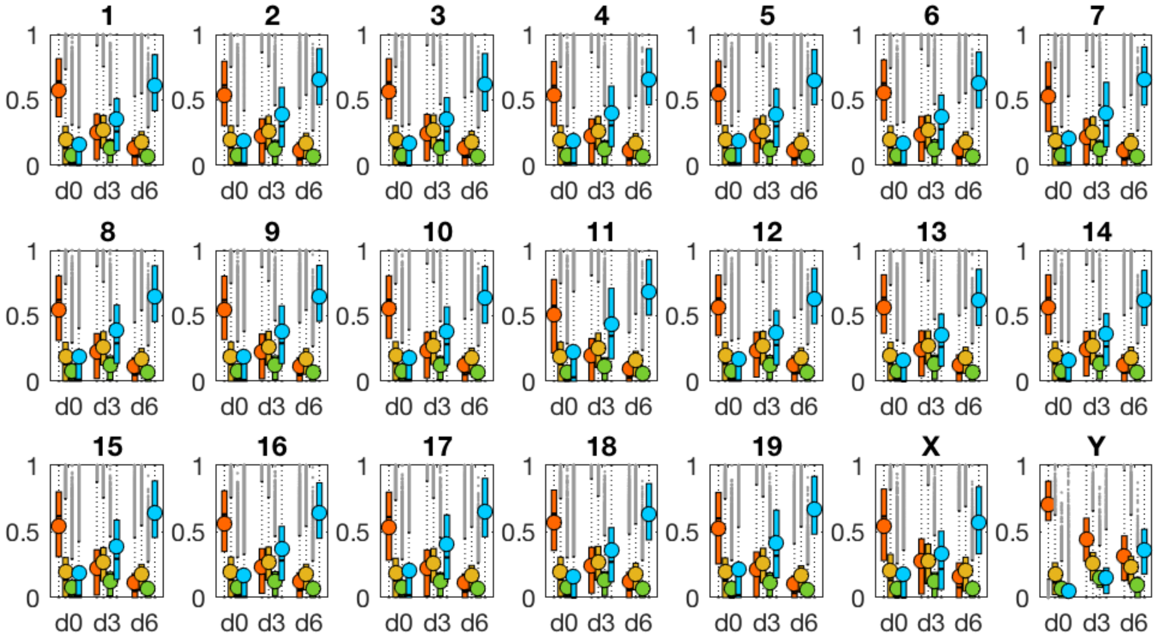


Figure 25: Bar plots for the hidden states levels over time of each individual chromosome in WT. Red = symmetric methylated CpG (mm - 5mC/5mC), yellow = 5hmC in all possible combinations (toth - 5hmC/C, C/5hmC, 5hmC/5mC, 5mC/5hmC, 5hmC/5hmC), green = hemi methylated CpGs (hemi - 5mC/C or C/5mC), blue = unmethylated CpGs (C/C).

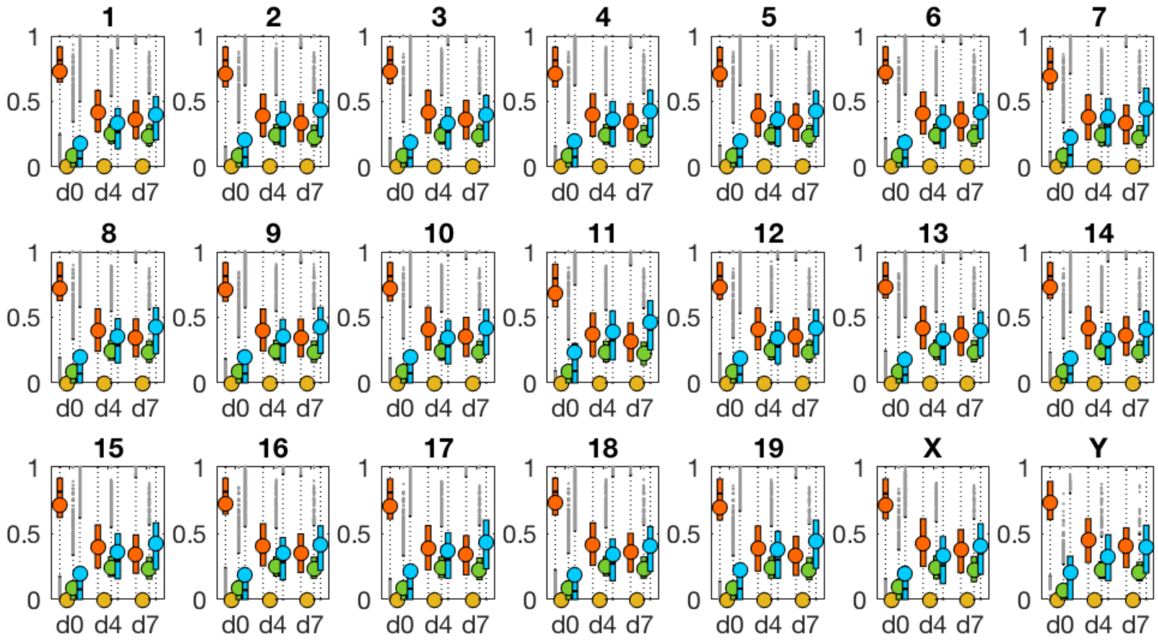


Figure 26: Bar plots for the hidden states levels over time of each individual chromosome in TET TKO. Red = symmetric methylated CpG (mm - 5mC/5mC), yellow = 5hmC in all possible combinations (toth - 5hmC/C, C/5hmC, 5hmC/5mC, 5mC/5hmC, 5hmC/5hmC), green = hemi methylated CpGs (hemi - 5mC/C or C/5mC), blue = unmethylated CpGs (C/C).

#### 11 Repetitive Elements

The majority of the mammalian genome is composed of repetitive elements (REs). Thus, we examined whether a subset of REs would reflect the average behavior of the genome. For this, we assign CpGs to individual REs. Figures 27 to 31 show methylation level and efficiencies for the 25 most frequent repetitive elements in our data set for WT and Tet triple TKO ES cells. Indeed, we observe that the majority of REs resemble closely the level and efficiency profile of individual chromosomes as well as the average genome profiles. However, we also observe some exceptions. Intracisternal A particle and major satellites exhibit considerable higher methylation level and methylation efficiency compared to the mean genome profile. In addition, GC rich elements show almost no 5mC/5hmC, low methylation efficiency but high hydroxylation activity of Tets. Thus, they resemble more the behavior of promoters and TSS.
In case of the Tet TKO cells, we observe, that the maintenance efficiency in the distinct repetitive elements converge. In addition, *de novo* methylation appears again reduced for the first time point, but remains present even after continuous incubation in 2i media (Fig.31).

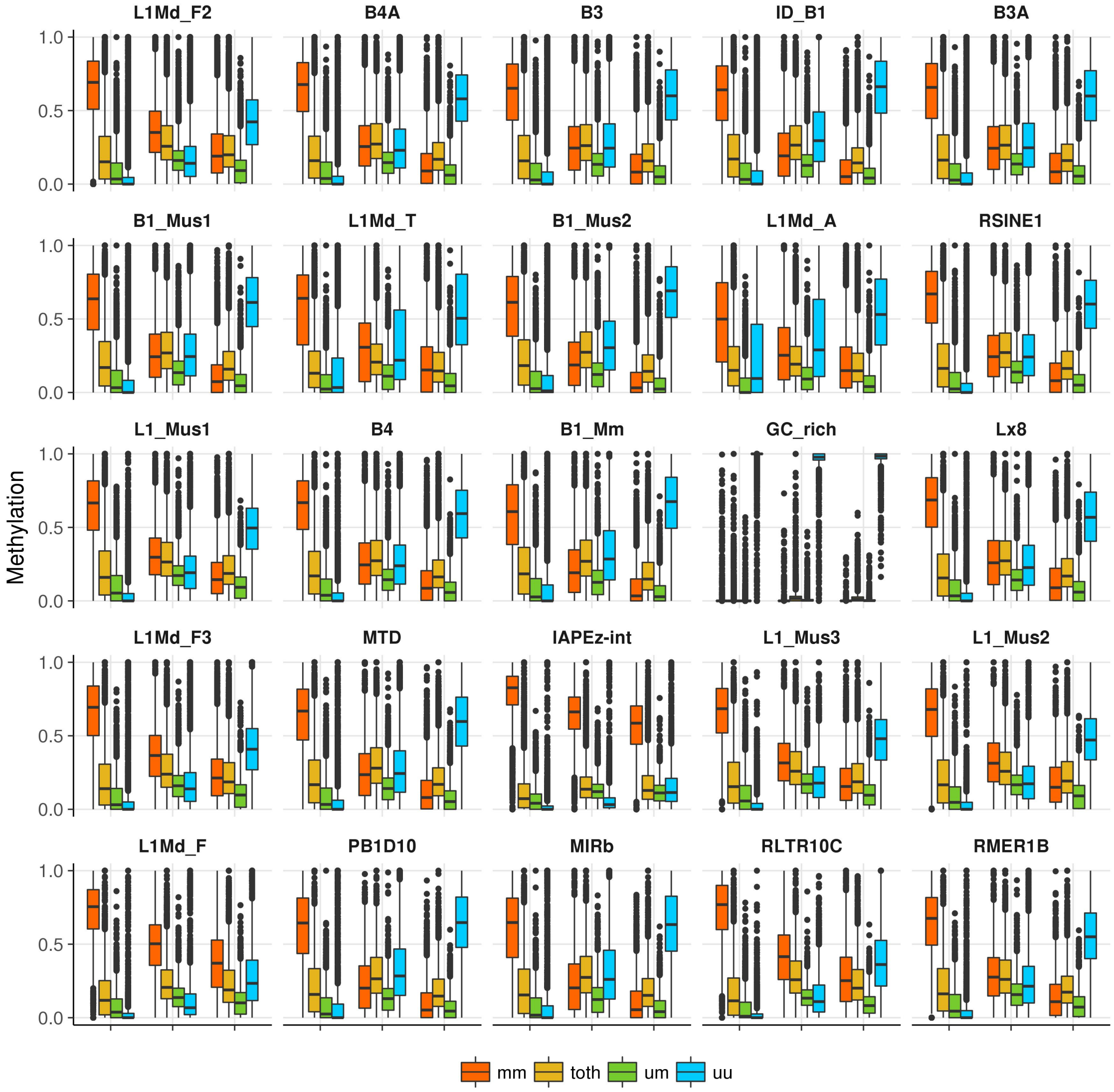


Figure 27: Methylation level at the 25 most frequent repetitive elements in our analysis for WT ES cells. Elements are presented in decreasing order, most frequent left top, least frequent right bottom. Annotation according to UCSC. y-axis = methylation frequency, x-axis = time in days (d0, d3, d6). Red = fully methylated CpGs (5mC/5mC), green = hemimethylated CpGs (5mC/C or C/5mC), yellow = 5hmC, blue = unmethylated CpGs (C/C).

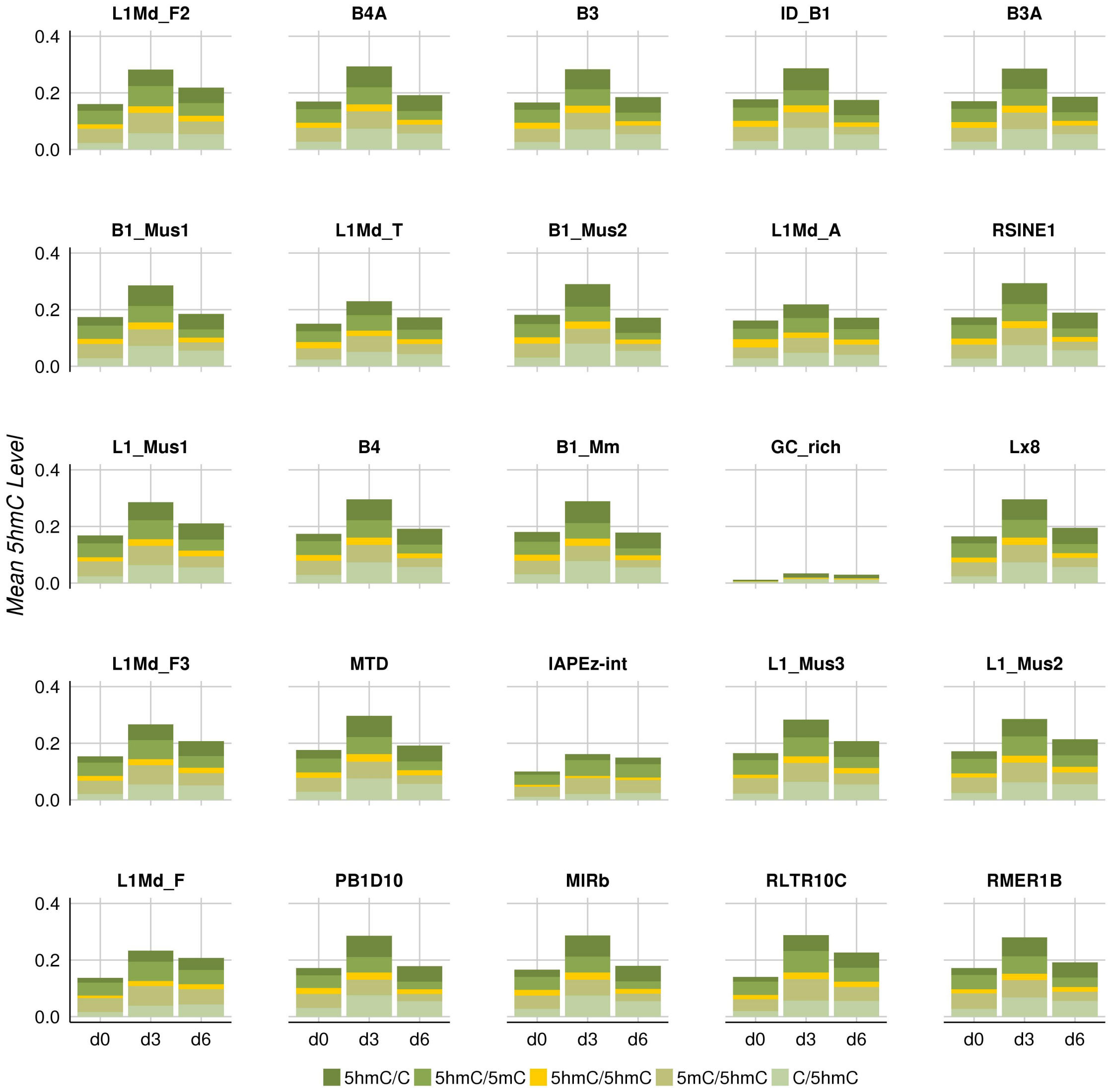


Figure 28: Level and distribution of 5hmC within the 25 most frequent repetitive elements in our data set for WT ES cells. Elements are presented in decreasing order, most frequent left top, least frequent right bottom. Annotation according to UCSC. y-axis = mean 5hmC level, x-axis = time in days (d0, d3, d6).

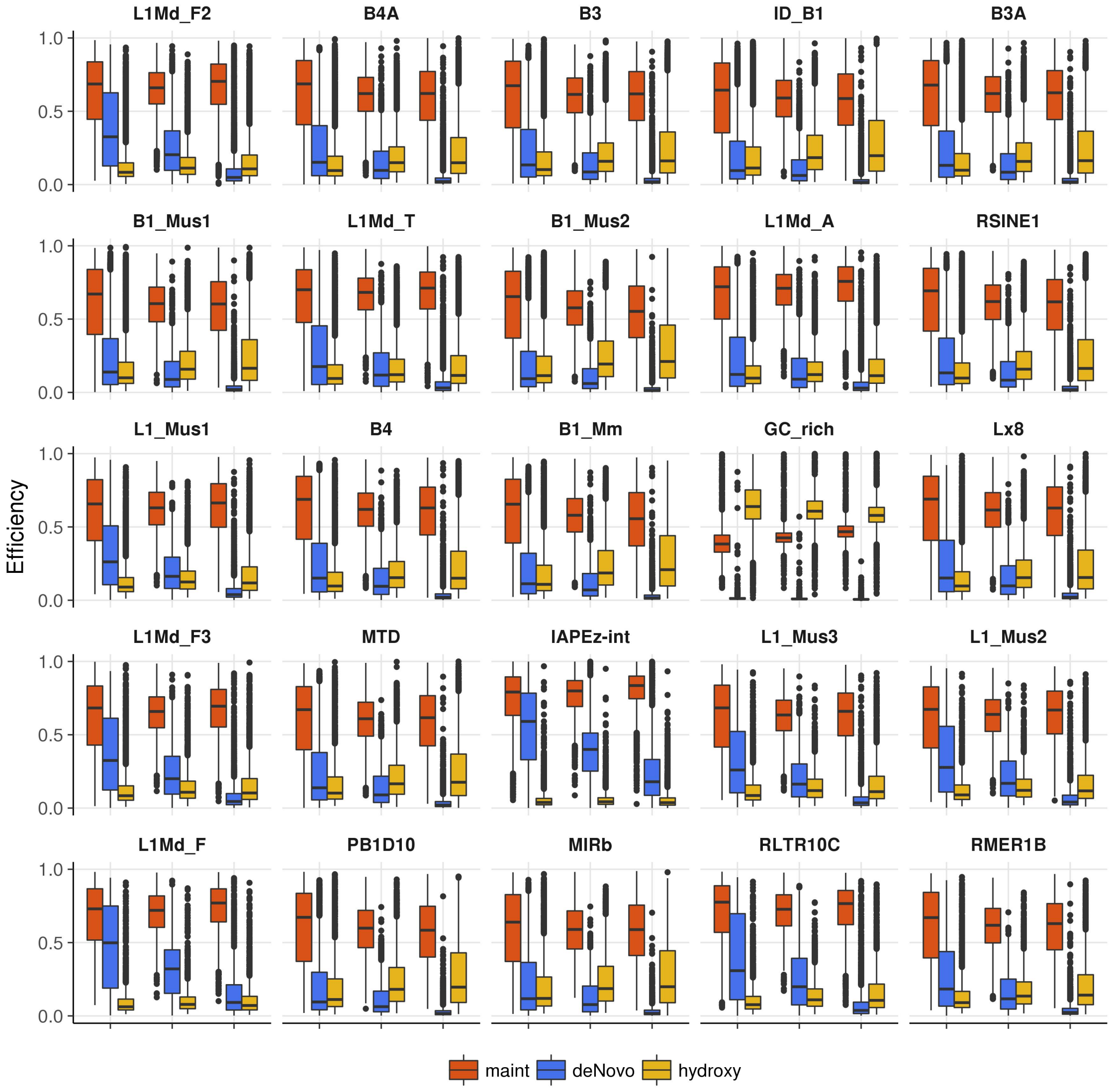


Figure 29: Efficiency profiles of the 25 most frequent repetitive elements in our analysis for WT ES cells. Elements are presented in decreasing order, most frequent left top, least frequent right bottom. Annotation according to UCSC. y-axis = efficiency; x-axis = time in days (d0, d3, d6), red = maintenance efficiency, blue = *de novo* efficiency, yellow = hydroxylation efficiency.

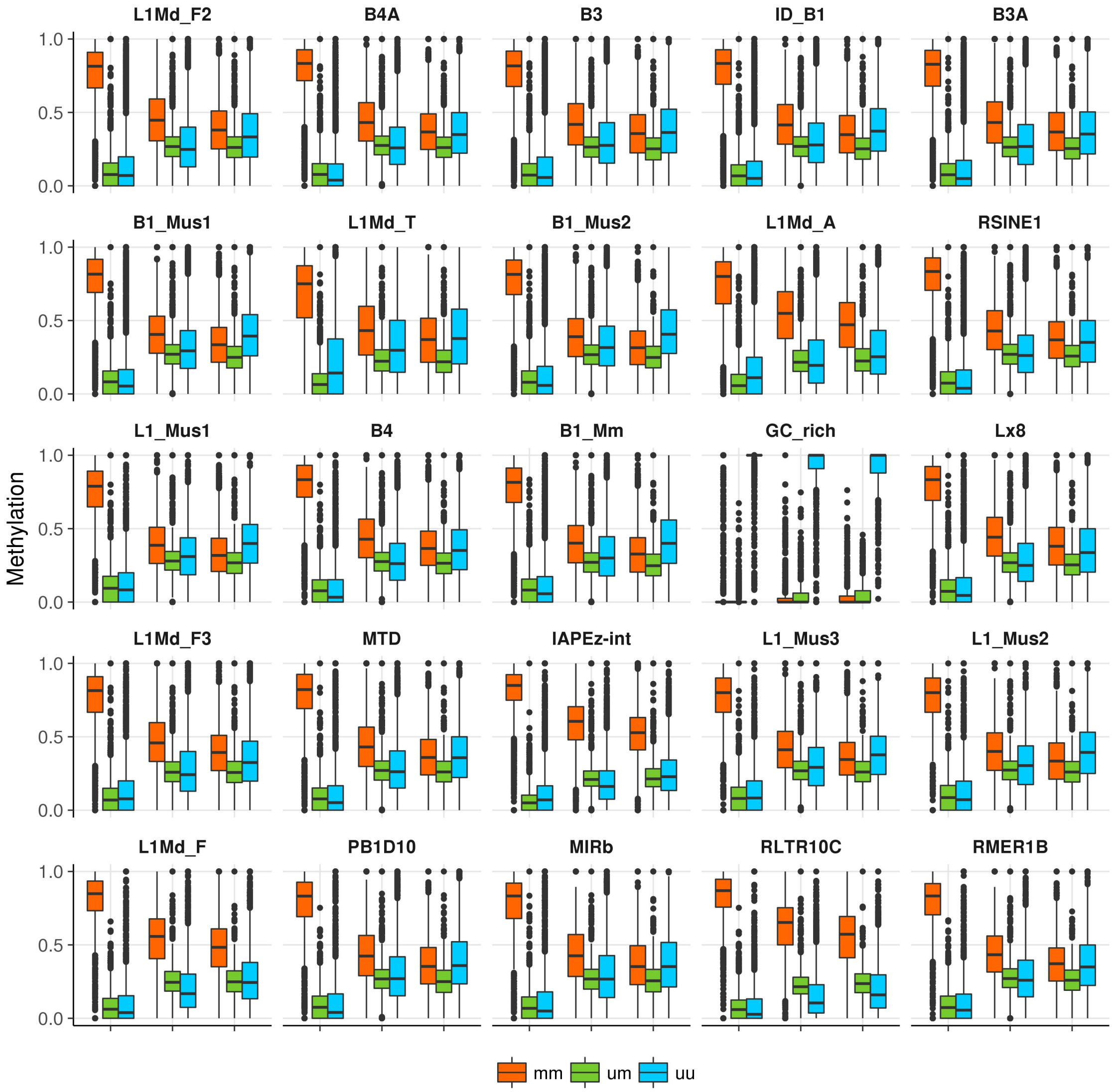


Figure 30: Methylation level at the 25 most frequent repetitive elements in our analysis for Tet TKO cells. Elements are presented in decreasing order, most frequent left top, least frequent right bottom. Annotation according to UCSC. y-axis = methylation frequency, x-axis = time in days (d0, d4, d7). Red = fully methylated CpGs (5mC/5mC), green = hemimethylated CpGs (5mC/C or C/5mC), blue = unmethylated CpGs (C/C).

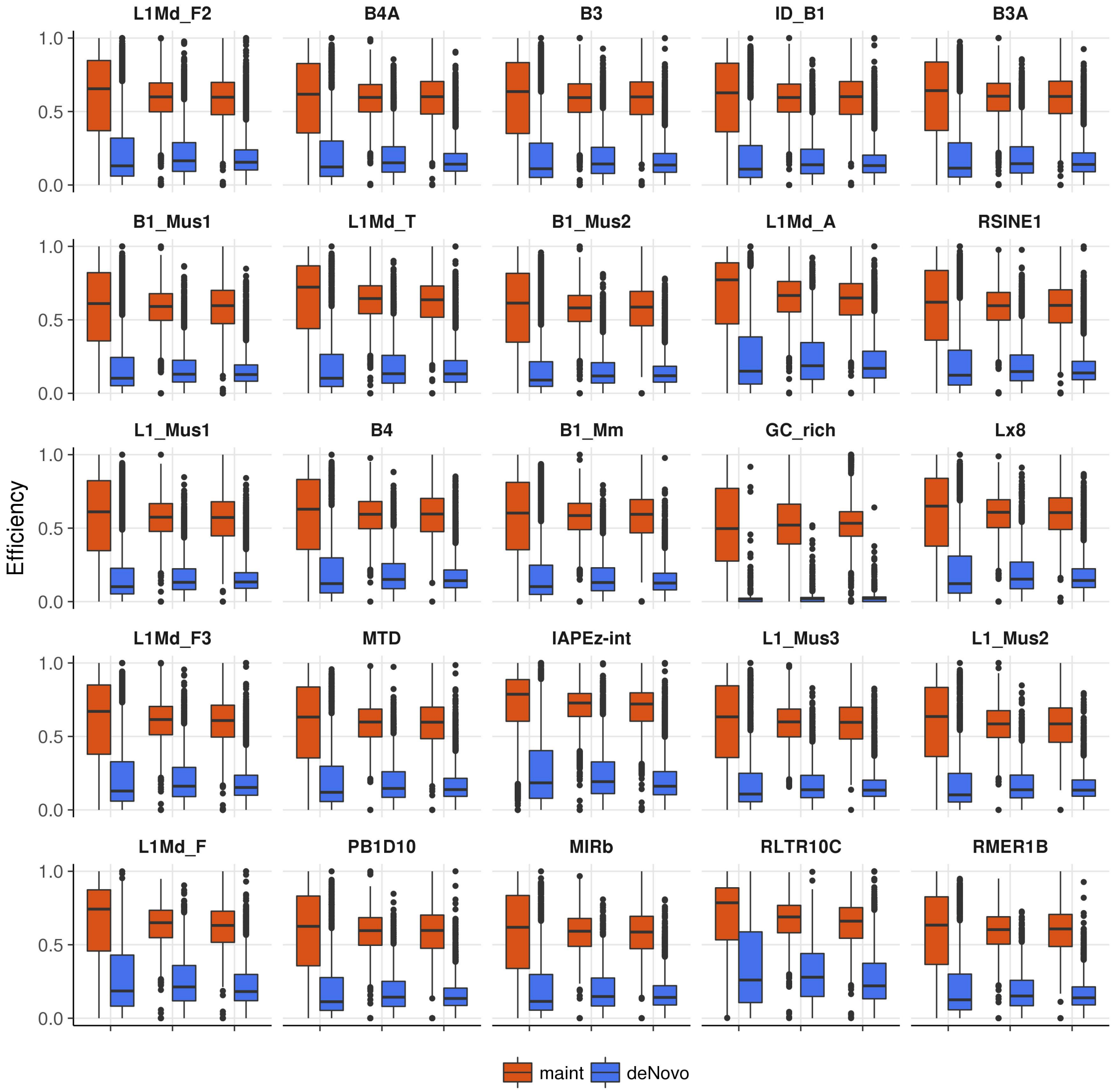


Figure 31: Efficiency profiles of the 25 most frequent repetitive elements in our analysis for Tet TKO cells. Elements are presented in decreasing order, most frequent left top, least frequent right bottom. Annotation according to UCSC. y-axis = efficiency; x-axis = time in days (d0, d4, d7), red = maintenance efficiency, blue = *de novo* efficiency.

#### 12 Relation of Efficiencies and DNA modifiers

In Figure 32 and 33 we compared the binding profiles (ChIP-Seq) of DNA modifiers from previous publications to maintenance, *de novo* and hydroxylation efficiencies estimated by our model (GSM659799, GSE57413, GSE100957) [13, 14, 15]. ChIP profiles for Dnmt3 iso-forms reveal a reduced binding around the TSS, which is in concordance with our model's prediction in reduced *de novo* efficiency. Interestingly, when comparing the ChIP profiles of Dnmt3s to transcriptome sequencing from Ficz *et al.* we observe distict profiles for expressed and non-expressed genes (Fig.: 32). In case of expressed genes, we observe a strong enrichment across the gene body and reduced binding around the TSS, in particular for Dnmt3a1, while non-expressed genes display the strongest binding precisely at the TSS.

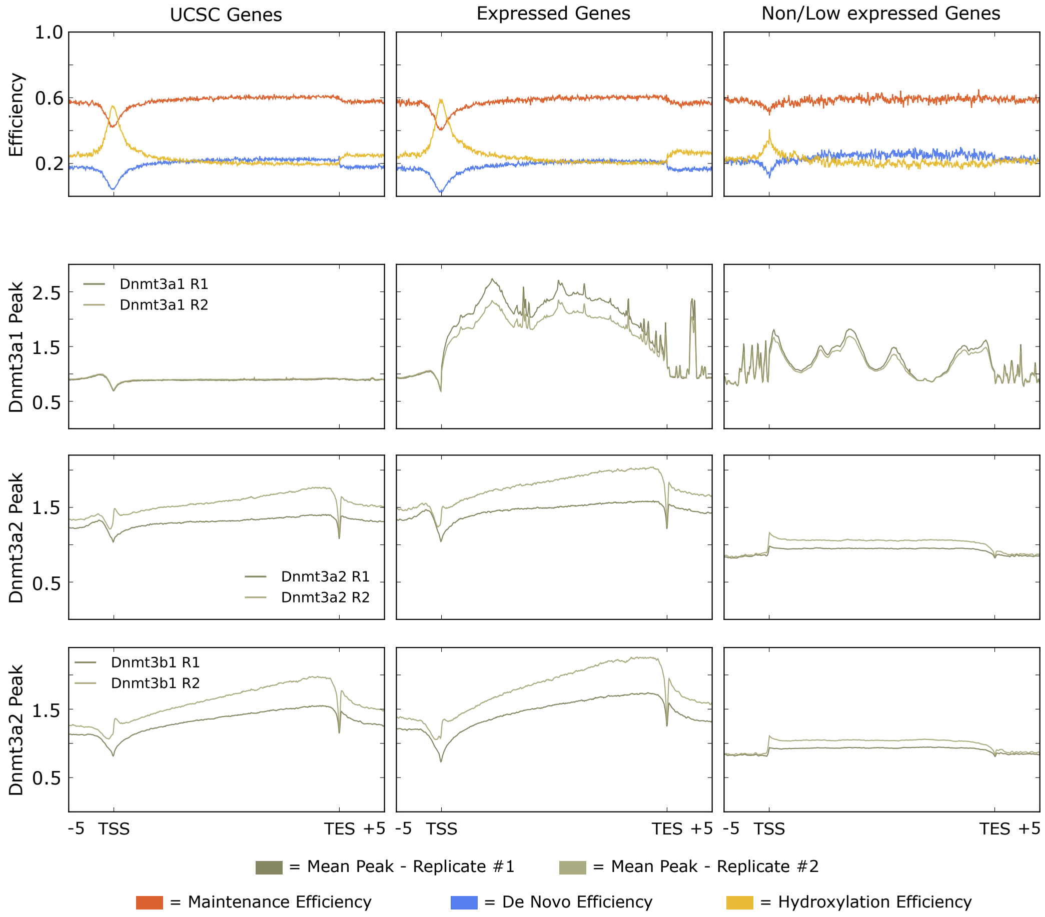


Figure 32: Estimated Efficiencies of Dnmts and Tets in form of maintenance methylation (red), *de novo* methylation (blue) and hydroxylation (yellow) as well as binding of Dnmt3a and 3b isoforms.

Additionally, we compared the efficiencies to ChIP binding profiles of Tet1 and Uhrf1, as one essential subunit of the maintenance machinery. Again, the ChIP profiles correspond nicely to our efficiencies of Dnmts and Tets (Fig.: 33). We observe, that Uhrf1 binds less frequent to TSS where we also observe a reduced maintenance, as well as *de novo* methylation efficiency. In contrast, Tet1 in displays a high enrichment at TSS matching the strongly increased hyroxylation efficiency observed by our model.

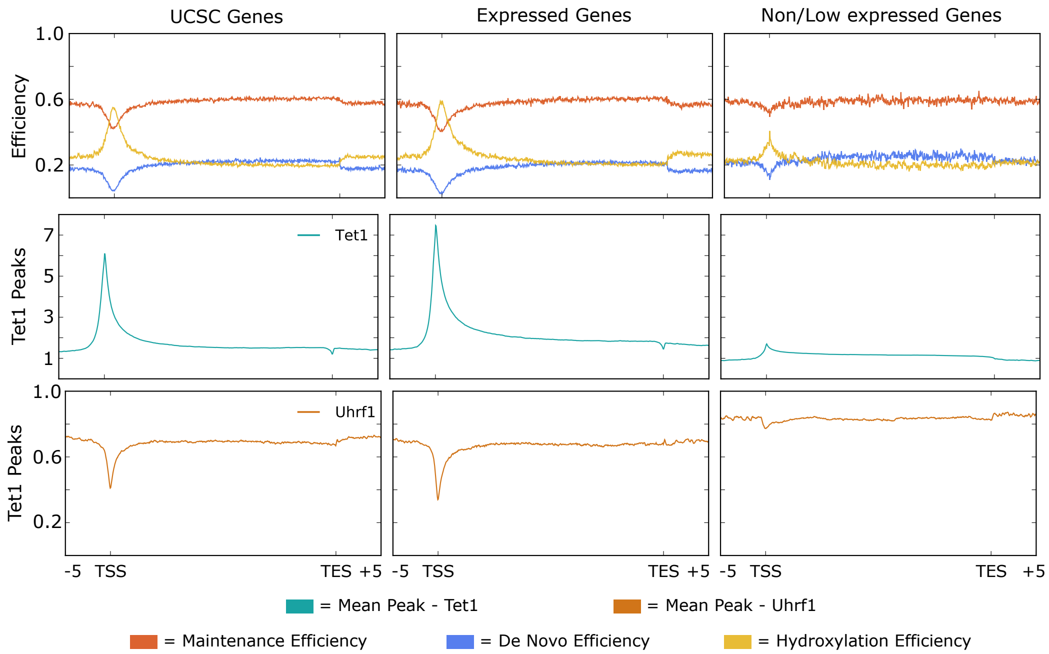


Figure 33: Estimated Efficiencies of Dnmts and Tets in form of maintenance methylation (red), *de novo* methylation (blue) and hydroxylation (yellow) as well as binding of Tet1 and Uhrf1.

### 13 Segmentation

We segmented the genome based on CpG methylation into HMRs, PMDs, LMRs and UMRs, using MethylSeekR (main manuscript Figure 9). Plotting available histone marks from ENCODE across the individual segment types reveals similar pattern as described before by Burger *et al.* [16].

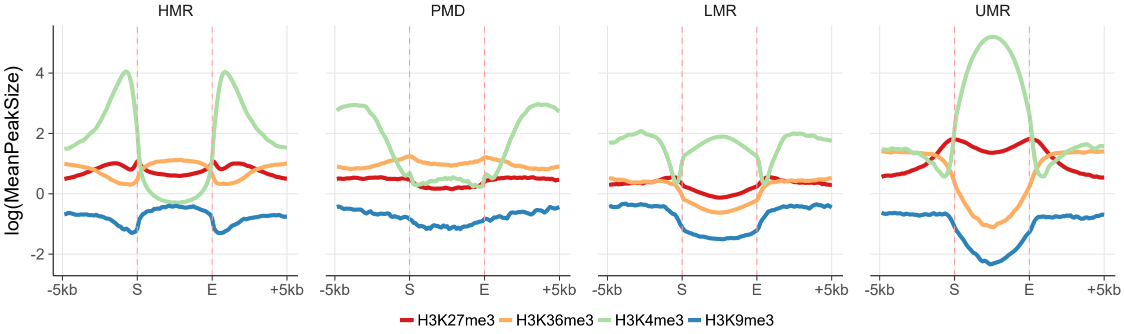


Figure 34: Profile of ENCODE histone modifications across HMRs, PMDs, LMRs and UMRs. Histone modifications are color coded; red dashed lines indicate start (S) and end (E) of a given segment.

Figure 35 shows how the clustered CpGs relate to the segmentation derived from the WGBS data. Expectedly, HMRs contain mainly CpGs from Cluster 1, 3 and 4, which display the highest methylation efficiency and consequently the highest methylation levels. A similar distribution can be observed for PMDs. However, CpGs of cluster 4, which exhibit high *de novo* efficiencies are notably under-represent compared to HMRs. Only a few measured CpGs can be assigned to LMRs, but the majority below to cluster two, defined by high hydroxylation efficiency. Finally, almost all CpGs can be assigned to UMRs

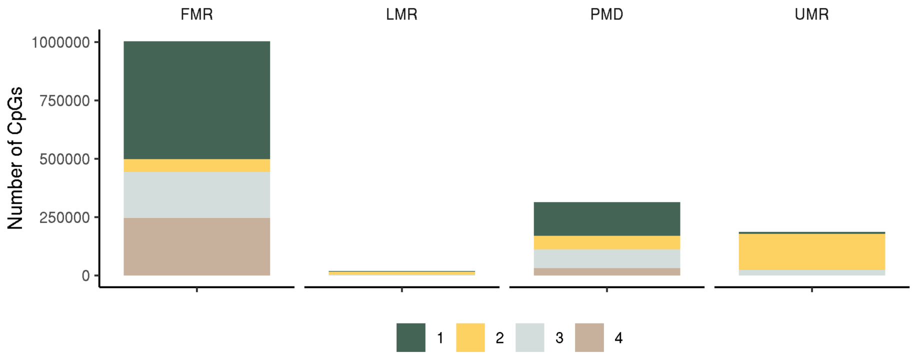


Figure 35: Distribution of clustered CpGs within the individual segments derived from partitioned WGBS data using MethylSeekR.

below to cluster 2.

Below, Figure 36 shows the average 5hmC level and distribution across the two DNA strands, estimated by our model. Under primed conditions, PMDs exhibit the highest level of 5hmC, followed by HMRs, LMRs and UMRs, respectively. Additionally, we observe that HMRs, PMDs and UMRs exhibit a transient increase in 5hmC after the transfer in 2i containing medium. In contrast, LMRs exhibit their highest amount of 5hmC under primed conditions and compared to the other segments types, display a relatively high amount of symmetrically hydroxylated CpGs (5hmC/5hmC, yellow).

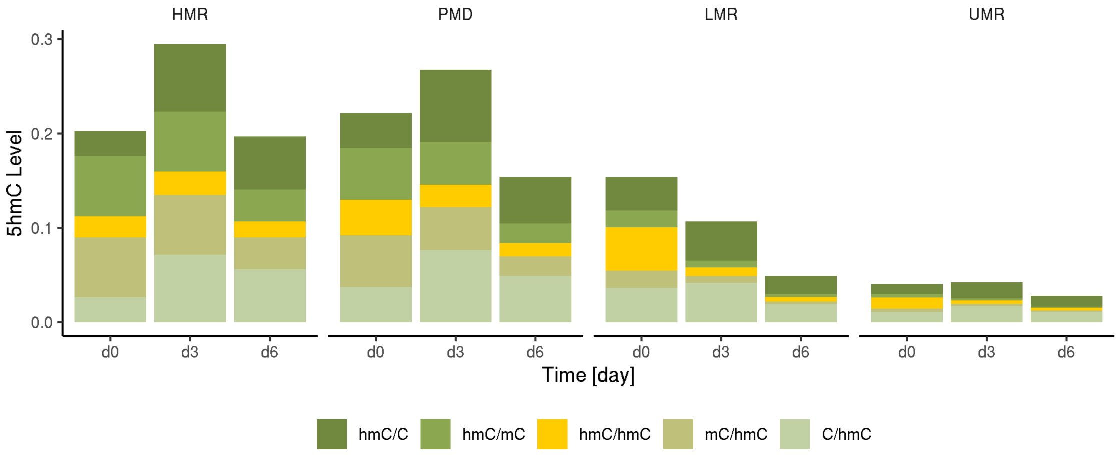


Figure 36: Average distribution of 5hmC across plus and minus strand for the individual segment classes.

### 14 Low Input Libraries

For the generation of Hairpin-RRBS libraries using 18 ng of genomic DNA, we performed three individual restriction reactions (R.AluI - NEB R0137S, R.HaeIII - NEB R0108S and R.HpyCH4V - NEBR0620S) in 1x CutSmart buffer with 6 ng of mouse ES cell DNA (72h 2i) each. 5 U of restriction enzyme were used in a total reaction volume of 10 µl. Reactions were incubated at 37 °C for 3 h. After heat inactivation at 80

°C for 20 min, reactions were pooled and 0.5 µl of CEGX spike-in control (1 pg/µl) was added. A-tailing was performed by adding 1 µl of 10X CutSmart buffer, 1 µl of 1 mM dATP, 2.4 µl of 5 U/µl Klenow exo- (NEB M0212S) and 5.1 µl ddH2O for a total reaction volume of 40 µl. Reaction was incubated at 37 °C for 30 min and heat inactivated at 75 °C for 20 min. For ligation of HP and SA, 0.5 µl of 100 µM HP, 0.5 µl of 100 µM SA, 5 µl of 10 mM ATP and 2 µl of 2000 U/µl T4 DNA Ligase (NEB M0202T) were added for a total volume of 48 µl, ligation reaction was incubated at 16 °C over night. Depletion of unbound HP using AMPure XP beads, enrichment of HP-DNA using Dynabeads, oxidation reaction and bisulfite conversion (both according to TECAN Methyl® oxBS Module manual), enrichment PCR and QC were done similarly to our protocol outlined in the methods section. Sequencing was carried out in a 2x 150 bp mode on the Illumina MiSeq using MiSeq Reagent Nano Kit v2 (300-cycles).

[18] Yudi Pawitan. *In all likelihood: statistical modelling and inference using likelihood*. Oxford University Press, 2001.

[19] Jonathan Kipling Knight. *Rotation points from motion capture data using a closed form solution*. PhD thesis, University of Colorado at Colorado Springs, 2008.
